# Long-read transcriptome sequencing of CLL and MDS patients uncovers molecular effects of *SF3B1* mutations

**DOI:** 10.1101/2024.01.26.576051

**Authors:** Alicja Pacholewska, Matthias Lienhard, Mirko Brüggemann, Heike Hänel, Lorina Bilalli, Anja Königs, Kerstin Becker, Karl Köhrer, Jesko Kaiser, Holger Gohlke, Norbert Gattermann, Michael Hallek, Carmen D. Herling, Julian König, Christina Grimm, Ralf Herwig, Kathi Zarnack, Michal R. Schweiger

**Affiliations:** Institute for Translational Epigenetics, Faculty of Medicine, University Hospital Cologne, University of Cologne, 50931 Cologne, Germany; Center for Molecular Medicine Cologne (CMMC), Faculty of Medicine and University Hospital Cologne, University of Cologne, 50931 Cologne, Germany; Department of Computational Biology, Max Planck Institute (MPI) for Molecular Genetics, 14195 Berlin, Germany; Buchmann Institute for Molecular Life Sciences and Institute of Molecular Biosciences, Goethe University Frankfurt, 60438 Frankfurt, Germany; Institute of Molecular Biology, 55128 Mainz, Germany; Genomics & Transcriptomics Laboratory, Biological and Medical Research Center, Heinrich-Heine-University, and West German Genome Center, 40225 Düsseldorf, Germany; Cologne Center for Genomics (CCG), Faculty of Medicine, University Hospital Cologne, University of Cologne, 50931 Cologne, Germany; Institute for Pharmaceutical and Medicinal Chemistry, Heinrich Heine University Düsseldorf, 40225 Düsseldorf, Germany; Institute of Bio- and Geosciences (IBG-4: Bioinformatics), Forschungszentrum Jülich, 52428 Jülich, Germany; Department of Haematology, Oncology and Clinical Immunology, University Hospital Düsseldorf, 40225 Düsseldorf, Germany; Department I of Internal Medicine, Center for Integrated Oncology Aachen-Bonn-Cologne-Düsseldorf, Faculty of Medicine, University Hospital Cologne, University of Cologne, 50937 Cologne, Germany; Medical Clinic and Policlinic 1, Hematology, Cellular Therapy, Hemostaseology, and Infectious Diseases, University of Leipzig Medical Center, 04103 Leipzig, Germany

**Keywords:** SF3B1, chronic lymphocytic leukemia, CLL, myelodysplastic syndrome, MDS, long-read sequencing, IsoSeq, differential splicing, iCLIP, molecular dynamics

## Abstract

**Background:** Mutations in splicing factor 3B subunit 1 (*SF3B1*) frequently occur in patients with chronic lymphocytic leukemia (CLL) and myelodysplastic syndromes (MDS). These mutations have a different effect on the disease prognosis with beneficial effect in MDS and worse prognosis in CLL patients. A full-length transcriptome approach can expand our knowledge on *SF3B1* mutation effects on RNA splicing and its contribution to patient survival and treatment options.

**Results:** We applied long-read transcriptome sequencing to 44 MDS and CLL patients with and without *SF3B1* mutations and found > 60% of novel isoforms. Splicing alterations were largely shared between cancer types and specifically affected the usage of introns and 3’ splice sites. Our data highlighted a constrained window at canonical 3’ splice sites in which dynamic splice site switches occurred in *SF3B1*-mutated patients. Using transcriptome-wide RNA binding maps and molecular dynamics simulations, we showed multimodal SF3B1 binding at 3’ splice sites and predicted reduced RNA binding at the second binding pocket of SF3B1^K700E^.

**Conclusions:** Our work presents the hitherto most complete long-read transcriptome sequencing study in CLL and MDS and provides a resource to study aberrant splicing in cancer. Moreover, we showed that different disease prognosis results most likely from the different cell types expanded during cancerogenesis rather than different mechanism of action of the mutated *SF3B1*. These results have important implications for understanding the role of *SF3B1* mutations in hematological malignancies and other related diseases.

**Highlights:**

- Long-read transcriptome sequencing data enables the identification of > 60% of novel isoforms in the transcriptomes of CLL and MDS patients and isogenic cell lines.
- *SF3B1* mutations trigger common splicing alterations upon *SF3B1* mutations across patient cohorts, most frequently decreased intron retention and increased alternative 3’ splice site usage.
- Mutation effect depends on alternative 3’ splice site and branch point positioning that coincide with bimodal SF3B1 binding at these sites
- Molecular dynamics simulations predict reduced binding of SF3B1^K700E^ to mRNA at the second binding pocket harboring the polypyrimidine tract.

## Background

Splicing is a fundamental step in eukaryotic gene expression in which non-coding introns are removed from pre-messenger RNA (pre-mRNA) transcripts and exons are joined to form mature mRNAs. This intricate process is often disrupted in cancer, either by mutations in spliceosomal genes or by other mechanisms that affect normal splicing function (1–5). In turn, this can lead to changes in the composition of expressed isoforms and the formation of new isoforms that alter the encoded proteins and can have far-reaching consequences for cellular function. One striking example of splicing alterations in cancer are mutations in the gene encoding the splicing factor 3B subunit 1 (*SF3B1*) that have divergent ramifications for treatment efficiency and prognosis (6,7). Somatic *SF3B1* mutations are frequently found in myelodysplastic syndrome (MDS, 20%), chronic lymphocytic leukemia (CLL, 15%), acute myeloid leukemia (3%), uveal melanoma (20%), cutaneous melanoma (4%), prostate cancer (1%) and in 2% of all breast, pancreatic and lung cancers (8).

In CLL patients, *SF3B1* mutations are typically subclonal and have been linked to disease progression and shorter survival (9,10). On the other hand, *SF3B1* mutations in MDS patients have been associated with specific disease phenotypes that show erythroid dysplasia with ring sideroblasts and ineffective erythropoiesis (11). Unlike in CLL patients, a positive effect of the *SF3B1* mutation on survival has been observed in almost all groups of MDS patients, except those with excess blasts, for whom no significant effect has been observed (12). However, so far there is no explanation for the divergent ramifications of *SF3B1* mutations in CLL and MDS pathology.

Within the cells, SF3B1 is part of the spliceosome and plays a critical role in 3’ splice site recognition. As a subunit of the U2 small nuclear ribonucleoprotein complex (snRNP), SF3B1 binds the branch point (BP) sequence upstream of the AG dinucleotide at intron–exon boundaries and thereby stabilizes spliceosome assembly at active 3’ splice sites. By influencing the recognition of different splice sites within a pre-mRNA transcript, SF3B1 can also modulate alternative splicing. The most common mutations in *SF3B1* accumulate in the HEAT domain at its C-terminus (Fig. 1a). The HEAT domain consists of 22 non-identical HEAT repeats that form the RNA binding interface for BP recognition. These mutations are predicted to impact the N-terminal domain involved in complex formation with other splicing factors (13). For example, hotspot mutations affect the interaction of SF3B1 with SURP and G-patch domain containing 1 (SUGP1), which is involved in BP recognition (14).

**Fig. 1.**
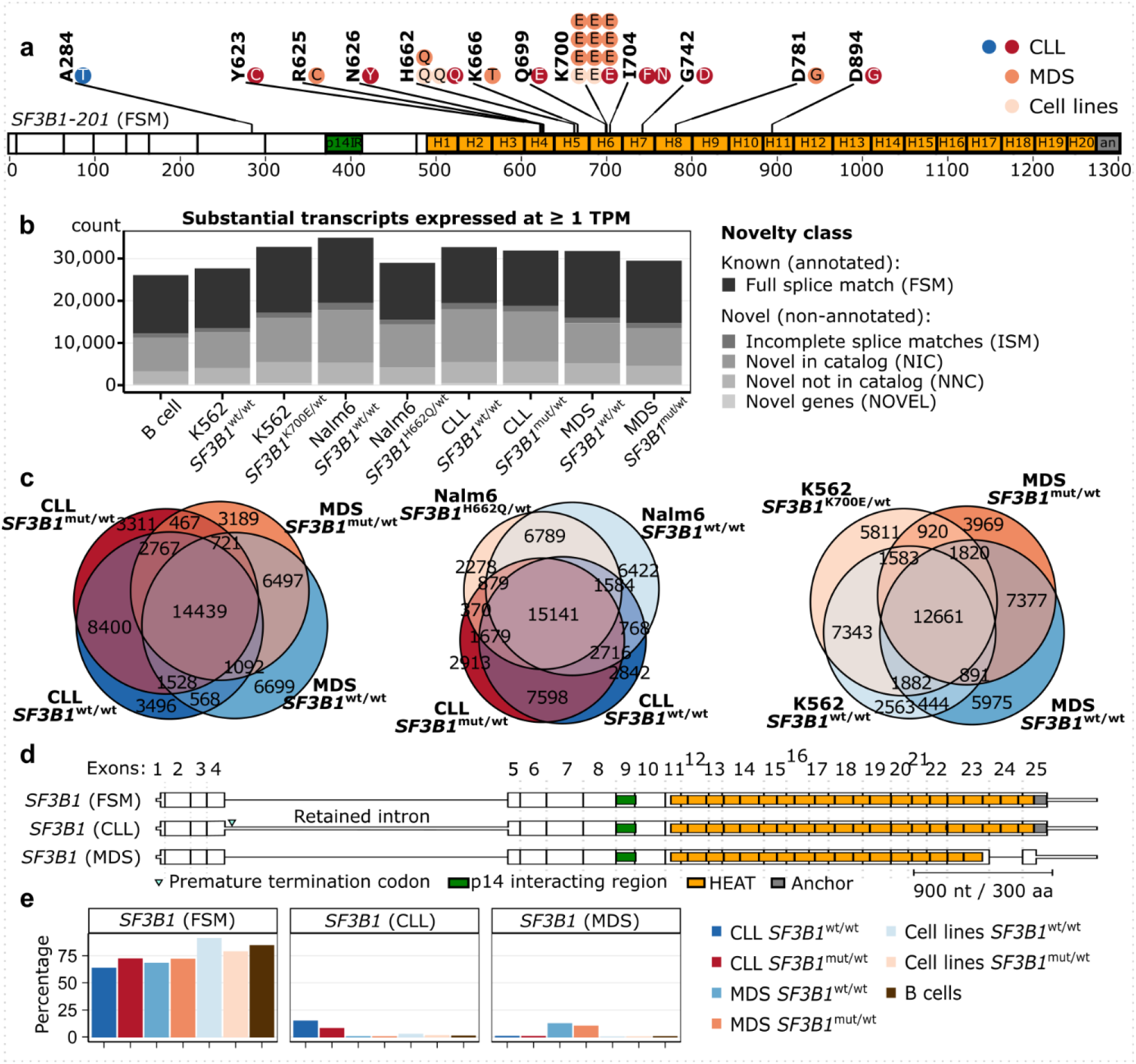
Long-read sequencing of CLL and MDS patient samples discovers novel isoforms. **a** Distribution of *SF3B1* mutations in CLL and MDS patient samples used for Iso-Seq: each dot represents a mutated sample. Note that the A284T mutation is outside the HEAT repeat domains, and thus was grouped as a wild-type sample, also according to further analysis described below. SF3B1 is shown as the major isoform expressed, with the full splice match to annotated isoform 201. **b** The number of substantial transcripts identified in each group and expressed at the level of at least 1 transcript per million (TPM) colored by the category of isoform novelty: full splice match (FSM), with incomplete splice matches (ISM), with combinations of annotated splice junctions (novel in catalog, NIC), with at least one novel splice site (novel not in catalog, NNC), or from novel genes (NOVEL) (21). **c** Venn diagrams showing the overlap between isoforms from b expressed at ≥ 1 TPM in each group. **d** *SF3B1* isoforms expressed at > 10% relative expression level. **e** Relative expression levels of *SF3B1* isoforms from d.

Previous studies based on short-read RNA sequencing (RNA-seq) have reported alternative 3’ splice site usage (3’AS) and intron retention (IR) as the most prominent splicing alterations in CLL and MDS patients with mutated *SF3B1* (3,15–18). The alternative 3’ splice sites (referred to as AG’) that were preferably used upon *SF3B1* mutation, are enriched at approximately 20 nucleotides (nt) upstream of the canonical splice sites (AG) (15–17,19). This strong positional constraint suggested that the mutations impacted SF3B1 binding and BP recognition upstream of the 3’ splice sites. Additionally, it was proposed that the mutation promotes the usage of otherwise inaccessible AG’ within the RNA secondary structure (18). Despite these hypotheses, the exact mechanism of the effect of mutations in *SF3B1* is still not resolved.

Here, we combined complementary data from MDS and CLL patients with isogenic cell lines to comprehensively characterize the effects of *SF3B1* mutation in cancer. Using long-read transcriptome sequencing, we identified > 60% novel transcripts in the samples that have not been covered in current gene annotations. Comparison of patients with a mutation in *SF3B1* (*SF3B1*^mut/wt^) to patients with wild-type *SF3B1* (*SF3B1*^wt/wt^) revealed that aberrant splicing was largely similar in both patient cohorts. We further examined the common changes in alternative splicing and isoform expression and their impact on the encoded protein functionalities. Integration with transcriptome-wide RNA binding maps and molecular dynamics simulations revealed that mutations in *SF3B1* obscured binding to the affected AG with increased usage of an alternative AG.

## Results

### Long-read RNA sequencing expands patient transcriptome landscapes in divergent biological contexts

To investigate the effect of *SF3B1* mutations on splicing, we characterized the transcriptomes of three datasets: CLL patients, MDS patients with ring sideroblasts, and isogenic cell lines with or without somatic *SF3B1* mutations. In brief, we collected CLL cells or whole-blood samples from 19 CLL and 25 MDS patients, including eight CLL patients and 14 MDS patients with mutations in the SF3B1 HEAT repeat domain (Fig. 1a). These were complemented by two isogenic leukemia cell line pairs (K562 and Nalm6), both with *SF3B1*^wt/wt^ and *SF3B1*^mut/wt^. K562 cells originated from a patient with chronic myeloid leukemia (CML) and Nalm6 cells from a patient with B-cell acute lymphoblastic leukemia (B-ALL). As controls, we further included B-cells from six healthy donors (Supplementary Table S1). To detect complete transcript isoforms, we performed long-read sequencing using Iso-Seq^®^ (Pacific Bioscience). We reached a mean sequencing depth of 582,135 full-length non-chimeric reads, cumulating in a total of 33,763,806 reads with an average length of 2,721 bp. Only 9% of reads were potentially affected by technology-specific technical artefacts (20) (Supplementary Fig. S1– S2).

In total, we identified 89,659 substantially expressed transcripts that contributed to at least 1% to a gene’s total expression and were covered by at least five full-length reads. Almost one third of these reads (28,261; 31.5%) were classified as full splice matches to annotated isoforms (FSM). Moreover, 58,168 (64.9%) represented novel isoforms that only partially overlapped with gene annotations, and 3,230 (3.6%) reads originated from non-annotated, novel genes (Supplementary Fig. S3). Even for transcripts expressed at ≥ 1 transcript per million (TPM), the novel isoforms consisted of more than half of all transcripts detected (Fig. 1b). A large fraction of isoforms was shared by the different patient cohorts and isogenic cell lines, with a larger overlap of expressed isoforms between *SF3B1*^mut/wt^ and *SF3B1*^wt/wt^ of the same dataset, than between datasets (Fig. 1c).

*SF3B1* gene has multiple isoforms annotated and was indeed expressed in several isoforms in both, samples with or without *SF3B1* mutations (Fig. 1d, Supplementary Fig. S4). Although the most frequently expressed *SF3B1*-FSM isoform (around 70% of *SF3B1* transcripts) fully corresponded to the annotated isoform, two shorter novel isoforms showed a disease-specific expression almost exclusively in either CLL or MDS patients These contributed approximately 10% each to the gene’s overall expression, irrespective of the *SF3B1* mutational status (Fig. 1e). The CLL-specific isoform (*SF3B1*-CLL) showed retention of the fourth intron which introduced a premature termination codon and likely targeted the isoform for nonsense-mediated mRNA decay (NMD). In the MDS-specific isoform (*SF3B1*-MDS), the penultimate exon was skipped and induced a frameshift that probably resulted in an NMD-resistant isoform that encoded for a C-terminally truncated protein devoid of HEAT repeats 18–20 and the anchor domain. In addition to the divergent splicing pattern, *SF3B1* also showed three times higher expression in CLL compared to MDS patients, whereas its levels were reduced in MDS patients when compared to B-cells from healthy donors (Supplementary Fig. S4).

Overall, our results demonstrated a high transcriptome diversity in the patient cohorts, which was dominated by a large number of novel transcripts.

### Patients and cell lines with *SF3B1* mutations show similar splicing defects

In order to investigate the transcriptome diversity at the splice-site level, we used the recently developed IsoTools (22) software to identify alternative splicing events (ASEs) in the transcripts expressed. IsoTools distinguishes exon skipping (ES), intron retention (IR), mutually exclusive exons (ME), and 5’ and 3’ alternative splice sites (5’AS and 3’AS) events, as well as alternative first and last junctions (AFJ and ALJ). Using a cut-off of ≥ 100 reads, ASEs were quantified as the proportion of reads supporting the ASE in relation to the sum of reads for all transcript isoforms, referred to as percent splice index (PSI). Across all samples, we discovered 82,028 ASEs in 9,333 genes, for which the less expressed ASE made up for at least 10% of the reads. For 75% of these events, at least one of the alternatives was not annotated (novel event) (Fig. 2a).

**Fig. 2.**
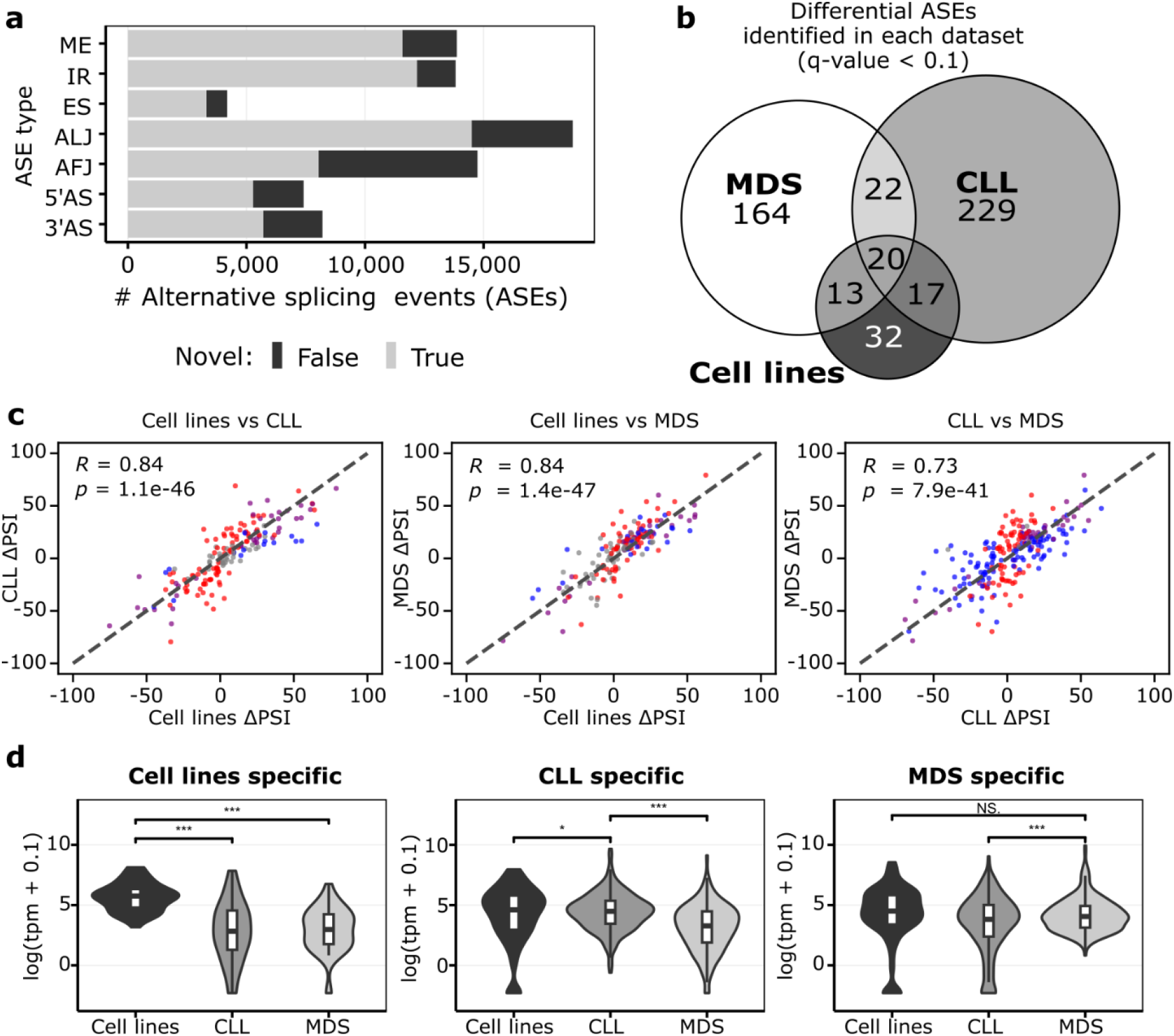
*SF3B1* mutation effect is independent of the biological background, but its manifestation depends on the transcriptomic profile. **a** The number of alternative splicing events identified with Iso-Seq separated by splicing event type in all groups investigated, differentiated by the novelty class. **b** Overlap between significantly altered alternative splicing events (ASEs) in samples with *SF3B1* mutation identified in the three datasets used (cell lines, CLL patients, or MDS patients). **c** Correlation among differences in isoform usage from b. Only events called significant in at least one of the two datasets per graph are shown. The colors of the dot correspond to significance reached only in one set (blue: X axis only; red: Y axis only; purple: both; grey: none). Pearson correlation coefficient (*R*) and associated p-value (*p*) are given. **d** Violin plots with boxplots show the distribution of expression values of the genes with dataset-specific ASEs from c. Significant differences are marked with *** for unpaired, two-tailed Student’s t-test p-value < 0.001, ** p-value < 0.01, * p-value < 0.05, N.S. – not significant with p-value ≥ 0.05.

Next, we used IsoTools (22) to detect significant differences in splicing associated with *SF3B1* mutations. Because *SF3B1* mutations have been reported to convey either beneficial (MDS) or disadvantageous (CLL) effects on patient survival (6,7), we first tested for differential splicing in *SF3B1*^mut/wt^ vs. *SF3B1*^wt/wt^ samples, separately in each dataset. We detected 82, 288, and 219 ASEs in the isogenic cell lines, CLL, and MDS patients, respectively (adjusted p-value [q-value] with false discovery rate, FDR (23) < 10%, Supplementary Table S2). Although we observed only moderate overlap of the identified events between the datasets (Fig. 2b), to our surprise, the correlation of the PSI changes for the ASEs identified was high (Fig. 2c), indicating a common mutational effect. We thus hypothesized that the disease-specific ASEs were due to variations in the transcriptomes of the expanded blood cell types in the two diseases, with the genes with higher expression levels in one disease reaching significant threshold only in this particular patient cohort. Indeed, the genes altered by the disease-specific ASE were generally expressed significantly higher in the corresponding group of patients (Fig. 2d, Supplementary Fig. S5) and out of the union of 531 genes with an ASE, 149 were relatively higher in MDS and 155 were higher in the CLL samples by at least 2-fold (FDR < 1%) based on the Iso-Seq data.

Our findings suggested that while *SF3B1* mutations introduced shared splicing effects in both CLL and MDS patients, the divergence in the disease outcome could be attributed to the differential transcriptomic profiles. Specifically, the mutation seemed to exert its most potent effects on genes that were already dominantly expressed in each disease, potentially magnifying the consequent functional impacts and underscoring the nuanced interplay between genetic mutations and disease-specific transcriptomic landscapes.

### *SF3B1* mutations affect alternative 3’ splice site usage and intron retention

Since the individual analyses for patients and isogenic cell lines indicated a common effect of *SF3B1* mutation, we combined all *SF3B1*^mut/wt^ and *SF3B1*^wt/wt^ samples to increase statistical power for an overall estimation of the *SF3B1* mutational impact. In total, we identified 775 differential splicing events in 530 different genes (Fig. 3a, Supplementary Table S3). As reported previously (3,15–18,24), the splicing changes upon *SF3B1* mutation were strongly enriched for 3’AS (326, 42%) and IR (213, 27%) events which together accounted for more than two thirds of the significant changes. The majority of IR events showed decreased IR (89%), whereas the majority of 3’AS events (78%) showed higher PSI values, corresponding to longer exons in *SF3B1*^mut/wt^ (Fig. 3b,c, Supplementary Fig. S6, Supplementary Table S3). Consistent with the common mutational effect, the regulated 3’AS events showed a uniform response across cohorts and allowed to cluster *SF3B1*^mut/wt^ and *SF3B1*^wt/wt^ samples (Fig. 3b).

**Fig. 3.**
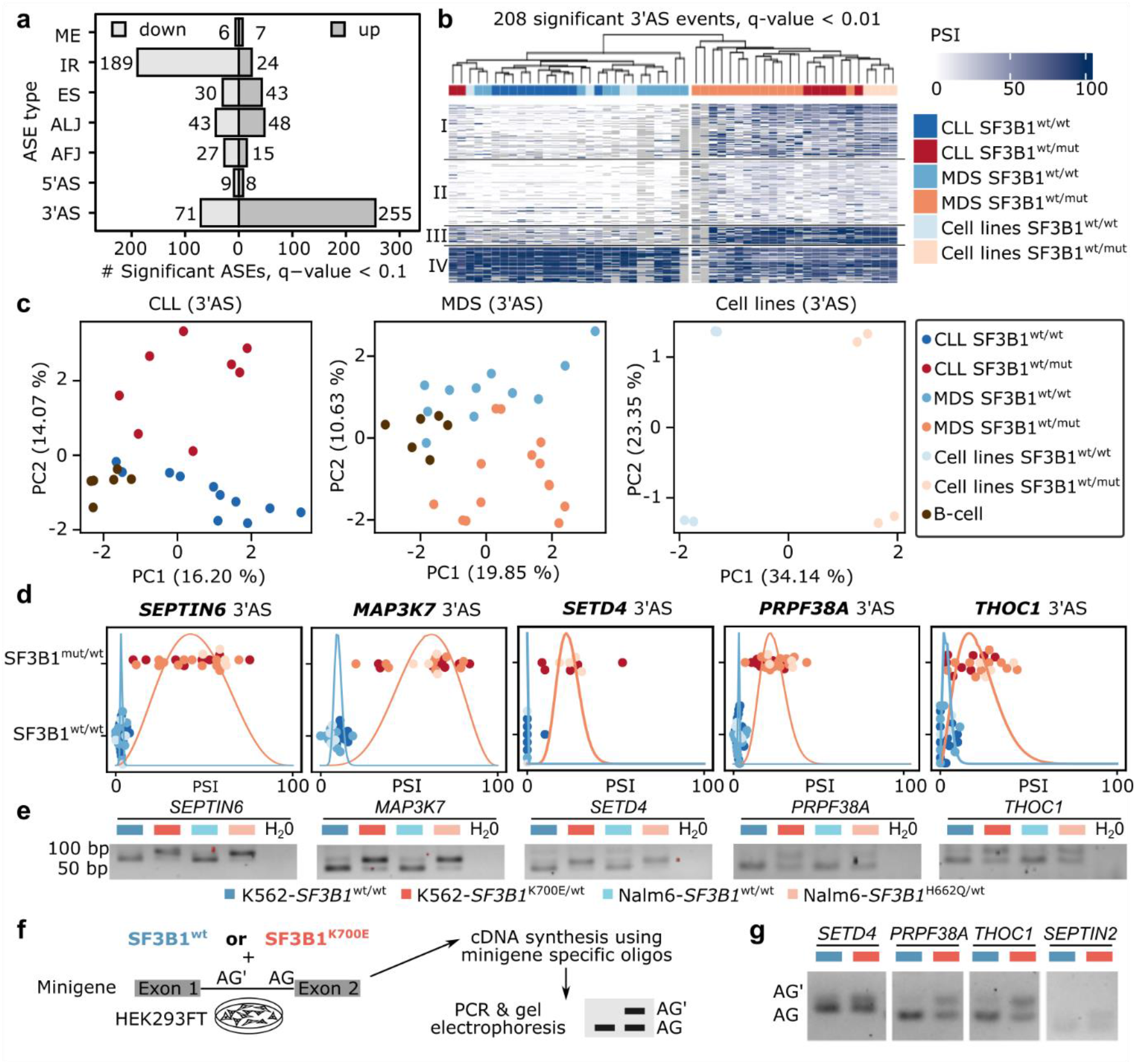
*SF3B1* mutation increases 3’ alternative splice sites usage and decreases intron retention. **a** Number of differential alternative splicing events with shorter or longer variant expression (percent spliced index, PSI) in *SF3B1*^mut/wt^ vs *SF3B1*^wt/wt^ samples. **b** Highly significantly altered (q-value < 0.01) 3’ alternative splice sites clearly separates samples by *SF3B1* mutations in leukemia cell lines as well as CLL and MDS patients based on the longer variant PSI values. **c** Principal component (PC) analysis, based on the isoform usage of 3’ alternative splice sites, clearly separates CLL and MDS patients, as well as cell lines, according to the *SF3B1* mutational status. **d** Swarm plots showing the distribution of the isoform usage (PSI) among groups with or without *SF3B1* mutation. **e** Validation of the differential splicing associated with *SF3B1* mutation with RT-PCR experiment in isogenic K562 and Nalm6 cell lines. **f–g** Minigene assays. HEK293T cells were co-transfected with minigenes and either *SF3B1*^wt^ or *SF3B1*^K700E^ for 48 h. RNA was extracted and used for amplification of splicing products with minigene-specific primers. The lower band in the agarose gel corresponds to the usage of the canonical AG, the upper band to the upstream AG’.

Upon closer inspection, we found that two CLL-*SF3B1*^mut/wt^ patient samples with mutations in the HEAT domain clustered with the *SF3B1*^wt/wt^ samples. One of these patients carried the rare *SF3B1* mutation Q699E, that had so far been reported only once, in a single patient with bladder urothelial carcinoma included in TCGA Pan-Cancer Atlas (according to cBioPortal accessed on 2023.10.16). Moreover, overexpression of the SF3B1-Q699H construct in HEK293FT cells did not lead to any aberrant splicing, suggesting that this mutation is weakly pathogenic (24). The second patient carried two mutations: D894G in HEAT repeat 11 (allele frequency (AF) = 51 %), and I704N (AF = 18%, Supplementary Table S1). For all other mutations, the position within the SF3B1 HEAT domain had no discernible influence on the clustering, indicating that the different mutations impair SF3B1 similarly (Fig. 3b). In fact, using general splicing information (PSI values), irrespective of regulation, we found that 3’AS events, but no other type of ASEs, clearly differentiated the samples based on the *SF3B1* mutational status in an unsupervised principal component analysis (Fig. 3c, Supplementary Fig. S7–S8), underlining the predominant effect of SF3B1 mutation on 3’AS events.

In addition to many new differential ASEs, previously published discoveries could be confirmed previously published discoveries, including e.g., five from 35 3’AS events (in *SEPTIN2*, *ERGIC3*, *RHNO1*, *FDPS*, and *SNRPN*) identified in CLL based on Nanopore sequencing (16), a 3’AS in *SEPTIN6* reported in MDS-*SF3B1*^mut/wt^ patients (25) (Supplementary Fig. S9), as well as thirteen 3’AS events (*BCL2L1*, *COASY*, *DPH5*, *DYNLL1*, *EI24*, *ERGIC3*, *MED6*, *METTL5*, *SERBP1*, *SKIV2L*, *TMEM14C*, *ZBED5*, and *ZDHHC16* that were consistently found in CLL, MDS, and uveal melanoma patients (26,27). Moreover, we confirmed 13 from 32 (28) and 8 from 11 genes (29) reported as aberrantly spliced in either MDS or CLL patients.

The LRTS data opened the possibility to assess the splicing alterations in the context of complete transcript isoforms. Generally, we found that the effect of the *SF3B1* mutation on splicing did not influence the choice of transcript start or end sites, nor the probability of other splicing events of the same gene. This means that the effect was local, and the resulting alternative transcript corresponded to the canonical transcript, except for the single alternative event. This is exemplified by the *SF3B1*^mut/wt^-induced inclusion of the poison cassette exon (PCE) in the *BRD9* gene that introduces a premature termination codon (PTC) (26) (Supplementary Fig. S10–S11). Of note, the full-length reads allowed us not only to confirm differential splicing of this PCE, but also to locate it to a specific isoform that has not been annotated yet (NIC class). This long-read-derived novel isoform otherwise resembled the canonical *BRD9* isoform, whereas the annotated PTC isoforms were presumed to also harbor an alternative first exon and additional splicing alterations. Such incomplete and incorrect isoform annotations are likely to cause problems in quantifying transcriptome changes, especially when using short-read sequencing data.

Notably, we found multiple splicing factors among the genes affected by *SF3B1* mutations, Overrepresentation analysis revealed 18 from the spliceosome pathway (KEGG: 03040, q-value 2.8e^-3^). This indicated a broad impact of *SF3B1* mutations on the general splicing machinery, which could potentially lead to secondary effects on splicing. Taking a closer look at the 208 highly significant 3’AS events (q-value. < 0.01), we identified four clusters based on PSI values detected in *SF3B1*^mut/wt^ and *SF3B1*^wt/wt^ (Fig. 3b, Supplementary Fig. S12). Interestingly, the cluster with low PSI values in *SF3B1*^mut/wt^ and no expression in *SF3B1*^wt/wt^ was enriched in spliceosome (q-value = 0.0106) and cell cycle genes (GO: 0007049 q-value = 2.8e^-4^, Supplementary Fig. S13).

To independently validate the detected splicing changes, we performed short-read RNA-seq on the isogenic cell line pairs as well as a subset of the 26 CLL patient samples and collected 398 publicly available MDS patient RNA-seq data (Supplementary Table S1, Supplementary Fig. S14). Using rMATS^21^ we observed a high and significant (p-values < 0.001) correlation of PSI values in the ASEs detected with IsoTools (22) and rMATS (30) on the same cell lines (Pearson correlation coefficient *R* = 0.840, p-value = 2.47e^-17^). The same held true for the samples from CLL and MDS patients (*R* = 0.794 with p-value = 2.95e^-36^ and *R* = 0.800 with p-value = 3.60e^-24^, respectively, Supplementary Fig. S15).

As an orthogonal approach, we employed semi-quantitative reverse transcription PCR (RT-PCR) to test 15 differential 3’AS events in the isogenic cell line pairs (Fig. 3e, Supplementary Fig. S16, Supplementary Table S4). From these 15 tested, 12 (80%) clearly showed an increase in alternatively spliced isoform expression in the *SF3B1*^mut/wt^ conditions. The strongest effects were observed for 3’AS events in *MAP3K7*, *SEPTIN6*, and *SETD4*, which showed an almost complete switch to the alternative splicing variant (Fig. 3e). We additionally performed splicing reporter assays using minigenes for six 3’AS events in HEK293T cells. Indeed, we observed differential 3’AS usage upon ectopic *SF3B1*^K700E^ expression in four out of six minigenes tested (*SETD4*, *PRPF38A*, *THOC1*, and *SEPTIN2*, Fig. 3f), supporting that the *SF3B1* mutation effects persist in an unrelated cell line. In the two remaining cases (*TPP2* and *BRCA1*) the usage of the upstream AG’ was already low in the validation assays using the K562 and Nalm6 cell line pairs (Supplementary Fig. S16).

Overall, these results supported a common effect of *SF3B1* mutations in different biological backgrounds and confirmed their predominant impact on alternative 3’ splice site usage and intron retention.

### Splicing alterations modulate protein function and stability

The capability of long-read sequencing to yield full-length transcripts provides a unique opportunity to annotate open reading frames (ORFs) and thereby predict potential impacts on the protein products. To assess the coding potential of the identified isoforms, we aligned the predicted ORFs to the pre-existing annotated coding sequences (CDS). The objective was to establish criteria that would reliably signify translation initiation and consequently, predict the coding potential of the transcripts.

Among the 28,261 known transcripts (FSM), we found that 73.4% had a matching reference CDS (Fig. 4a). In contrast, the majority of the 58,168 novel transcripts presented less evident coding potentials. Specifically, 20.1% lacked an ORF with coding potential, 13.2% initiated from an unannotated start codon, and 56.4% began at an annotated initiation site but deviated from the reference CDS due to alternative splicing. We also calculated the probability of nonsense-mediated decay for each isoform. Intriguingly, we observed that 35.1% of the expressed novel transcripts were likely to be targeted by NMD, compared to only 8.2% among the known transcripts (Fig. 4a).

**Fig. 4.**
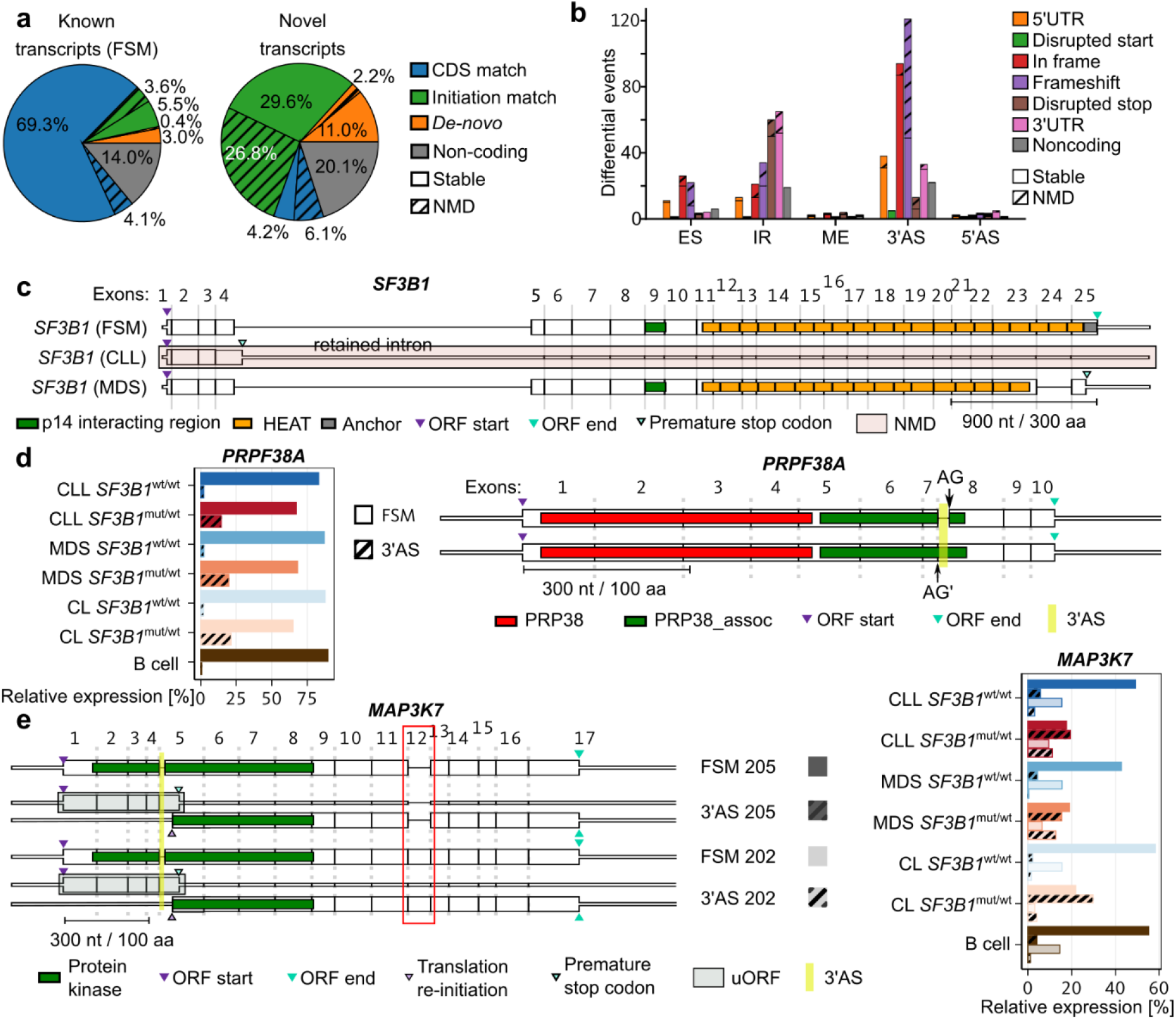
*SF3B1* mutation leads to protein disfunction. **a** Coding potential of the known and novel isoforms identified divided by CDS similarity to annotated isoforms and NMD prediction. **b** Effect of *SF3B1*^mut^-associated ASEs on the protein-coding potential. **c** *SF3B1* isoforms detected in this study with CLL- or MDS-specific isoforms. **d** The *PRPF38A* isoform expression levels (left) and structure with Pfam domains indicated (right). Highlighted in yellow is the *SF3B1*^mut^-associated 3’AS that may influence the protein function. **e** Schematic of major *MAP3K7* isoforms (left) with the protein kinase domain showed as green boxes. The *SF3B1*^mut^-associated 3’AS is highlighted in yellow and ORF start/end indicated by the triangles. Highlighted in light green are predicted uORFs. Red box highlights the additional exon 12 in the isoform 202, which is absent in the isoform 205. Expression of each isoform is shown on the right. The expression of isoforms with 3’AS event is shown as striped bars.

We next determined how *SF3B1* mutation-induced alternative splicing impacts the coding potential and the function of the proteins. To this end, we classified ASEs into categories based on their relative location and the impact on the coding sequence: 5’UTR, disrupted start codon, in-frame, frameshift, disrupted stop codon, and 3’UTR. Of the 326 events featuring 3’AS, the majority led to either frameshift modifications (121 events) or in-frame changes (94 events) in the CDS (Fig. 4b). As anticipated, frameshifts were predicted to lead to NMD for about half of the events, while in-frame modifications predominantly produced stable protein products. In total, we identified 274 ASEs predicted to yield stable alternative proteins.

Next, we examined more thoroughly the functional consequences of the two novel *SF3B1* isoforms, one predominantly expressed in CLL and one in MDS (Fig. 1d,e). In the *SF3B1*-CLL transcript, the fourth intron was retained which contained a PTC that shortened the CDS to 465 nt. Our prediction showed in a strong signal for NMD due to the presence of 19 downstream exon-exon junctions. In contrast, in the *SF3B1*-MDS transcript, the penultimate exon was skipped, through this a frameshift was introduced and, subsequently, a PTC. However, because this PTC was located within the last exon, the *SF3B1*-MDS transcript was unlikely to be targeted by NMD and resulted in a protein product missing its C-terminal section i.e., HEAT domains 18–20 and the terminal anchor domain (Fig. 4c).

To further investigate the functional outcomes of the altered proteins, we examined the impact on protein domain levels, by aligning Pfam (31) domains to the predicted protein sequences. For 57 of the ASEs, we found at least one Pfam domain that overlapped the divergent part of the protein sequence, indicating partially altered protein functions (Supplementary Table S3). For example, in *PRPF38A*, the 3’AS event overlapped the hydrophilic domain at the C terminus (Pfam: PF12871), which had seven additional amino acids in the *SF3B1*^mut/wt^ variant (Fig. 4d). This suggested that the alternative splicing event could influence the protein’s solubility or interactions with other molecules, thereby leading to potentially diverse functional consequences.

Interestingly, contrary to our initial predictions, we found evidence that some aberrantly spliced transcripts may escape NMD by translation re-initiation, if specific conditions are met. For example, in *MAP3K7*, a 3’AS event at the fourth splice junction led to a frameshift that created a PTC after 414 nt (Fig. 4e). This would ordinarily be a strong trigger for NMD due to the 11 downstream exon-exon junctions. Interestingly, however, the PTC in this transcript coincided with an in-frame start codon that aligned with the canonical CDS. Such overlapping start/stop codons are often seen in regulatory upstream open reading frames (uORFs, Supplementary Fig. S17). Another favorable signal for the translation re-initiation was a good Kozak sequence (Kozak score = +0.27) associated with this codon, and the fact that the prematurely terminated ORF was relatively short. This translation re-initiation would produce a protein with a truncated N-terminal region, impacting its protein kinase domain (Pfam: PF00069).

In summary, leveraging the advantages of long-read sequencing, we demonstrated a nuanced picture of how *SF3B1*^mut/wt^-induced splicing modifies both known and novel transcripts. Our data indicated that these splicing alterations also influence protein domain architecture and, in certain cases, escaped quality-control mechanisms like NMD. Intriguing examples such as *PRPF38A* and *MAP3K7* illustrated the variety of functional outcomes, from altered domain properties to the potential for translation re-initiation.

### *SF3B1* mutation effect depends on the distance and sequence context of alternative 3’ AS splice sites

When we plotted the fraction of differential 3’AS against the splice site differences we noticed, consistent with previous findings, that the alternative splice sites of differential 3’AS events were enriched within 12–21 nt upstream of the canonical splice site (AG) mainly used in the *SF3B1*^wt/wt^ samples (Fig. 5a) (15–19). Within this range, 30.8% of 3’AS were significantly differentially used in *SF3B1*^mut/wt^ compared to 1.8% outside this range. To validate the relevance of the AG’–AG distance for the *SF3B1* mutation effect, we constructed a minigene assay with a part of the *THOC1* transcript harboring the significantly affected 3’AS. The insert was then modified by replacing the 21 nt fragment between the AG’ and AG of *THOC1* with 45–50 nt long AG’–AG fragments from alternatively but non-differentially spliced introns (*PABCL1*, *USP1*, *ZNF124*, Fig. 5b, Supplementary Fig. S18). As a control, we mutated AG’ to GG’, to disrupt any alternative splicing (Fig. 5b). Then, we co-transfected HEK293T cells with the minigene constructs and either *SF3B1*^wt^ or *SF3B1*^K700E^ for 48 h and analyzed the expression by semiquantitative RT-PCR. As expected, with the original intronic sequence from *THOC1*, overexpression of *SF3B1*^K700E^ resulted in an increased usage of AG’, both for endogenous *THOC1* and the *THOC1* minigene (lanes 1–4, Fig. 5b). Of note, the insertion of the longer fragments from the *PABCLI* and *USP1* intronic sequences abolished the usage of the alternative splice site for the minigene constructs (lanes 7–8, 11–12, Fig. 5b). Moreover, the insertion of the longer intronic fragment of the *ZNF124* led to the sole usage of AG’ and increased intron retention (lanes 15–16, Fig. 5b). This could be explained by the weak canonical AG strength compared to the alternative AG’ (0.72 vs. 0.95, as calculated with SpliceRover (32), Fig. 5c). Importantly, no difference in splice site usage was observed when comparing *SF3B1*^K700E^ and *SF3B1*^wt^ overexpressing cells for each of the constructs tested. These results suggested that increasing the AG’–AG was sufficient to remove the 3’AS from SF3B1 regulation.

**Fig. 5.**
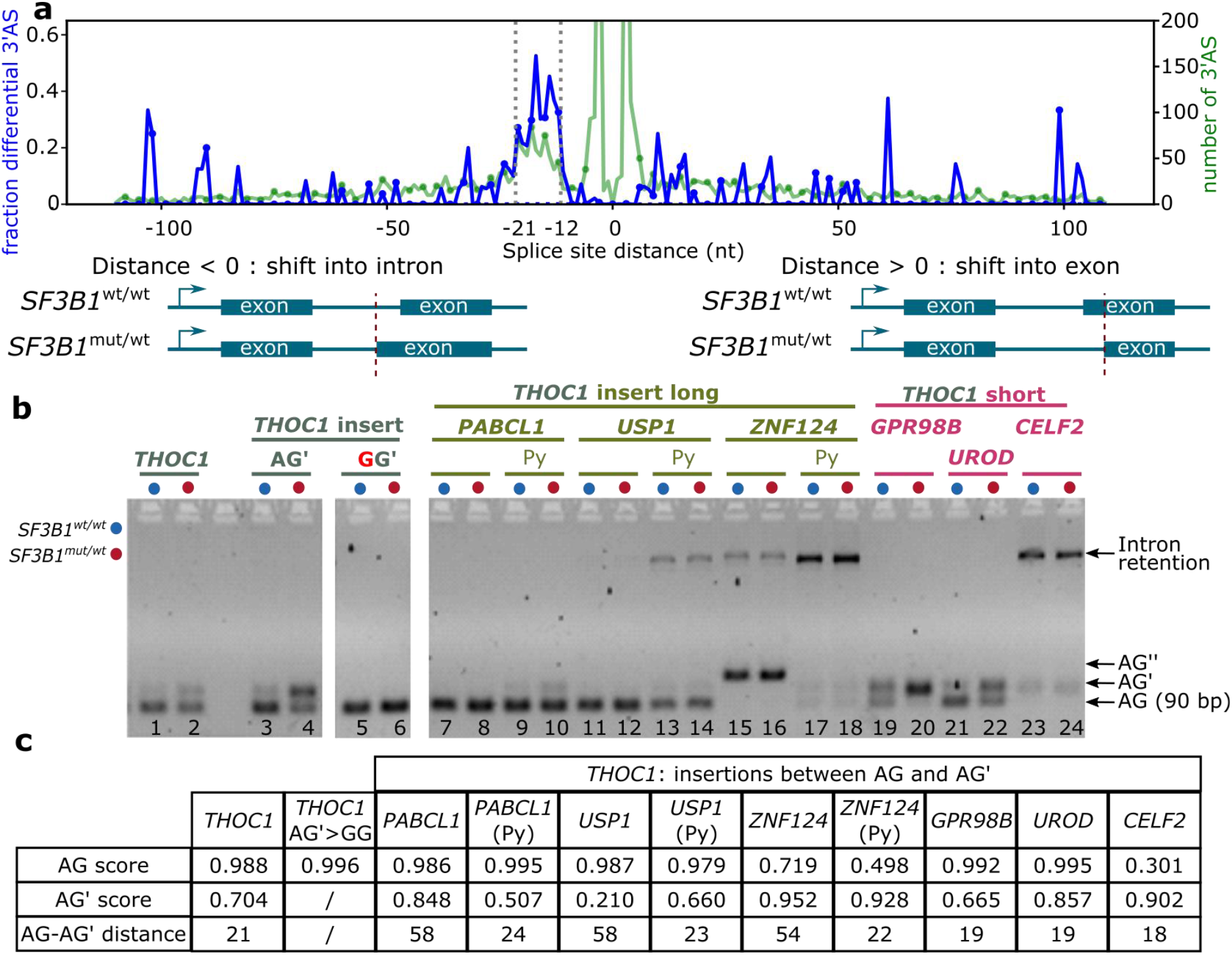
*SF3B1* mutations promote upstream alternative 3’ splicing sites and partially dependent on the sequence context. **a** 3’ alternative splice site distance distribution. Negative distances indicate an alternative was located upstream and positive values indicate alternative located downstream leading to a shorter exon. Blue line represents proportion and green total number of 3’alternative splicing events (3’AS). Dotted vertical lines indicate the enriched region of 12–21 nt upstream of the canonical AG. **b** Minigene assays with long (45–50 bases) AG–AG’ inserts, shortened inserts containing about 20 nt directly upstream the AG including the polypyrimidine (Py) tract, and short (15–20 nt) AG–AG’ inserts from non-differentially alternatively spliced 3’AS events. The chosen events without differential splicing detected with *SF3B1* mutation were from *PAPCL1*, *USP1* and *ZNF124* (AG’–AG distance > 50 nt) as well as *GPR98B*, *UROD* and *CELF2* (AG’–AG distance < 20 nt). **c** Table showing splice site strength for AG and AG’ calculated with SpliceRover (32).

Next, we asked if the AG’–AG distance was the main factor that influenced the 3’AS and created minigene assays with shorter, approx. 20nt versions of the previously used constructs with preserved polypyrimidine (Py) tracts. Indeed, for the *PABCL1*-Py construct the usage of AG’ was stronger upon overexpression of *SF3B1*^K700E^ compared to *SF3B1*^wt^ (lanes 9–10, Fig. 5b). This indicated that the AG’–AG distance rather than the specific sequence of this construct was determinant for the SF3B1 regulation. This was different when assessing the shortened versions of the other two constructs.

The *USP1*-Py construct was spliced mainly at the canonical AG and to a little extent at AG’, but also led to IR (lanes 13–14, Fig. 5b). The *ZNF124*-Py construct was only weakly spliced, but did not show any preference towards AG or AG’ and resulted in increased IR instead (lanes 17–18, Fig. 5b). Interestingly, in this instance again the canonical AG was weak (0.498, Fig. 5c). Neither the *USP1*-Py nor the *ZNF124*-Py constructs were differentially spliced when comparing *SF3B1*^wt^ and *SF3B1*^K700E^ overexpressing cells.

We therefore hypothesized that not only the distance, but also the sequence content between AG’ and AG plays a role for the alternative splicing in *SF3B1*^mut/wt^. To test this, we replaced the short AG’–AG region of the *THOC1* minigene with short AG’–AG fragments (12–21 nt) from three non-differentially spliced 3’AS events (in *GPR98B*, *UROD*, and *CELF2*). Interestingly, we noticed differential 3’AS usage between *SF3B1*^K700E^ and *SF3B1*^wt^ in two of the three constructs tested, indicating that the Py-tract of *GPR98* and *UROD* surrounded by *THOC1* sequences resulted in increased *THOC1* AG’ usage in *SF3B1*^mut^-expressing cells (lanes 19–22, Fig. 5b). Most likely *THOC1* specific sequences, such as the branch point region upstream of AG’ were responsible for the differential splicing between *SF3B1*^wt^ and *SF3B1*^K700E^ expressing cells.

In contrast, the *CELF2* insert led to a slight usage of AG’ and an increased IR in both, *SF3B1*^wt^ and *SF3B1*^K700E^ expressing cells (lanes 23–24, Fig. 5b). As for the weakly spliced *ZNF*-Py construct, also the *CELF2* construct had a weak canonical AG (0.301, Fig. 5c). This suggested that a strong AG is required for AG’ usage, in line with previously published work (24).

Our results confirmed that mutations in *SF3B1* primarily affected proximal AG’, with the distance itself did not seem to be the sole factor required for the usage of AG’. Moreover, we did not find any motif enriched at this position that could indicate a binding of another protein potentially disrupting or competing with SF3B1. We therefore speculated that SF3B1 binding at the sites with ASE may be altered in patients carrying *SF3B1* mutation.

### K700E mutation leads to destabilization of SF3B1-mRNA binding

To scrutinize the effect of the most common *SF3B1* mutation, K700E, we performed molecular dynamics (MD) simulations of 20 transcripts with a 3’AS within 50 nt distance, including 14 transcripts which were differentially spliced between *SF3B1*^mut/wt^ and *SF3B1*^wt/wt^ (*PRPF38A*, *RWDD4*, *SLC3A2*, *THOC1*, *TNPO3*, *SEPTIN2*, *IMMT*, *FDPS*, *INTS13*, *CDC27*, *PHKB*, *EIF4B*, *TRIP12*, and *LETMD1*) and 6 non-differentially spliced transcripts (*NAPG*, *SNX13*, *RIC8A*, *PIGB*, *PDCD4*, and *RMDN1*), see Methods for details. For each transcript, we performed four replicas of 200 ns long MD simulations of the mRNA with: i) the BP of the downstream AG (BP) bound to SF3B1^wt^; ii) the BP of the upstream AG (BP’) bound to SF3B1^wt^; iii) the BP bound to the SF3B1^K700E^ mutant; and iv) the BP’ bound to the SF3B1^K700E^ mutant. During the MD simulations, SF3B1 remained structurally invariant (Supplementary Fig. S19–S23). There were no differences in BP binding between all transcript–protein combinations, as analyzed by the number of contacts of all heavy atoms of the BP adenine to the heavy atoms of aromatic amino acids surrounding this nucleobase (first binding pocket): phenylalanine (F) 1153, and tyrosine (Y) 1157 (both located in SF3B1), or Y36 (located in PHF5A) Fig. 6a–c. In the SF3B1 wild-type, K700 interacts with the negatively charged oxygens of the mRNA backbone (second binding pocket, Fig. 6d). Importantly, the frequency of this interaction decreased in SF3B1^K700E^ due to repulsive interactions of like-charged atoms (Fig. 6e). The frequency of contacts between all heavy atoms of the SF3B1 residue 700 and all mRNA heavy atoms decreased too in the SF3B1^K700E^ (Fig. 6e). To investigate whether this decrease in interactions is associated with an increase in mRNA mobility, we computed the per-residue root mean square fluctuations (RMSF) of the mRNA nucleobases. Indeed, the mobility of nucleobases significantly increased for nucleobases in the 3’ direction after the K700 binding site for SF3B1^K700E^ compared to SF3B1^wt^, when the BP’ was bound to SF3B1 (Fig. 6f).

**Fig. 6.**
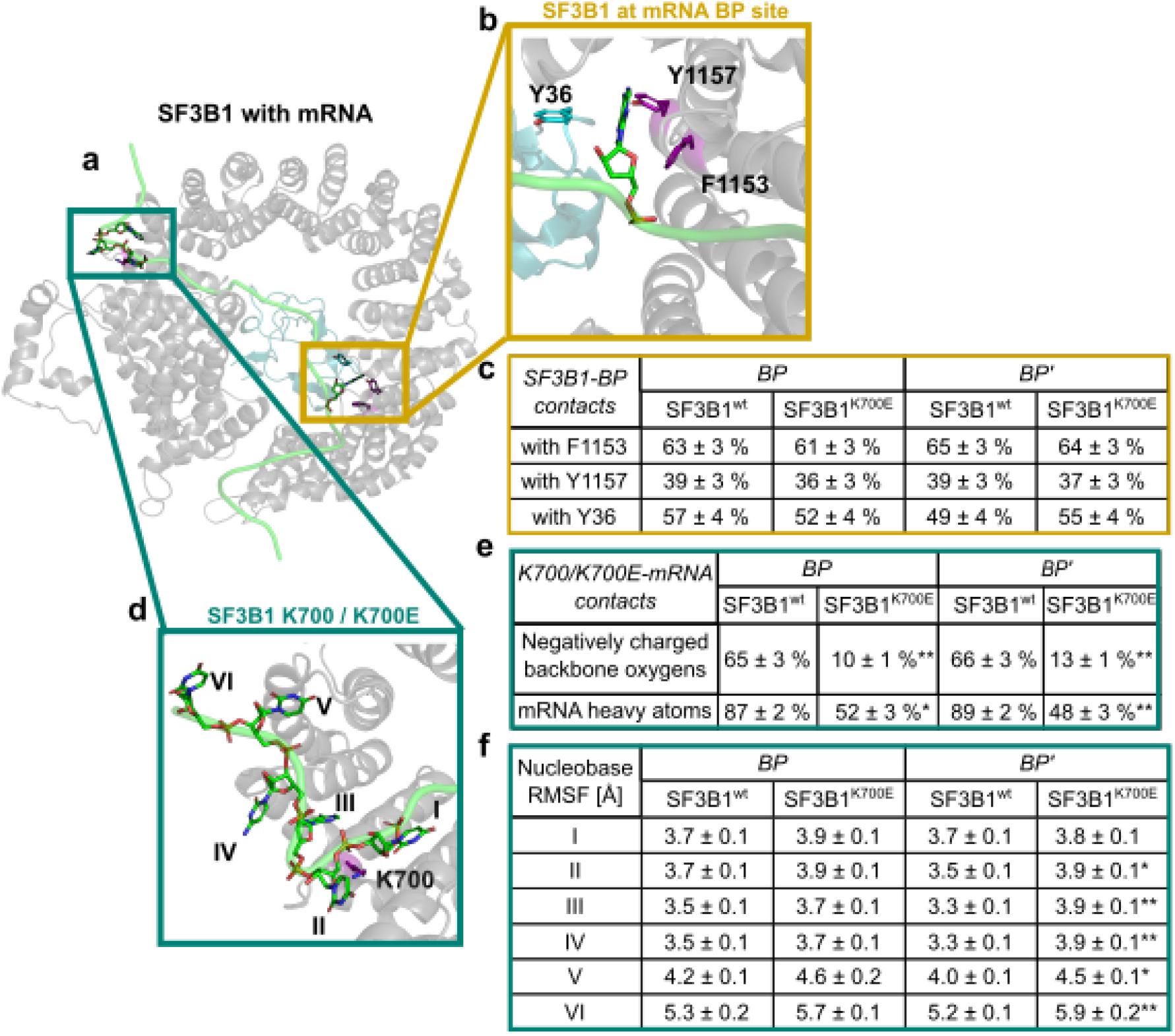
K700E mutation results in a destabilization of mRNA binding at the second pre-mRNA binding pocket around position K700. **a** simulated structure of SF3B1-mRNA: mRNA (green) bound to SF3B1 (grey) and the PHD finger-like domain-containing protein 5A (PHF5A, light blue). **b** Close-up of the BP recognition site. The branch point nucleobase (green) and aromatic amino acids surrounding the nucleobase (purple for SF3B1 and cyan for PHF5A) are shown. **c** Contact frequency of heavy atoms of the BP/BP’ nucleobase with heavy atoms of aromatic amino acids Y36, Y1157, and F1153 in the surroundings. **d** Close-up of the K700 binding site. K700 (purple) and surrounding nucleobases of the pre-mRNA (green) are shown exemplarily for the *NAPG* pre-mRNA. Nucleobases are numbered with Roman numbers such that K700 is positioned between nucleobase I and II in the starting structure of MD simulations. **e** Interaction frequency of the functional group in the side chain of K700/E700 to the negatively charged oxygen atoms in the pre-mRNA backbone and contact frequency of heavy atoms of K700/E700 with heavy atoms of the pre-mRNA. **f** RMSF of nucleobases as shown in panel d. Significant changes between SF3B1 and SF3B1^K700E^ are denoted by “*” for p-values < 0.05 and “**” for p-values < 0.01 (unpaired, two-tailed Student’s *t*-test).

Taken together, these results indicated that the K700E mutation did not lead to differences in the BP recognition. However, the mutation led to significant differences in SF3B1-mRNA contacts within the second mRNA-binding pocket at least for the 20 mRNAs tested with an upstream alternative AG within 50 nt.

### SF3B1 shows multimodal binding at 3’ splice sites

Our splicing analyses showed that mutations in *SF3B1* resulted in the activation of alternative splice sites at short distance from the canonical AG (Fig. 5a). To understand how SF3B1 recognizes these sites, we performed individual-nucleotide resolution UV crosslinking and immunoprecipitation (iCLIP) to map SF3B1 binding sites throughout the transcriptome (33). After UV crosslinking, we immunoprecipitated SF3B1 from K562-SF3B1^wt/wt^ and K562-SF3B1^K700E/wt^ cells, yielding together more than 100 million SF3B1 crosslink events (Supplementary Fig. S24, Supplementary Table S1 and S6). To facilitate direct comparisons, we randomly subsampled the sequencing reads to adjust the library size of the replicates (see Methods). Based on the merged iCLIP data we identified 96,852 SF3B1 reproducible binding sites with an optimal width of 5 nt (Supplementary Fig. S25–S27). The binding sites occurred in 8,127 genes, with the vast majority being protein-coding genes (93%). As expected, within the protein coding transcripts, SF3B1 mostly bound introns (94%, Fig. 7a,b).

**Fig. 7.**
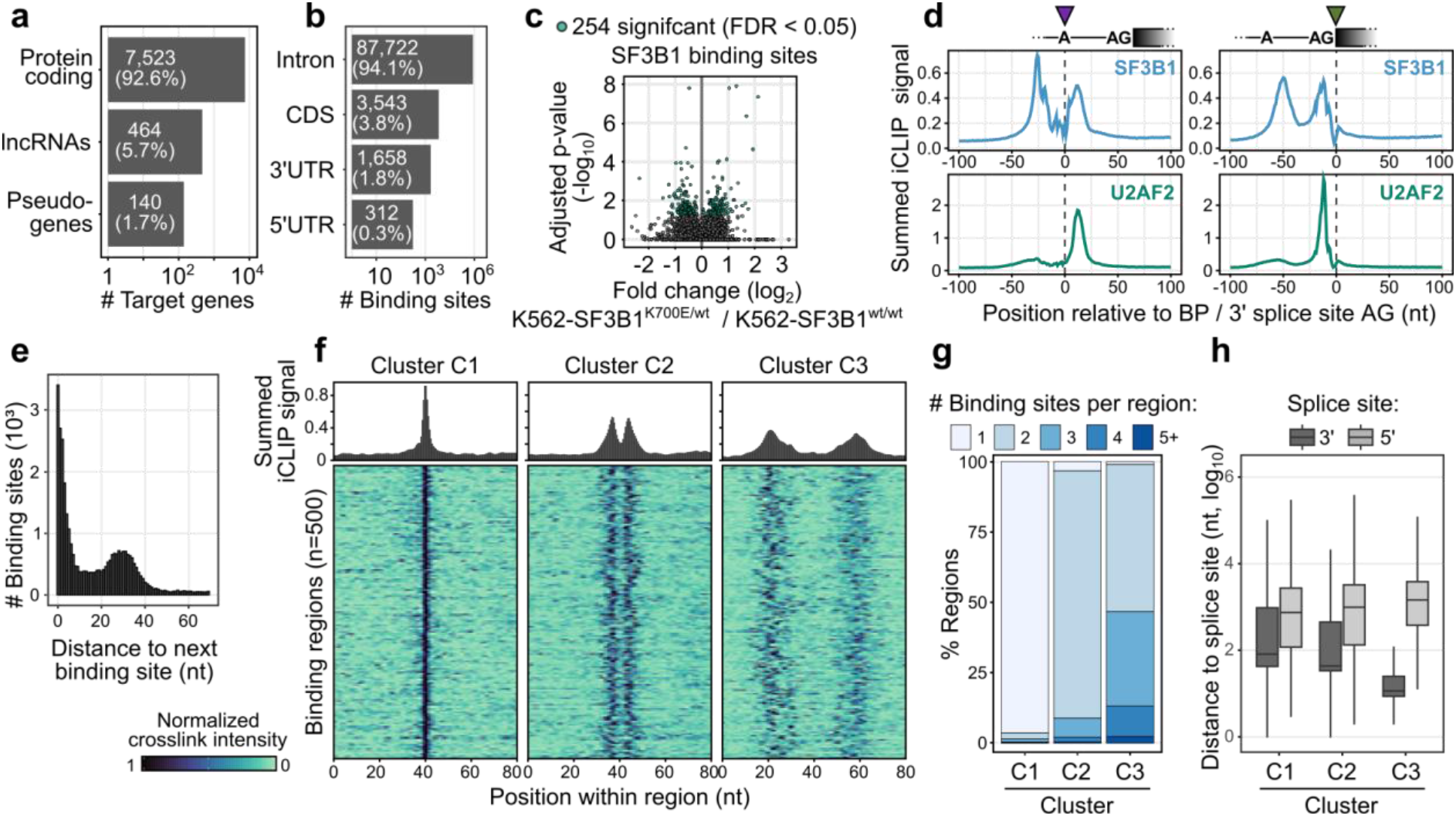
Multimodal SF3B1 binding. **a** Gene classes targeted by SF3B1 based on iCLIP. **b** Transcript regions of protein-coding genes targeted by the SF3B1 based on iCLIP. **c** differential SF3B1 binding sites in K562-SF3B1^K700E/wt^ vs K562-SF3B1^wt/wt^ **d** Meta-profiles of SF3B1 (top) and U2AF2 (bottom) binding centered at branch point adenosine (left) and 3’ splice site AG (right). **e** Distribution of distances between neighboring binding sites. **f** Density plot and heat map showing SF3B1 patterns in regions from three cluster types identified in Supplementary Fig. S29. **g** Distribution of the number of binding sites per region for each of the three clusters. **h** Distribution of the distance between SF3B1 binding region and closest splice site.

Since K562-SF3B1^K700E/wt^ cells expressed both wild-type and mutated SF3B1 protein and both variants were recognized by the anti-SF3B1 antibody that specifically binds to the SF3B1 N-terminus, we tested for differences in the SF3B1 binding between K562-SF3B1^K700E/wt^ and K562-SF3B1^wt/wt^. Consistent with a recent study (34), the K700E mutation did not generally impair RNA binding (Supplementary Fig. S27). Moreover, at the level of binding sites, we detected only minor differences between K562-SF3B1^K700E/wt^ and K562-SF3B1^wt/wt^ (Fig. 7c). Thus, although subtle differences may have been masked by the overlay of both protein variants in the heterozygous cells, these results suggested that the K700E mutation does not obviously change the global RNA binding behavior of SF3B1. However, local changes as observed with the molecular dynamics simulations might be too dynamic to be caught by the global iCLIP analysis

Next, we examined SF3B1 binding at 3’ splice sites. Using meta-profiles, we detected two prominent peaks of SF3B1 binding (Fig. 7d). The two peaks were centered at about 50 nt and 10 nt upstream of the 3’ splice site. Visual inspection indicated multiple SF3B1 binding sites within each peak (Supplementary Fig. S28). In line with SF3B1’s role in BP recognition and U2 snRNP recruitment (35), the distal peak clearly aligned with the predicted BP positions. The proximal peak, aligned to the 3’ splice site where it coincided with the Py-tract region bound by U2AF2 (36). To test this, we performed iCLIP experiments with U2AF2 in K562-SF3B1^wt/wt^ cells which confirmed that the proximal SF3B1 peak overlapped with U2AF2 binding (Fig. 7d). Together, these observations indicated that SF3B1 binds at both the BP and the Py-tract at the 3’ splice sites.

Consistent with the two peaks of SF3B1 binding at 3’ splice sites, we found that SF3B1 binding sites frequently occurred at distances of ∼30 nt to each other (Fig. 7e). To globally classify the SF3B1 binding pattern, we merged adjacent binding sites into equal-sized binding regions (80 nt, 56,224 regions) and performed unsupervised uniform manifold approximation and projection (UMAP) clustering. This yielded three distinct clusters: Cluster C1 harbored mostly isolated binding sites (35,907 regions), cluster C2 included two closely spaced binding sites (5,635 regions), whereas cluster C3 showed a more complex arrangement of 3–4 binding sites with wider spacing (13,847 regions) (Fig. 7f,g, Supplementary Fig. S29). The latter were located closest to 3’ splice sites (Fig. 7h), suggesting that multiple SF3B1 binding sites assemble into complex binding patterns at 3’ splice sites. We then investigated the differences in binding between K562-SF3B1^K700E/wt^ and K562-SF3B1^wt/wt^ solely in the C3 cluster. We noticed that K562-SF3B1^K700E/wt^ showed a slight decrease in the left peak, that was more distal to the canonical AG (Supplementary Fig. S30).

Altogether, our SF3B1 iCLIP data showed that SF3B1 adopted a multimodal mode of binding at 3’ splice sites, with two major peaks of SF3B1 binding that align with both BP and py-tract at 3’ splice sites. The peaks include multiple binding sites, which may reflect binding of several SF3B1 molecules or multiple contact points of the same molecule. The strong enrichment of this binding pattern specifically at 3’ splice sites suggested that the defined arrangement of binding sites is required for SF3B1’s function in splicing.

### SF3B1 alternates within a small window of alternative splice site distances and the K700E mutation leads to the use of the proximal upstream AG

We and others (14,17,24,37) found that 3’ splice sites are particularly sensitive to *SF3B1* mutations when they are directly preceded by an alternative 3’ splice site. To test how this relates to binding, we overlaid the iCLIP data with the splicing quantifications from the same isogenic cell lines (K562-SF3B1^wt/wt^ and K562-SF3B1^K700E/wt^). As shown above, the *SF3B1* mutations showed most prominent effects on alternative 3’ splice sites separated by 12–21 nt (Fig. 5a). Strikingly, at this distance, the upstream AG was at the edge of the proximal peak of SF3B1 binding (Fig. 8a). Moving further away, the effects subsided drastically as soon as the upstream AG emerged from the proximal peak. Moreover, within the 12–21 nt window, SF3B1^wt^ seemed to align to the downstream AG rather than the upstream AG, while it stayed more closely to the upstream AG at both shorter (< 12 nt) and longer distances (> 21) (Fig. 8b, Supplementary Fig. S30).

**Fig. 8.**
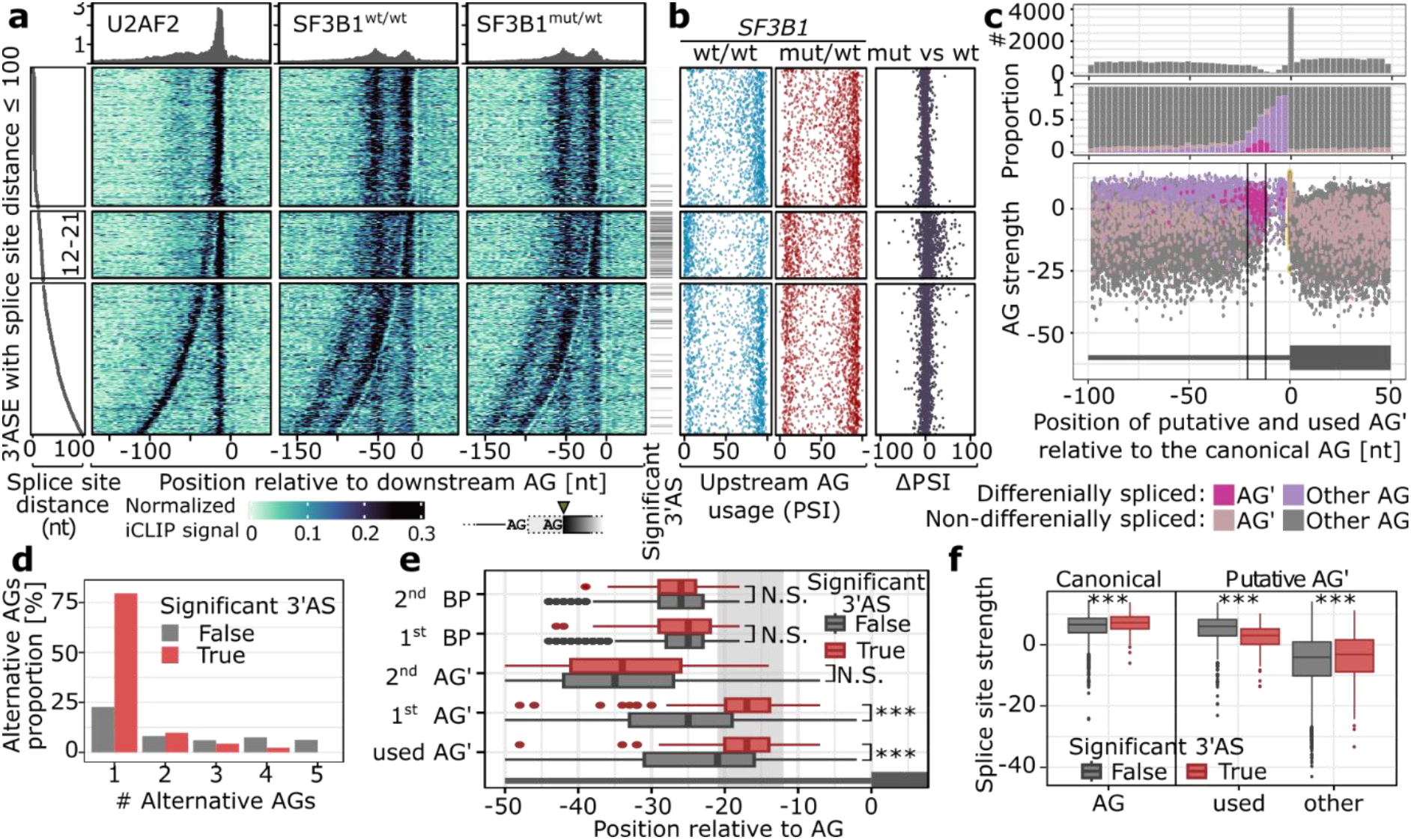
SF3B1 chooses alternative AG’ proximal to AG. **a** SF3B1 binding based on the iCLIP signal. On the left panel the splice site distance is shown, followed by the iCLIP signal aligned to the more downstream AG used. Significance of the differential 3’AS is shown as annotation bar on the right side of the iCLIP signal heatmap. **b** For every 3’AS from **a** the upstream AG PSI value is denoted for *SF3B1*^wt/wt^ (blue, left), *SF3B1*^mut/wt^ (red, center), and the difference between *SF3B1*^mut/wt^ and *SF3B1*^wt/wt^ (black, right) is shown. **c**–**d** Number of alternative AGs (AG’) among regions with multiple AG’s. AG’s within 6 nt from AG were removed to avoid NAGNAG acceptor sites (39). **e** Splice site strength of canonical and alternative AGs as calculated with MaxEnt score (38). **c** Distribution of AG occurrence (top), proportion of significantly alternatively used AG’ (middle) and AG’ scores among introns with significant (pink) or non-significant (violet) difference in usage between *SF3B1*^mut/wt^ and *SF3B1*^wt/wt^. **g** Distance of AG’s and BP’s from AG among significant (grey) and significant (red) events. For figures a–g Only 3’ASs with the following features were used for the analyses in c–g: i) placed chromosome scaffold; ii) classical AGs; iii) an intron became shorter in the mutant (the use of upstream alternative AG’); and iv) overlap with an iCLIP crosslink. The more used AG in *SF3B1*^wt/wt^ was set as canonical AG.

Interestingly, the 3’AS events within the critical window showed a distinct behavior already in wild-type K562-SF3B1^wt/wt^ cells. If the distance between 3’ splice sites was either < 12 nt or > 21 nt, splicing predominantly occurred at the upstream AG in the vast majority of cases, indicating that the spliceosome generally favored upstream AG usage. However, within the critical window, this pattern was inverted, such that upstream AG at these distances were generally outdone by the canonical downstream AG. This indicated that in presence of a downstream AG within 12–21 nt SF3B1^wt/wt^ binds predominantly the downstream AG. In line with this notion, we found that upstream AG were generally depleted from this region (Fig. 8c, top panel). Moreover, upstream AG still falling into this window showed basal usage already in wild-type cells in more than 50% of cases and were predominantly activated upon *SF3B1* mutation, which could be facilitated by the less stable binding of SF3B1 to the downstream AG as shown in the molecular dynamics model (Fig. 8c, middle panel).

Based on these observations, we hypothesized that diminished SF3B1^K700E/wt^ binding to downstream AG results in increased upstream AG usage within a critical window that may particularly impair splicing fidelity. Indeed, *SF3B1* mutation preferentially affected the first (nearest) upstream AG (Fig. 8d) which lay significantly closer to the 3’ splice site compared to non-differential 3’AS (Fig. 8e). In contrast, the BP of the differential spliced 3’AS events were neither moved closer to the 3’ splice site nor differed in their predicted strengths (Fig. 8e, Supplementary Fig. S31), indicating that the positioning of the upstream AG rather than the BP was the primary determinant for the observed effects. The differentially used upstream AG had slightly lower predicted splice site strengths (MaxEnt score (38)) than non-differential 3’AS helping SF3B1^wt/wt^ to bind to downstream AG. The downstream AGs were considerably stronger than unused upstream AGs (Fig. 8c, middle panel). Surprisingly, this was accompanied by higher splice site strengths of the canonical 3’ splice sites, indicating that a strong canonical 3’ splice site was required to support SF3B1^wt/wt^ binding within differentially spliced 3’AS events (Fig. 8f and Supplementary Fig. S32).

Taking together, we propose that SF3B1 binding often directly overlaps with alternative AG’ in a constrained window upstream of 3’ splice sites (> 20 % Fig. 5d), thereby using downstream 3’ splice site in wild-type conditions. In the presence of SF3B1 mutations, SF3B1 cannot properly bind the Py-tract of the downstream canonical AG when a strong upstream AG’ lies within a short distance, leading to increased alternative splicing with the use of an alternative branch point (Fig. 9). This is partly explained by changes in the second binding pocket of SF3B1, as predicted by molecular dynamics simulations, that destabilize the SF3B1-mRNA interaction. Thus, although mutated SF3B1 still binds to the BP in this scenario, the changed SF3B1 protein structure impairs its binding to the downstream AG, resulting in widespread splicing defects.

**Fig. 9.**
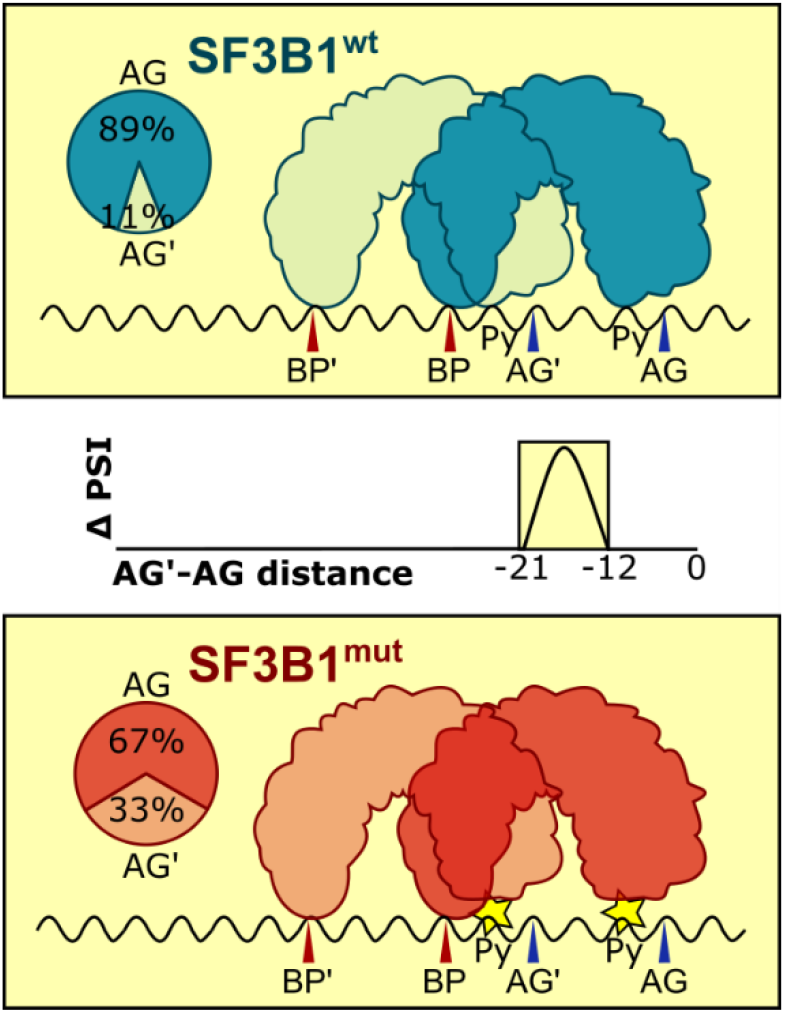
The model of SF3B1 effect on splicing. Within a short AG’–AG distance, the SF3B1^mut^ preferably uses the first (upstream) AG (AG’) compared to the SF3B1^wt^. This process often requires a strong downstream canonical AG. The switch between AG’ and AG may be explained by changes in the second SF3B1 mRNA binding pocket that contains the most frequent K700E mutation that lower the number of contacts between mRNA and E700 of SF3B1, resulting in increased mobility of the mRNA (indicated by the star). The percentages for upstream (AG’) and downstream (AG) AGs show the median PSI values calculated for all events with -12 nt ≤ AG’–AG distance ≥ -21 nt. BP – branch point, Py – polypyrimidine tract.

## Discussion

Alternative splicing plays a critical role in generating transcriptome diversity, and its dysregulation has been linked to various diseases, including cancer. Yet, complex transcriptome studies are challenging to perform with classical RNA-seq due to the frequent ambiguity in mapping short reads and the difficulties in identifying novel isoforms (40–42). Of note, our capacity to study transcriptomes and alternative splicing events has expanded with the recent advancements in long-read sequencing technologies. Nonetheless, up to now only two studies applied long-read Oxford Nanopore sequencing to analyze splice site alterations mediated by mutated *SF3B1* in CLL (bulk) (16) and MDS (single-cell) (43), both based on three patients only. For example, using Nanopore technology, 35 3’AS events were reported between *SF3B1*^K700E/wt^ and *SF3B1*^wt/wt^ CLL samples. However, this study used a low number of replicates and showed high differences in read coverage with a median read length below 1 kb, indicating that many events may have been missed due to quality issues (16).

Here, we used long-read Iso-Seq (PacBio) sequencing of 44 patients to investigate the impact of *SF3B1* mutations on alternative splicing. Our long-read sequencing analysis revealed a wide variety of transcripts, with more than two thirds of yet unannotated, or even over 3,000 transcripts from novel genes. This highlighted the importance of long-read sequencing for comprehensive transcriptome profiling, particularly for detecting novel splice variants and alternative splicing events. Our results supported previous findings that *SF3B1* mutations specifically alter the usage of 3’ alternative splice sites and intron retention (17,24,37). Importantly, using a comprehensive setup of two cohorts of CLL and MDS patients, complemented by two isogenic cell line pairs we were able to substantially expand the catalog of differentially spliced 3’AS to a total of 326 3’AS events in 266 genes (Supplementary Table S3). We observed similar effects in both patient cohorts and the cell lines studied, indicating a common effect of *SF3B1* mutation on splicing. The clinical differences and prognosis of the CLL and MDS patients with *SF3B1* mutations most likely depend on the specific gene expression profiles and thus their different relevance of splicing alterations.

Our results revealed that *SF3B1* mutations affect genes involved in the major mRNA splicing pathway, indicating a broader impact on the splicing machinery. This may trigger secondary effects on splicing, potentially leading to altered transcriptomes and disease phenotypes. In particular and in concordance with previous studies, we observed an enrichment of the 3’AS events 12–21 nt upstream of the canonical 3’ splice sites (17,24,37). This region is typically depleted of alternative AG dinucleotides. Why AGs are depleted in the critical region remains unclear. One possibility is that they are removed by purifying selection and might have an evolutionary advantage. However, a subset of introns still contains AGs within this critical region leading to alternative 3’AS. Why AGs are present in the critical region remains unclear. Interesting in this regard is that we and others (24) observed that in these cases, the canonical AG is often stronger than at other 3’ splice sites, indicating that a strong 3’ splice site may be required to tolerate an upstream AG within the critical window.

Our minigene assays did not confirm that these 3’AS events depend predominantly on the distance between canonical and alternative AGs. Coinciding with the critical range of AG’– AG distances of 12–21 nt, we observed a bimodal binding of SF3B1 at both, BP and Py-tract, whereby the BP often coincided or was near to the upstream AG’. Thereby, SF3B1 binding may shield the upstream AG from recognition during splicing Although, we did not detect obvious global changes in SF3B1^K700E^ binding, which might be due to a temporary effect in lower binding affinity of the SF3B1^K700E^ that might not be caught by iCLIP analyses. However, our MD simulations supported our model and indicated that the number of contacts between the mRNA and residue 700 of SF3B1 resulted in increased mobility of the mRNA at both, canonical and alternative 3’ splice sites in SF3B1^K700E^.

We confirmed with minigene assays that SF3B1^mut^ uses an alternative BP that leads to 3’AS usage in SF3B1^mut/wt^ cells (24,37). However, transcriptome-wide studies revealed that about one third of all human exons have multiple branch points (44,45), which argues against the hypothesis that mutated SF3B1 always prefers an alternative BP. Consistently, we did not observe any strong alterations in the binding of SF3B1^K700E/wt^ to mRNAs, although a slight increase of the second peak at the Py-tract directly upstream of the 3’ splice site was observed (Supplementary Fig. S33). Although we cannot exclude that we partially co-precipitated U2AF2, we and others did not find obvious changes in U2AF2 binding to SF3B1 in immunoprecipitations (37,46). Furthermore, U2AF2 binds to the Py-tract only during early stages of the splicing process and is released during transition to the activated B complex (47). In contrast, SF3B1 is assembled into the spliceosome a bit later, and subsequently replaces U2AF2 in the activated B complex and binds to the Py-tract with its binding pocket consisting of HEAT domains 3–7 that harbor most of the mutational hotspots (48). Therefore, the downstream peak observed in our SF3B1 iCLIP experiments most likely corresponds to SF3B1 binding.

The differential splicing observed in *SF3B1*^mut/wt^ may result from a weakened binding of SF3B1^mut^-containing spliceosomes. There might be an additional effect of decreased binding of DDX46/PRP5, a kinase involved in proof reading of the pre-mRNA branch site (49–52). Other splicing proteins that have been shown to bind less to SF3B1^mut^, are DDX42 (51), DHX15 (53) and SUGP1 (14,54). Interestingly, DDX42 and DDX46 have been shown to sequentially occupy the RNA binding pocket consisting of HEAT repeats 3–7 during early steps of the splicing process (52,53,55).

Besides gaining additional insight into the SF3B1 splicing mechanism, we also explored the splicing and expression alterations identified through the IsoSeq sequencing approach in CLL and MDS. Besides identification of large numbers of new differentially spliced genes, we were able to specifically map the toxic exon of *BRD9* to its isoform and predict its amino acid composition. We also identified *SF3B1* isoforms specifically more present in MDS or CLL, albeit their expression level was at approximately 10%. Further functional analyses will show the impact of these altered SF3B1 proteins and if they influence MDS and CLL pathomechanisms. Another example, which is frequently reported to be differentially spliced within *SF3B1* mutated cancers is *MAP3K7*. We were able to show that mutations in *SF3B1* led to reduced expression of longer isoforms and increased expression of isoforms with shortened protein kinase domain, likely impacting its function. Thus, with this data at hand it is possible to not only identify splicing events but to also map them to their cognate isoform and thus to provide information on the resulting protein composition. This is a fundamental information for understanding splicing data and to gain insight into pathomechanisms underlying CLL and MDS.

## Conclusions

Our study provides new insights into the mechanism by which *SF3B1* mutations affect splicing regulation, and the potential consequences on protein function. Our findings highlight the importance of long-read sequencing for investigating differential alternative splicing usage and splicing factor function. These results have important implications for understanding the role of *SF3B1* mutations in hematological malignancies and other diseases, and suggest new approaches for developing targeted therapies for these conditions.

## Methods

### Cell lines

The isogenic cell line pairs, K562-SF3B1^K700E/wt^ and it’s parental K562-SF3B1^wt/wt^ (RRID:CVCL_0004), as well as Nalm6-SF3B1^H662Q/wt^ and its parental Nalm6-SF3B1^wt/wt^ were obtained from Horizon Discovery (HD181-012, HD115-110). Since homozygous *SF3B1* mutations were reported to be lethal (56), we used heterozygous cell lines. The K700E mutation is the most frequent *SF3B1* mutation reported in CLL and MDS, and H662Q mutations is also frequently reported (4,6,9). The *SF3B1* mutated cell lines were described previously (24). HEK293-FT (RRID: CVCL_6911) was purchased from Thermo Fisher Scientific (#R70007). Cell line authenticity was tested by short tandem repeat (STR) profiling at the Cologne Center for Genomics, Cologne, Germany with 13 markers: *CSF1PO*, *D3S1358*, *D5S818*, *D7S820*, *D8S1179*, *D13S317*, *D16S539*, *D18S51*, *D19S433*, *D21S11*, *THO1*, *TPOX*, *vWA* and *AMELOGENIN* for sex determination. Subsequently, STR results were analyzed with the Cellosaurus STR similarity search tool CLASTR 1.4.4 and compared to the Cellosaurus data set 46.0 (algorithm: Tanabe, mode: non-empty markers, *AMELOGENIN* not included) run on 20230724. HEK293-FT and the K562 cell line pair were correctly assigned and the Nalm6 cell line pair matched 100 % to the Nalm6 derived cell lines Nalm6/H (RRID:CVCL-B7AM) and Nalm6/HDR (RRID:CVCL_B7AN) (57), compared to 83 or 81 % to Nalm6 (RRID:CVCL_0092). Moreover, the Nalm6 cell line pair differed at two marker loci (Supplementary Table S6). K562 cells were cultivated in Iscove Modified Dulbecco Media (IMDM) supplemented with 2 mM L-glutamine (Sigma-Aldrich), Nalm6 cells in RPMI-1640 (Thermo Fisher Scientific, #21875034) and HEK293-FT in DMEM, high Glucose, GlutaMAX™ (Thermo Fisher Scientific, 10569010). All media were supplemented with 10% fetal bovine serum (SIGMA-Aldrich) and 100 U/ml penicillin/streptomycin (Sigma-Aldrich). Cells were cultivated at 37°C with 5% CO_2_ and 95% humidity. Cell lines were tested negative for *Mycoplasma* using the Mycoalert Plus Mycoplasma detection kit (Lonza) according to the manufacturer’s protocol.

### Patient samples

CLL and B-cell samples were obtained from the CLL-Biobank Cologne. *IGHV* mutational status was determined, as previously described (58). Peripheral blood B cells were isolated via negative selection using RosetteSep immunodensity-based cell separation (Stemcell Technologies, Vancouver, BC, Canada). The purity of CLL/ B cells was analyzed by flow cytometry and revealed that ≥ 90% cells co-expressed CD5/CD19.

Specimens from MDS with ring sideroblast (MDS-RS) patients were obtained from the MDS Biobank of the University Clinic Düsseldorf. Either RNA or cells were obtained from the Biobank. If cells were obtained, RNA was isolated using the Nucleospin RNA kit (Macerey Nagel). RNA quality was accessed by RNA ScreenTape analysis (Agilent) or a Bioanalyzer (Agilent)

Clinical information on the patients is summarized in Supplementary Table S1.

Informed consent was obtained from all patients and the study was approved by the local ethics committee.

### Plasmids

pCMV-3Tag-1A-SF3B1^wt^ and pCMV-3Tag-1A-SF3B1^K700E^ plasmids (37) were designed by Angelos Constantinou (Department of Molecular Bases of Human Diseases, CNRS UPR 1142, IGH-Institute of Human Genetics, Montpellier 34090, France) and kindly provided by Marc-Henri Stern, Institut Curie, Paris, France. pcDNA3.1-FLAG-SF3B1-WT and pcDNA3.1-FLAG-hSF3B1-K700E (18) were obtained from Addgene (#82576 and #82577). The human full-length *SF3B1* sequence has been previously reported to be impossible to clone into bacteria (59,60). Therefore, the plasmids consisted of synthetic sequences, codon-optimized for expression in bacteria (18,37). For the minigene constructs the intron and parts/complete adjacent upstream and downstream exons were PCR-amplified from K562 genomic DNA using Phusion Hot Start Flex polymerase (New England Biolabs) and cloned by the Hot Fusion (61) method into the BamHI restriction site of pcDNA3 (Invitrogen, https://www.addgene.org/vector-database/2092/). The open reading frame of the exons was left intact. The oligonucleotides used for cloning of the constructs are listed in Supplementary Table S4. Mutations and insertions were introduced by site directed mutagenesis using the Q5 site-directed mutagenesis kit (New England Biolabs). Oligonucleotides for site-directed mutagenesis were designed using the NEBaseChanger version 1.3.3 (New England Biolabs) and are listed in Supplementary Table S4. All constructs were verified by Sanger sequencing (Microsynth Seqlab AG, Göttingen, Germany).

### Transfection

The HEK293-FT cells in the amount of 2x10^5^ were seeded per 6-well and incubated over-night. The following day, cells were co-transfected with either 0.5 µg pCMV-3Tag-1A-SF3B1^wt^, pCMV-3Tag-1A-SF3B1^K700E^, pcDNA3.1-FLAG-SF3B1-WT, or pcDNA3.1-FLAG-hSF3B1-K700E and 0.5 µg of the respective minigene construct using the ratio 1:2 of DNA:PEI MAX (PolyScience, #24765-1) according to the manufacturer’s recommendations. Cells were harvested 48 h after transfection. RNA was isolated using the NucleoSpin RNA Mini kit (Macherey-Nagel, #740955.250). DNase digestion was performed on a column as recommended by the manufacturer. We did not observe any obvious difference in the outcome of the minigene assays when comparing the pCMV-SF3B1 and the pcDNA3-SF3B1 constructs.

### cDNA synthesis and validation of the splicing alterations

An amount of 500 ng total RNA was reverse transcribed using SuperScript^TM^ II (Thermo Fisher Scientific, #18064014) and hexamer oligonucleotides for the cDNA synthesis from K562 and Nalm6 RNA. For the minigene assays 500 ng RNA was reverse transcribed using SuperScript^TM^ IV Reverse Transcriptase (Thermo Fisher Scientific, #18090010) with the plasmid specific BGH-rev oligo in 20 µl. Subsequently, RNA in DNA-RNA hybrids was digested by RNase H incubation. For RT-PCR we used 1 µl of cDNA, Taq DNA polymerase, recombinant (Thermo Fisher Scientific, #10342020), and specific oligonucleotides (Supplementary Table S4) in a volume of 25 µl. The PCR run for 35 PCR cycles. PCR products were separated on a 3–4% TAE-agarose gel.

### PacBio Iso-Seq library preparation and sequencing

For the cDNA synthesis we used oligo-dT oligonucleotides and the TeloPrime v2 Kit (Lexogen) to ensure the amplification of full-length mRNAs that contain a Cap-structure. Barcoded primers were used in the cDNA amplification step to enable multiplexing before library preparation. To enrich for slightly larger cDNAs, we adjusted the magnetic bead concentration in the bead clean-up after cDNA amplification. Subsequent library preparation was performed with the Express Template Kit 2.0 (PacBio).

In total, 61 libraries were sequenced on the PacBio Sequel II platform, by multiplexing 4 samples per 8M SMRT cell at the Genomics and Transcriptomics Laboratory, the production site of the West German Genome Center in Düsseldorf (Heinrich-Heine Universität, Düsseldorf, Germany).

### Processing of PacBio Iso-Seq data

Preprocessing of raw Iso-Seq sequencing data was performed with *IsoSeq* (62) software version 3.4, using the recommended parameters. In brief, we used the *ccs* tool to call circular consensus sequences, *lima* to remove primers and adapters, and *isoseq refine* to demultiplex samples and filter out reads not featuring poly-A sequences. This resulted more than 33 million aligned full-length non-chimeric (flnc) polyA HiFi reads (100,302-1,258,653 per library, average 582,135, Supplementary Table S1), with an average length of 2,721 bp. At this sequencing depth, transcript isoforms expressed at one transcript per million (TPM) are expected to be sequenced by at least 25 reads, and two TPM isoforms by at least 50 reads, with more than 95% probability (Supplementary Fig. S1).

According to the base quality values, 57 samples had an error rate of less than 1% in at least 99.7% of the HiFi reads and only four samples had higher error rates with 96.7% to 96.9% reads with < 1% error rate. Overall, the quality of the reads was high, with only 9% of reads potentially affected by technology-specific technical artefacts (20) (Supplementary Fig. S2).

The flnc reads were converted to FASTQ using samtools v.1.18 (63), and, without an additional clustering step, aligned to the human genome GRCh38.p13 using minimap2 (64) version 2.22 with the preset parameters for high quality spliced reads (-ax splice:hq). For each sample, at least 99.85% of the reads were mapped to the genome, and at least 91.9% were uniquely mapped. Samples for which we sequenced more than one library were merged using samtools v.1.18 (63) after the mapping step. Sequencing and mapping statistics per sample are detailed in Supplementary Table S1.

*SF3B1* mutation calling was done with bcftools v.1.13 (65) mpileup and snpEff v.5.1d (66).

Further analysis of Iso-Seq data was performed in python v3, using IsoTools (22) version 0.2.8. In brief, aligned reads were imported and compared to the human reference annotation version 36 from GENCODE, to call, annotate, classify, and quantify transcripts, using IsoTools’ add_sample_from_bam function.

For exploratory analysis, alternative splicing events were detected using IsoTools’ alternative_splicing_events function. For each sample, individual events were quantified by percent spliced index (PSI) values, i.e., the number of reads supporting transcripts that include additional exonic sequence over all transcripts spanning that event. Based on these PSI values, principal component analysis (PCA) plots for different alternative splicing categories were computed using IsoTools’ plot_embedding function.

Differential splicing events between *SF3B1*^mut/wt^ and *SF3B1*^wt/wt^ CLL, MDS and cell line samples were computed with the IsoTools altsplice_test function, using the betabinomial likelihood ratio test. This test models the variability within the tested groups with a beta-binomial mixture distribution, a binomial distribution where the probability parameter, *p*, of the binomial distribution *B(n,p)* follows a beta distribution, *Beta(a, b)*. This formulation allows for considering within-group variability in a similar manner as tests based on negative binomial distribution for RNA-seq data, which is crucial for heterogeneous samples such as individual cancer patients. To be tested, we required the events to be covered by at least 10 reads in at least 4 samples per group, and the minor alternative to be covered by at least 5% of the total reads (test = ’betabinom_lr’, min_n = 10, min_sa = 4, min_alt_fraction = 0.05).

All 3’ alternative splicing events (including those that were not differentially expressed between *SF3B1*^mut/wt^ and *SF3B1*^wt/wt^) were exported for further analysis (Supplementary Table S3).

### Estimation of branch point position and splice-site strength score

For all 3’ alternative splicing events, we used the R Bioconductor package branchpointR (67) to predict branchpoint probabilities for both, canonical and alternative splice sites. We used the position with the highest branchpoint probability as the predicted branch point.

The strength of the 3’ splice sites for the minigene constructs was calculated with SpliceRover (32) (http://bioit2.irc.ugent.be/rover/splicerover, accessed on 23.06.2023 using the model “human acceptors”).

### Transcript coding potential

An ORF was deemed to possess a translation initiation if its 5’ intron chain coincided with that of an annotated protein-coding transcript and the corresponding annotated CDS began at that exact same location. Moreover, we predicted additional translation initiation for the first ORF that fulfilled criteria we inferred from the distribution of the annotated CDS (Supplementary Fig. S17) i.e.,: i) an ORF spanned a length of ≥ 300 nt, (ensuring a CDS length of ≥ 300 nt); ii) the ORF was situated within the initial 500 nt of the transcript (thus ensuring a 5’UTR length of ≤ 500 nt); iii) and the similarity to the Kozak consensus sequence was at least moderate (log odds ratio > -2.5), ensuring a favorable initiation sequence context. Furthermore, the potential for nonsense-mediated mRNA decay (NMD) was assessed for all ORFs, using the 55 nt rule, wherein transcripts with a termination codon located more than 55 nt upstream of the last exon-exon junction are likely subjected to NMD.

### Illumina RNA-seq library preparation and sequencing

RNA from K562 and Nalm6 cells was isolated using the NucleoSpin RNA Mini kit (Macherey Nagel) followed by DNase-digestion with the DNase I set (Zymo Research, #E1010) and clean-up using the NucleoSpin RNA Clean-up Mini Kit (Macherey-Nagel, #740948.50). RNA quality was surveyed using RNA ScreenTapes (Agilent) and the RNA integrity number (RIN) was 10 for all samples. CLL cells were stored in RNA later and RNA was isolated using the RNeasy Mini Kit (Qiagen) followed by DNase digestion using the DNase I Amplification Grade Kit (Invitrogen) and a clean-up using RNeasy Min Elute columns (Qiagen). RNA-seq libraries for the cell lines and the CLL RNA were prepared using the TruSeq-stranded-total RNA SamplePrep kit (Illumina) according to the manufacturer’s protocol. In brief, 2 µg of total RNA were depleted for ribosomal RNA using a Ribo-Zero-rRNA removal kit (Illumina), followed by random primed cDNA synthesis. Sequencing libraries were run at the sequencing core unit of the Max-Planck Institute for Molecular Genetics on a HiSeq2500 (Illumina) using 50 bp paired-end reads.

### Processing of Illumina RNA-seq data

Illumina RNA seq reads were aligned to the human reference genome GRCH38.p13 using STAR aligner version 2.7.6a (68), with provided gff annotation from GENCODE release 36 including annotation of non-chromosomal scaffolds. Alternative splicing events were called and quantified using rMATS (v4.1.1) (30).

Differential expression between *SF3B1*^mut/wt^ and *SF3B*^mut/wt^ in CLL patients and cell lines for genes with at least 5 FPKM in at least 2 samples was performed with edgeR (v 3.30.3) (69) based on the read counts computed with RSEM (70). A gene was considered differentially expressed if the false discovery rate (FDR) was below 0.05 and absolute log2 fold change was at least 1.

Mutations in *SF3B1* were called as for Iso-Seq described above.

### iCLIP experiments

iCLIP of K562-SF3B1^wt/wt^ and K562-SF3B1^K700E/wt^ was performed as described previously (71). To this end, exponentially growing K562 cells were pelleted, washed once with PBS and 10x10^6^ were subjected to UV-cross linking at 400 mJ/cm^2^ at 254 nm in 6 ml PBS in a 10 cm petri dish on ice. Crosslinked cells were scraped from the dish, collected by centrifugation for 2 min at 500 xg at 4 °C, snap frozen in liquid nitrogen and stored at -80 °C. About 3x10^6^ cells were immunoprecipitated using 10 µg of a monoclonal SF3B1 antibody (clone 16, MBL D221-3) or 10 µg of the monoclonal U2AF2 antibody (U4759, SIGMA).

### iCLIP data processing

Initial quality control was done using FastQC (72) before and after quality filtering). All reads having at least one position with a sequencing quality < 10 in the barcode area (positions 1–3, 4–7, 8–9) were removed. De-multiplexing and adapter trimming were done on quality filtered data using Flexbar (73). No mis-matches were allowed during de-multiplexing, while an error rate of 0.1 was accepted when trimming the adapter at the right end of the reads. Furthermore, a minimal overlap of 1 bp between reads and adapter was required and only trimmed reads with a minimal length of 15 bp (24 bp including the barcode) were kept for further analysis. Barcodes of remaining reads were trimmed (but kept as additional information with the reads). Trimmed reads were then mapped to genome assembly version hg38 using STAR (v. 2.6.1b) (68) with 4% mismatched bases allowed and turned off soft-clipping on the 5-prime end, and GENCODE gene annotation v31 (74). While technical duplicates were removed with UMI-tools (75) with *unique* method, all real duplicate reads were kept. Then, we checked the crosslink quality with iCLIPro (76).

For facilitate comparisons, the crosslink events, i.e., reads after duplicate removal, of the replicates were randomly subsampled to the size of the smallest replicate (n = 13,892,358) (replicate 2 of SF3B1^wt/K700E^; Supplementary Table S1).

### Definition and classification of SF3B1 binding sites

Binding sites were identified from the merged crosslink events of SF3B1^wt/wt^ and SF3B1^wt/K700E^ as described in Busch *et al.* (77) (Supplementary Fig. S23). For this, the crosslink events of all five replicates were combined and subjected to peak calling with PureCLIP (version 1.3.1) (78) with default parameters. The PureCLIP-called sites (Psites) were filtered by first removing 5% of the Psites with the lowest score associated and then keeping only the top 20% of Psites within each gene annotated (GENCODE release 36, GRCh38; only annotations with a gene support level of 1 or 2 and transcript support level from 1 to 3). The Psites were then merged into binding sites using the R/Bioconductor package BindingSiteFinder (version 1.0.3) (79), using the following options: width of 5 nt (bsSize =5); ≥ 2 Psites (minWidth = 2, minClSites = 1) and ≥ 1 crosslink positions within each binding site (minCrosslinks = 1). In brief, Psites closer than 5 nt were merged into regions, and isolated Psites were discarded. Within each region, binding site centers were iteratively placed at the position with most crosslink events and extended by 2 nt on both sides. Binding site centers were required to harbor the maximum crosslink signal within the binding site. The optimal binding site width of 5 nts was determined by an evaluation of the ratio of crosslink events within binding sites of increasing width over the mean background signal in flanking windows of the same size (Supplementary Fig. S24). Next, binding sites that were not supported by all replicates in at least one condition (SF3B1^wt/wt^ or SF3B1^wt/K700E^) were filtered out (Supplementary Fig. S25). The threshold for sufficient coverage in a replicate was determined using the 5th percentile and a lower boundary of two crosslink events as described in Busch *et al.* (77). Finally, binding sites were assigned to target genes using GENCODE annotation (release 36, GRCh38; filtered as above) as described in Busch *et al.* (77). In total, this procedure identified 96,852 SF3B1 binding sites in 8,127 genes.

To classify distinct SF3B1 binding patterns in introns, bound regions were defined by merging intronic binding sites within a distance < 55 nt and resizing the obtained regions to 81 nt around the center, resulting in 56,224 regions harboring 87,199 binding sites. Following the approaches suggested in Heyl *et al*. (80), we used unsupervised clustering to separate the crosslink patterns in the bound regions. For this, the crosslink coverage (sum of all replicates) was subjected to min-max normalization (81) within each window (i.e., scaling such that the lowest and highest number of crosslink events are set to 0 and 1, respectively), followed by spline-smoothing using the smooth.spline function (R package stats, version 4.1.0) with lambda 0.2 (spar = 0.2) and inflated dimensions (dim = 150). The normalized and smoothed crosslink coverages were then subjected to dimension reduction using uniform manifold approximation and projection (UMAP) (82) with the umap function (package umap, version 0.2.7) with parameters n_epochs = 5000, n_components = 2, min_dist = 0.01 and n_neighbors = 5 (Supplementary Fig. S29A). The UMAP results were assigned to clusters using density-based clustering of applications with noise (DBSCAN)(83) with the dbscan function (R package dbscan (84), version 1.1, eps = 0.3), with a minimum number of 150 points per cluster (MinPts = 150), yielding three clusters: C1 (n = 35,907 regions), C2 (n = 5,635) and C3 (n = 13,847). Bound regions in cluster C0 (n = 835) were deemed as outliers that could not be assigned to any of the fitted density centers and excluded from further analysis. Bound regions in cluster C3 (wide pattern) were smoothed more finely (spar = 0.1, dim = 500) and then subjected to a second round of UMAP dimension reduction (parameters as above) and DBSCAN clustering (MinPts = 60, eps = 0.23), yielding subclusters #0–33 (Supplementary Fig. S29B). Cluster numbering is based on the increasing distance between the two modes in the arrangement of binding sites, calculated on the summed and smoothed coverages within each cluster using the locmodes function (R package multimode, version 1.5 (85)) (Supplementary Fig. S29C).

### SF3B1-mRNA model generation

The structure of SF3B1 was taken from the cryo-electron microscopy (cryo-EM) structure of the human activated spliceosome (PDB ID: 5Z56 (86) ). Besides SF3B1 and the pre-mRNA bound to SF3B1, proteins interacting with SF3B1 and the bound pre-mRNA (RNA-binding motif protein, x-linked 2 (RBMX2); splicing factor 3A subunit 2 (SF3A2); PHD finger-like domain-containing protein 5A (PHF5A); cell division cycle 5-like protein (CDC5L); and the U2-snRNA) were also extracted from the structure. Only structural information resolved in the PDB was considered for the final model except for the RNA-binding motif protein, x-linked 2. There, to also include the N-terminal part in proximity to the pre-mRNA bound to SF3B1, structural information for the amino acids 1–7 and 113–140 was taken from the cryo-EM structure of the activated human minor spliceosome (PDB ID: 7DVQ (87)). The RBMX2 from PDB ID 7DVQ was aligned to the RBMX2 in our model using UCSF Chimer (88). Atoms not resolved in our model were included into our model, missing bonds within the RBMX2 were created, and a rotamer of K8 not overlapping with the N-terminus was selected using Schrödinger Maestro, v. 2022-4 (89). Amino acids 82–89, 173–190, 205–259, and 311–335 of the SF3B1 not connected to the rest of the protein and not part of pre-mRNA binding were removed. The protein was protonated, missing side chains were created, and N-termini and C-termini of chains were capped with ACE or NME using Schrödinger Maestro, v. 2022-4 (89). Missing loops (amino acids 450–452, and 486–489 in SF3B1 and 36–44 in splicing factor 3A subunit 2) were modeled using a *de novo* loop generation approach, as implemented in Maestro, v. 2022-4 (89). All side chains of the pre-mRNA were removed. Because side chain creation using Maestro led to unsatisfactory results with incomplete bonds within prolines (Supplementary Fig. S34), all proline side chains were further minimized with CCG MOE2022.02 (90) using default parameters, resulting in the final model used for all pre-mRNA transcripts Supplementary Fig. S35). For the K700E mutant, the lysine was mutated to glutamate using CCG MOE2022.02 (90) and the most favored rotamer according to MOE was selected. For each RNA construct, missing side chains of the pre-mRNA were filled using *tleap*, as implemented in AmberTools22 (91).

### Molecular dynamics simulations

The models were solvated in a capped octahedral water box using OPC (92) water with a water shell of at least 12 Å around solute atoms and neutralized using sodium ions. The AMBER22 package of molecular simulations software (92,93) was used in combination with the *ff19SB* force field for protein atoms (94) and the RNA OL3 force field (95) for RNA atoms. MD simulations were performed as described earlier (96). In short, a combination of steepest descent and conjugate gradient minimization was performed while lowering positional restraints on solute atoms from 25 kcal mol^-1^ Å^-2^ to zero, followed by a stepwise heating procedure and lowering harmonic restraints on solute atoms from 1 kcal mol^-1^ Å^-2^ to zero. For each RNA-model complex (downstream BP / SF3B1^wt^; alternative upstream BP’ / SF3B1^wt^; downstream BP / SF3B1^K700E^; alternative upstream BP’ / SF3B1^K700E^) four replicas of 200 ns length each were performed; resulting in 16 simulations per transcript and a total simulation time of 64 μs (20 * 4 * 4 * 200 ns). The 20 transcripts were selected based on: overlap with iCLIP binding site (DoubleNarrow or DoubleWide); median IsoSeq read coverage ≥ 10; AG’– AG distance ≤ 50 nt; both variants not predicted to undergo NMD; and q-value for differential splicing between *SF3B1*^mut/wt^ and *SF3B1*^wt/wt^ of < 0.01 for differentially spliced and > 0.999 for non-differentially spliced transcripts. These filters were applied before the latest version of the protein functionality prediction was implemented in the IsoTools and four of the differentially spliced, as well as three non-differentially spliced transcripts did no longer fulfilled these criteria for the following reasons: longer isoform predicted for NMD: *PHKB*, *PDCD4*; shorter isoform predicted for NMD: *RWDD4*, *TNPO3*, *SEPTIN2*, *RIC8A*, *NAPG* (in addition, q-value for *NAPG* decreased to 0.998). For temperature control the Berendsen thermostat with a collision frequency of 10 ps^-1^ was used. Final simulations were analyzed using CPPTRAJ(97) from AmberTools (91). Contacts between residues were considered if the distance was < 4 Å, as done previously (98). A cutoff of 4 Å was used to describe salt bridges (99). For each frame, the backbone (CA, C, N) of SF3B1 was fitted to the first frame within the respective simulation before computing the RMSF.

## Supplementary Tables

**Supplementary Table S1.** Clinical and technical information on the samples used.

**Supplementary Table S2.** Union of significant alternative splicing events detected in cell lines, CLL, or MDS patients.

**Supplementary Table S3.** Significant alternative splicing events detected in merged dataset.

**Supplementary Table S4.** Oligonucleotide sequences used in the experiments.

**Supplementary Table S5.** Differential expression and gene overrepresentation results.

**Supplementary Table S6.** SF3B1 binding sites detected with the region class.

## Declarations

### Ethics approval

The study was approved by the Ethics Committee of the University of Cologne (Ethikvotum 11-319 from 11th December 2011, with an amendment from 7th June 2016) and the Ethics Committee of the University of Düsseldorf (Ethikvotum 3768, amendment from 24^th^ October 2018). Informed consent has been obtained from all patients involved.

### Consent for publication

Not applicable

### Availability of data and materials

Cell lines transcriptome raw FASTQ files or PacBio CCS unaligned BAM files were deposited to Sequence Read Archive under PRJNA1037338 project ID, whereas patient data were deposited to European Genome-Phenome Archive under EGAS50000000053 study ID. iCLIP raw FASTQ data, as well as the analysis files were deposited to Gene Expression Omnibus archive under GSE247658. All data is available to reviewers upon request and will be released upon manuscript acceptance for a publication. Patient data access will be controlled by the Data Access Committee at the Institute for Translational Epigenetics at the University Hospital Cologne, University of Cologne, Cologne, Germany.

The complete IsoSeq/iCLIP data analysis is available as a Jupyter Notebook in the GitHub repository https://github.com/ZarnackGroup/go_long2023.

### Authors’ contributions

M.R.S., C.G., R.H., K.Z., H.G., N.G., M.H., and J.K. designed the study. H.H., L.B., A.K., K.B., K.K. and C.D.H. acquired the data. A.P., M.L., M.B., J.K., H.G., C.G., R.H., K.Z., and M.R.S. analyzed and interpreted the data. A.P., M.L., C.G., R.H., K.Z., and M.R.S. drafted and wrote the manuscript. All authors have read and approved the final manuscript.

## Acknowledgements

We acknowledge IMB Genomics Core Facility and its NextSeq 500 sequencer [funded by the Deutsche Forschungsgemeinschaft (DFG, German Research Foundation) INST 247/870-1 FUGG]. We would like to thank Angelos Constantinou (IGH-Institute of Human Genetics, France) and Marc-Henri Stern (Institut Curie, Paris, France) for sharing the *SF3B1* plasmids, Anke Busch (IMB Mainz, Germany) for iCLIP data pre-processing, Elena Wasserburger-Zichel (University Hospital Cologne, Germany), and Bernd Timmermann (Sequencing Core Unit, Max Planck Institute for Molecular Genetics, Berlin, Germany) for their technical assistance. We furthermore thank the Regional Computing Center of the University of Cologne (RRZK) for providing computing time on the DFG-funded (Funding number: INST 216/512/1FUGG) High Performance Computing (HPC) system CHEOPS as well as IT support. In addition, we acknowledge computational support of the Centre for Information and Media Technology, especially the HPC team at the Heinrich-Heine University, as well as the computing time provided by the John von Neumann Institute for Computing on the supercomputer JUWELS at Jülich Supercomputing Centre (user IDs: VSK33). This work was supported by the DFG Research Infrastructure West German Genome Center (407493903) as part of the Next Generation Sequencing Competence Network (project 423957469). NGS analyses were carried out at the West German Genome Center, and production sites in Cologne and Düsseldorf.

