## Supplementary Figures for "Long-read transcriptome sequencing of CLL and MDS patients uncovers molecular effects of *SF3B1* mutations"

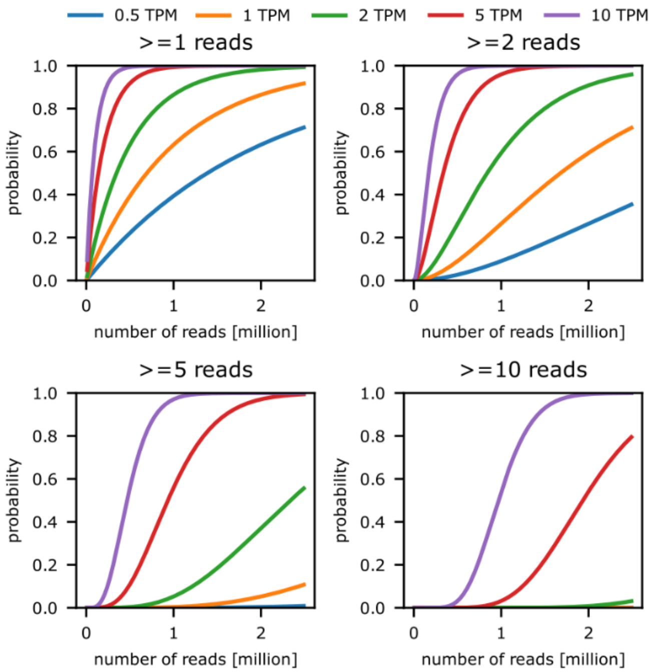


**Supplementary Figure S1.** Read saturation analysis in IsoSeq data analysed. The probability of a transcript expressed at 0.5, 1, 2, 5, or 10 transcripts per million (TPM) to be identified at sequence depth of 0 to 2.5 million reads with at least 1, 2, 5, or 10 reads.

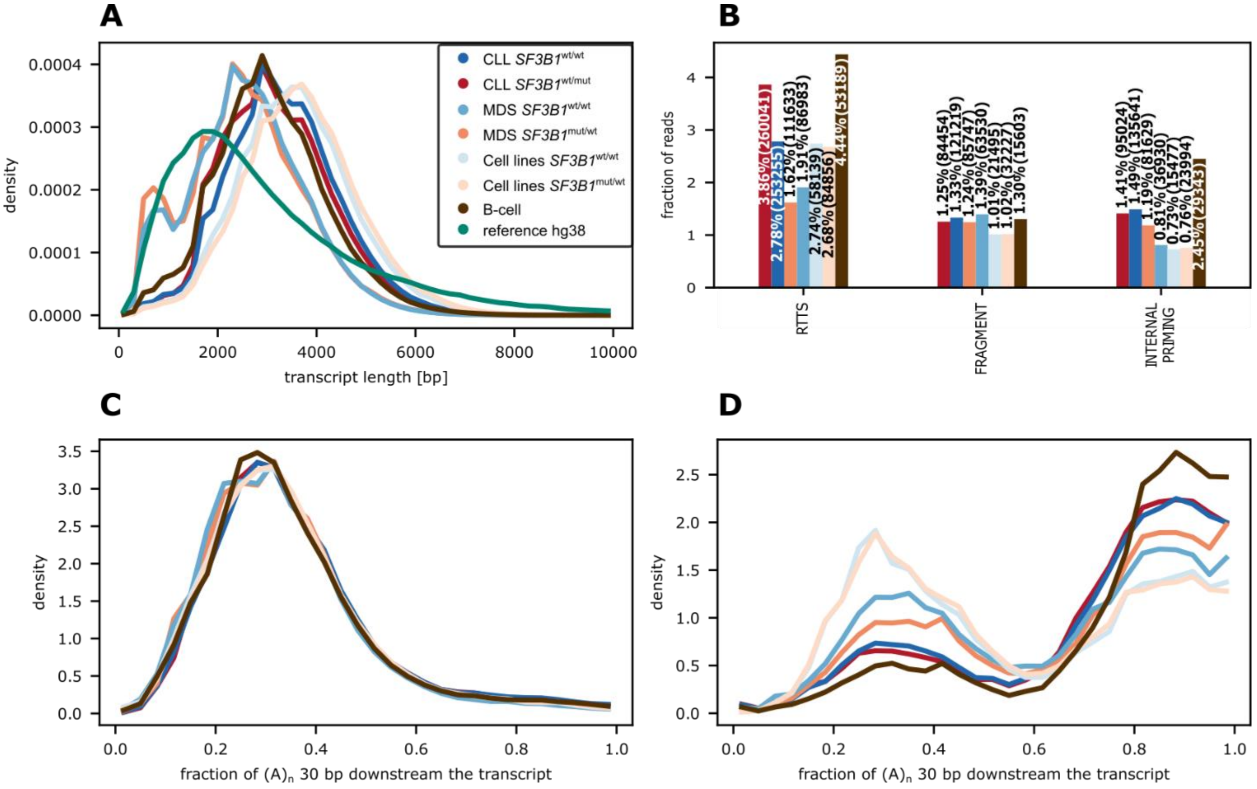


**Supplementary Figure S2**. Quality control of the Iso-Seq reads in eight datasets: cell lines (CL), CLL, MDS, each separated by the *SF3B1* mutational status, as well as normal B cells from this and the ENCODE studies. **A** – Density distribution of transcript length sequenced in each of the sample group. The y axis was cut at 10,000 nt. **B** – Percentage of reads within each of artefacts type per group of samples. **C** – Density distribution of the fraction of adenosines (A) within 30 bp downstream the transcript within known transcript annotated in GENCODE version 36. **D** – Same as C but for novel transcripts.

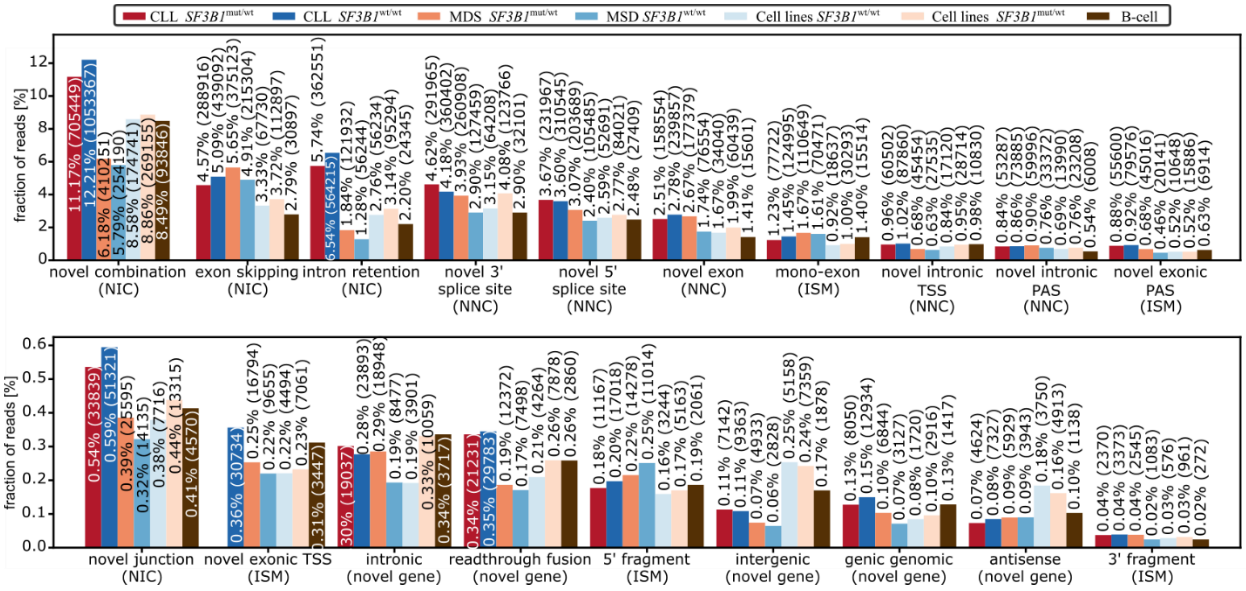


**Supplementary Figure S3.** Novel isoforms identified with Iso-Seq separated by transcript type in all group investigated. Only substantial transcripts were taken into account.

SF3B1

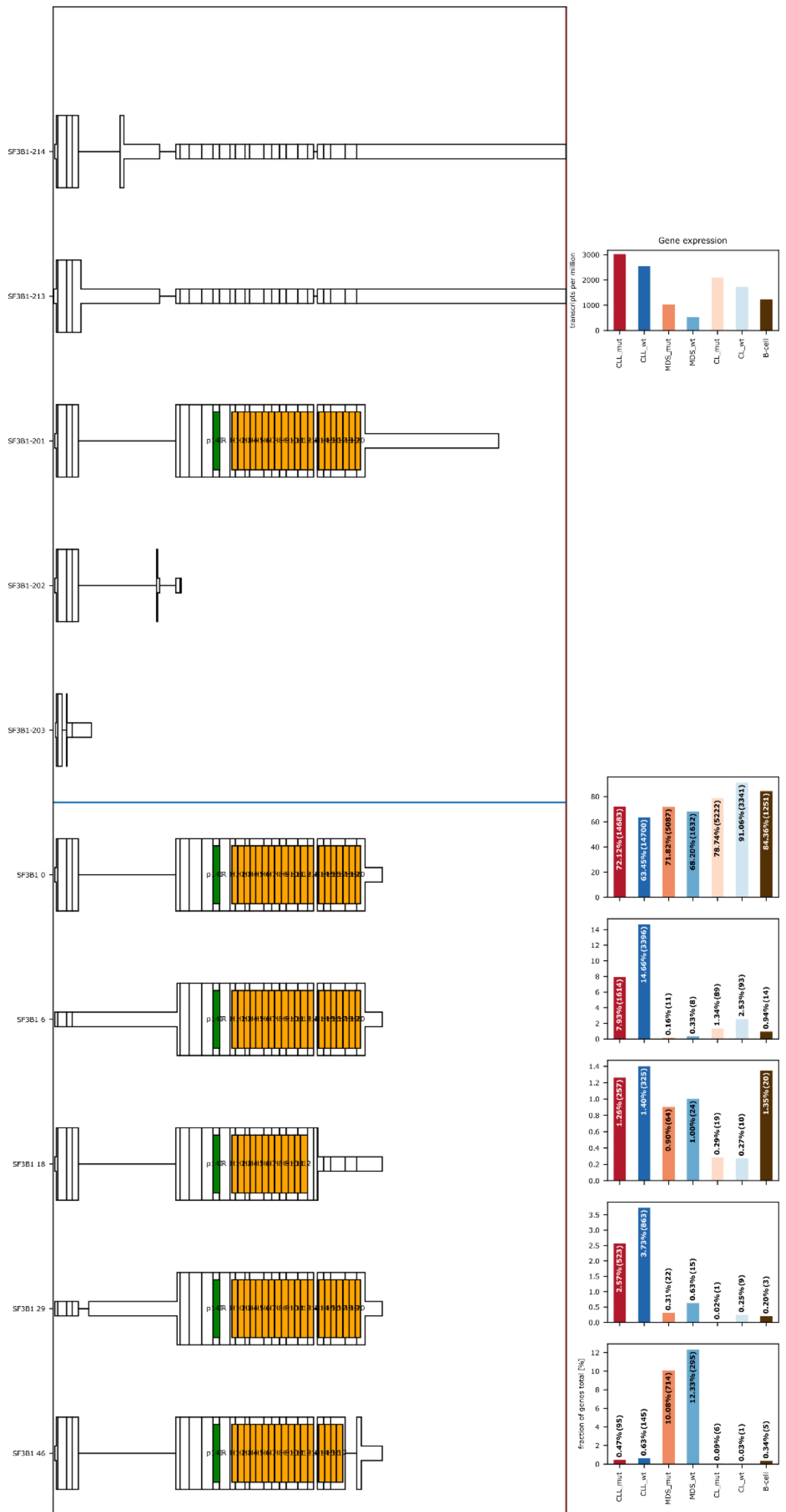

SF3B1-214
SF3B1-213
SF3B1-201
SF3B1-202
SF3B1-203
SF3B1 0
SF3B1 6
SF3B1 18
SF3B1 29
SF3B1 46
0
2000
4000
6000
8000
Gene expression
transcripts per million
CLL_mut
CLL_wt
MDS_mut
MDS_wt
CL_mut
CL_wt
B-cell
72.12%(14683)
63.45%(14700)
71.82%(5087)
68.20%(1632)
78.74%(5222)
91.06%(3341)
84.36%(1251)
7.93%(1614)
14.66%(3396)
0.16%(11)
0.33%(8)
1.34%(89)
2.53%(93)
0.94%(14)
1.26%(257)
1.40%(325)
0.90%(64)
1.00%(24)
0.29%(19)
0.27%(10)
1.35%(20)
2.57%(523)
3.73%(863)
0.31%(22)
0.63%(15)
0.02%(1)
0.25%(9)
0.20%(3)
0.47%(95)
0.63%(145)
10.08%(714)
12.33%(295)
0.09%(6)
0.03%(1)
0.34%(5)
fraction of genes total [%]

**Supplementary Figure S4.** *SF3B1* substantially expressed isoforms. Blue lines separate GENCODE annotated isoforms (top) and Iso-Seq isoforms (bottom). For each Iso-Seq isoform, the fraction of reads identified for each isoform is shown on the right.

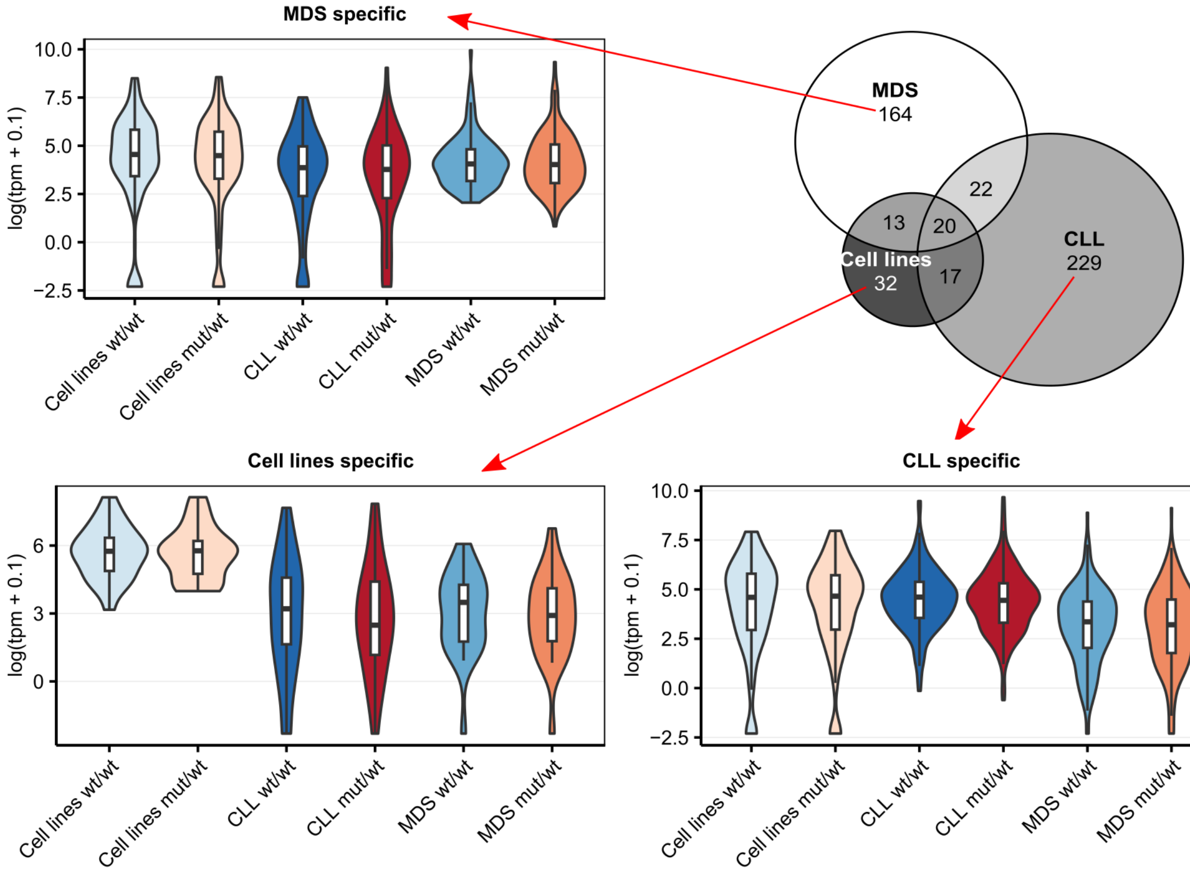


**Supplementary Figure S5.** Overlap of significant alternative splicing events (ASEs) detected using the three sets (cell lines, CLL and MDS patients separately. Although most of the events seem set-specific, the dPSI has high correlation among the datasets (see Fig. 2). For each subset-specific ASE, the overall expression at gene level per group is shown to highlight the fact, the specificity seems to be related to cell-specific transcriptomic profile, rather than specific SF3B1$^{mut}$ mode of action in each dataset.

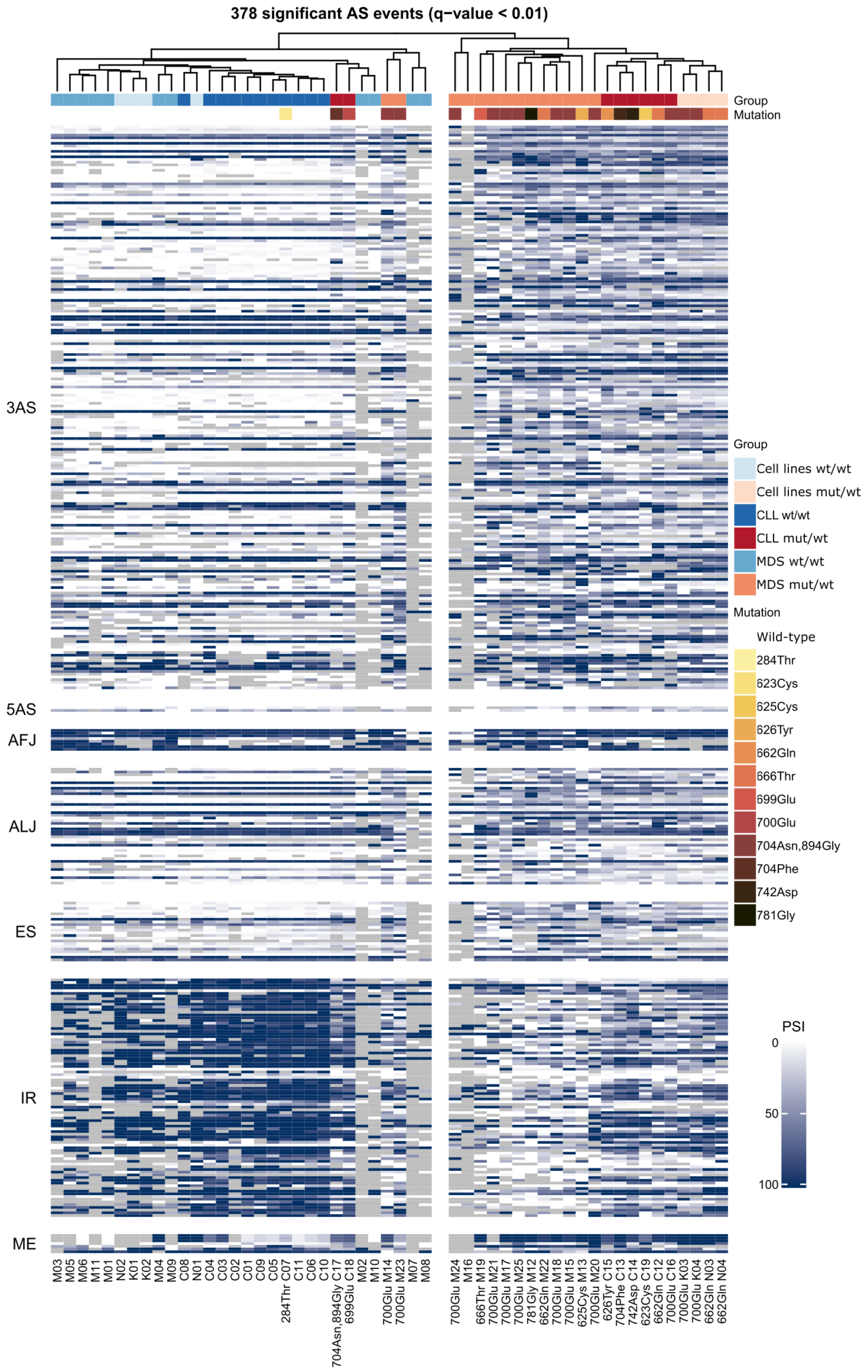

378 significant AS events (q−value < 0.01)
Group
Mutation
3AS
5AS
AFJ
ALJ
ES
IR
ME
Group
Cell lines wt/wt
Cell lines mut/wt
CLL wt/wt
CLL mut/wt
MDS wt/wt
MDS mut/wt
Mutation
Wild-type
284Thr
623Cys
625Cys
626Tyr
662Gln
666Thr
699Glu
700Glu
704Asn,894Gly
704Phe
742Asp
781Gly
PSI
0
50
100
M03
M05
M06
M11
M01
N02
K01
K02
M04
M09
C08
N01
C04
C03
C02
C01
C09
C05
284Thr C07
C11
C06
C10
704Asn,894Gly C17
699Glu C18
M02
M10
700Glu M14
700Glu M23
M07
M08
700Glu M24
M16
666Thr M19
700Glu M21
700Glu M17
700Glu M25
781Gly M12
662Gln M22
700Glu M18
700Glu M15
625Cys M13
700Glu M20
626Tyr C15
704Phe C13
742Asp C14
623Cys C19
662Gln C12
700Glu C16
700Glu K03
700Glu K04
662Gln N03
662Gln N04

**Supplementary Figure S6.** A subset of the most significant alternative splicing events (p.adjusted < 0.01) affected by *SF3B1* mutations in leukemia cell lines as well as CLL and MDS patients show increased 3' alternative splice sites usage and decreased intron retention in samples with *SF3B1* mutation.

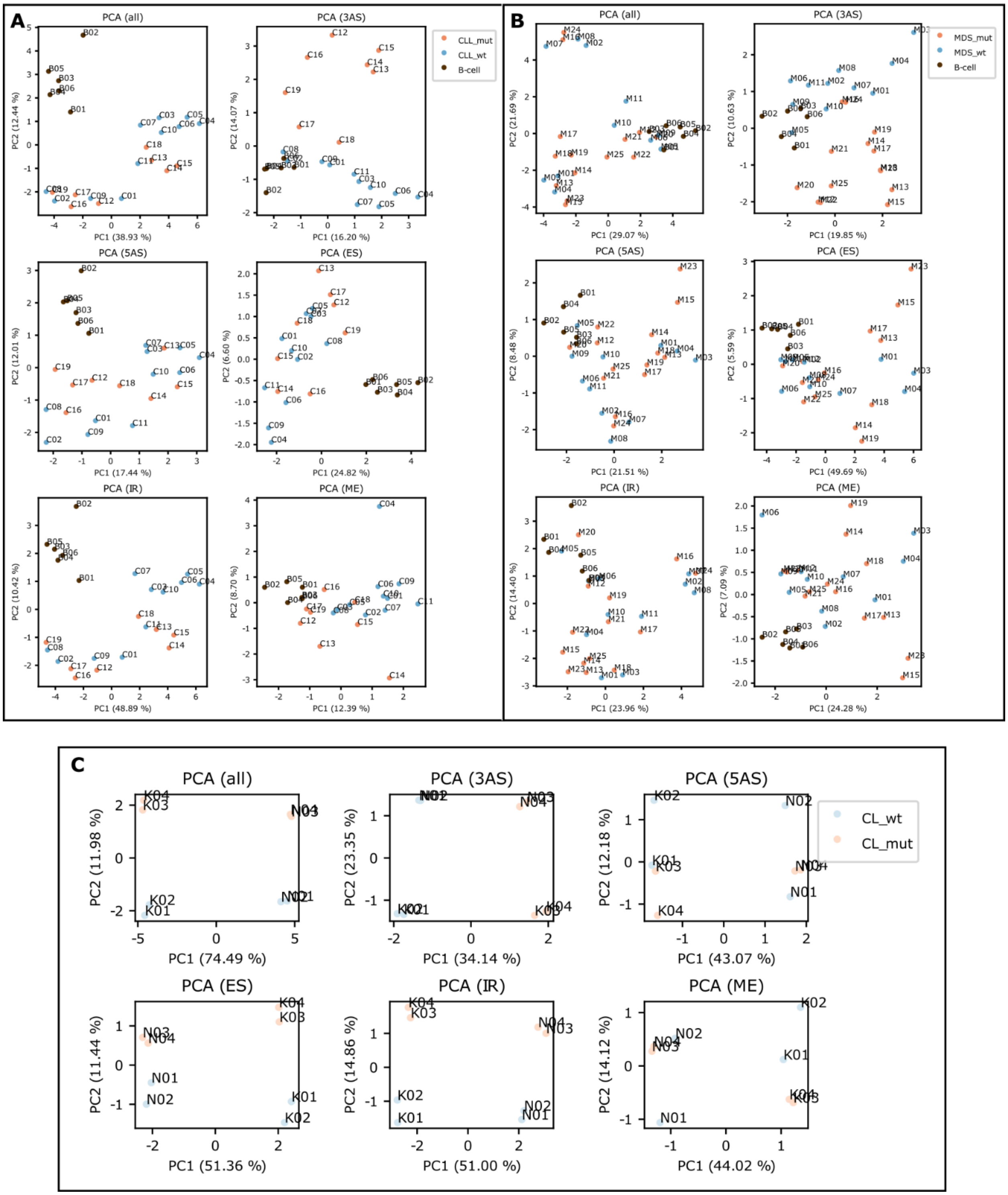


**Supplementary Figure S7.** Principal component (PC) analysis based on the isoform usage of CLL (**A**). MDS (**B**), or cell lines samples (**C**) using all or each of the splice event types separately.

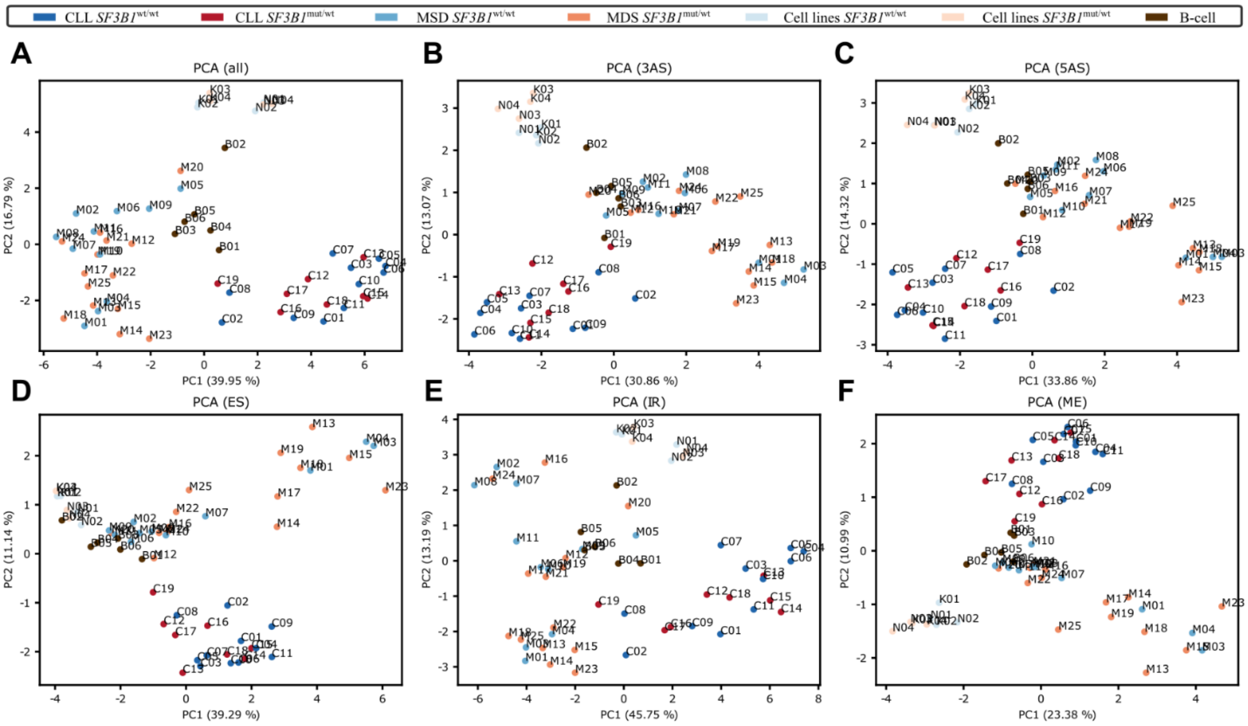


**Supplementary Figure S8.** Principal component (PC) analysis based on the isoform usage of all (**A**) or single type splicing events (**B-F**).

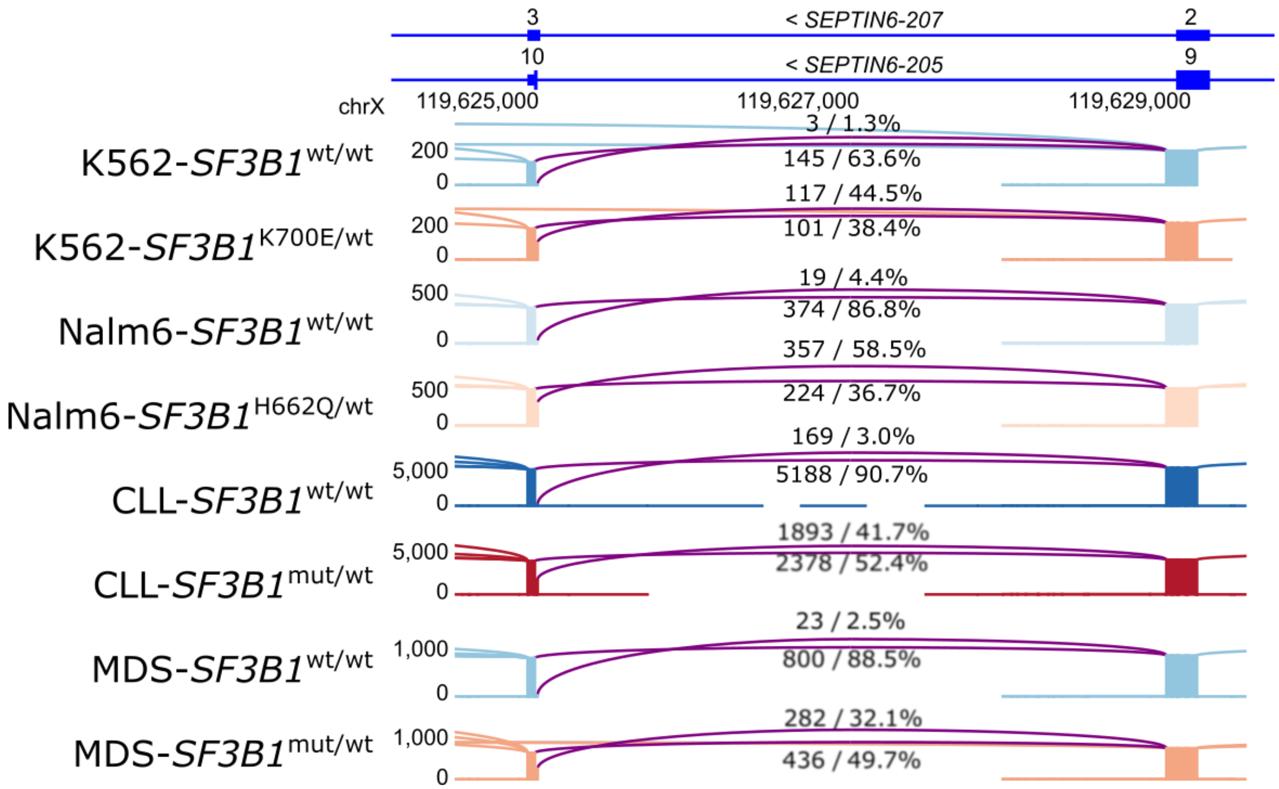


**Supplementary Figure S9.** *SEPTIN6* isoforms with already reported 3'AS event in *SF3B1*$^{mut/wt}$. Each track shows sum of the reads identified in each group. The number of reads supporting each junction and percentage is shown. The width of the lines correlates with significance. Significant differences are coloured in purple.

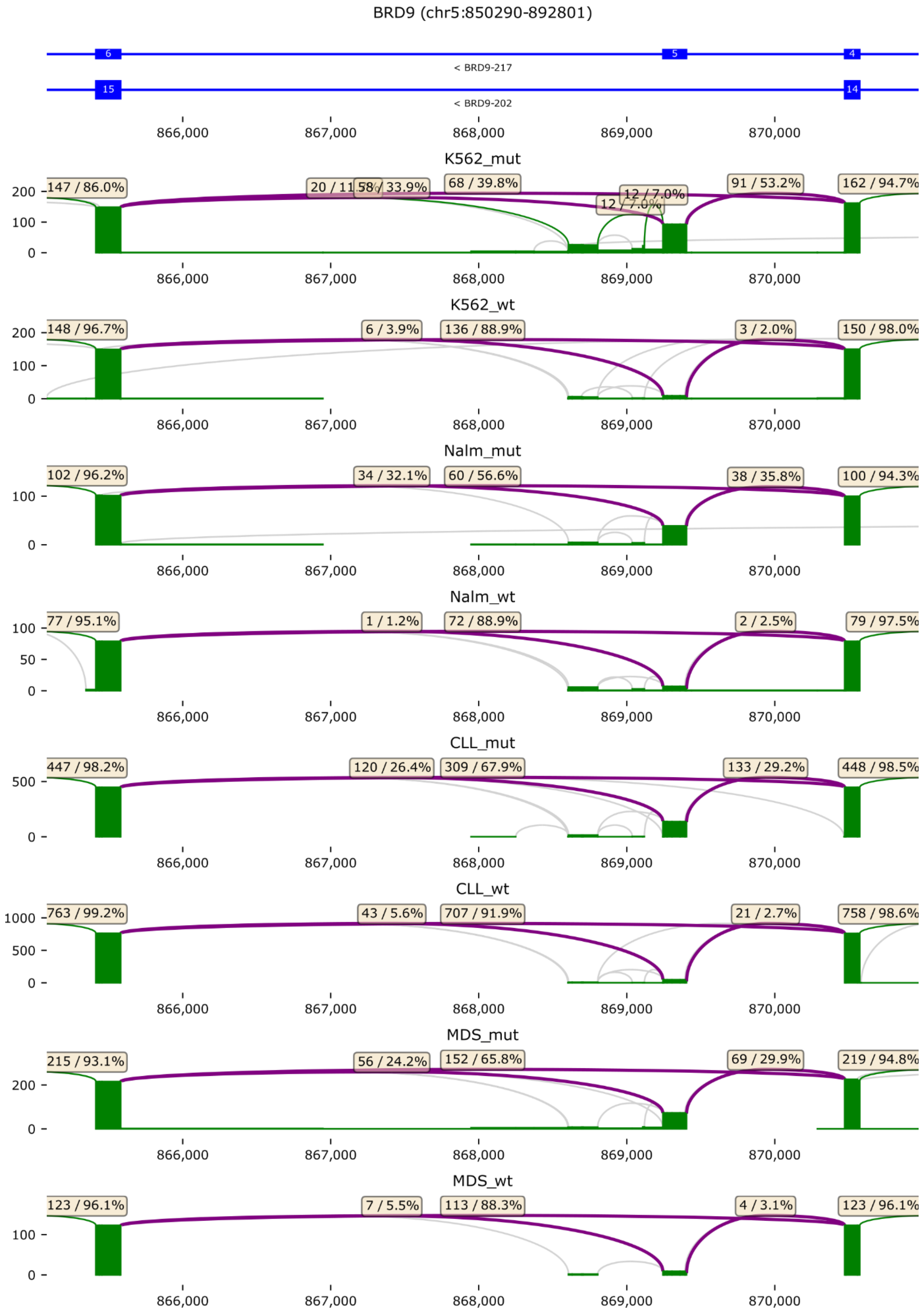
BRD9 (chr5:850290-892801)
< BRD9-217
< BRD9-202
866,000
867,000
868,000
869,000
870,000
K562_mut
147 / 86.0%
58 / 33.9%
68 / 39.8%
12 / 7.0%
91 / 53.2%
162 / 94.7%
K562_wt
148 / 96.7%
6 / 3.9%
136 / 88.9%
3 / 2.0%
150 / 98.0%
Nalm_mut
102 / 96.2%
34 / 32.1%
60 / 56.6%
38 / 35.8%
100 / 94.3%
Nalm_wt
77 / 95.1%
1 / 1.2%
72 / 88.9%
2 / 2.5%
79 / 97.5%
CLL_mut
447 / 98.2%
120 / 26.4%
309 / 67.9%
133 / 29.2%
448 / 98.5%
CLL_wt
763 / 99.2%
43 / 5.6%
707 / 91.9%
21 / 2.7%
758 / 98.6%
MDS_mut
215 / 93.1%
56 / 24.2%
152 / 65.8%
69 / 29.9%
219 / 94.8%
MDS_wt
123 / 96.1%
7 / 5.5%
113 / 88.3%
4 / 3.1%
123 / 96.1%

**Supplementary Figure S10.** *BRD9* isoforms with already reported *BRD9* toxic exon inclusion in *SF3B1*$^{mut/wt}$. Each track shows sum of the reads identified in each group. The number of reads supporting each junction and percentage is shown. Significantly different splicing event are shown as purple arcs whereas non-significantly different splicing events supported by > 5% reads are shown as green arcs, and events supported by > 0.1% reads as grey arcs.

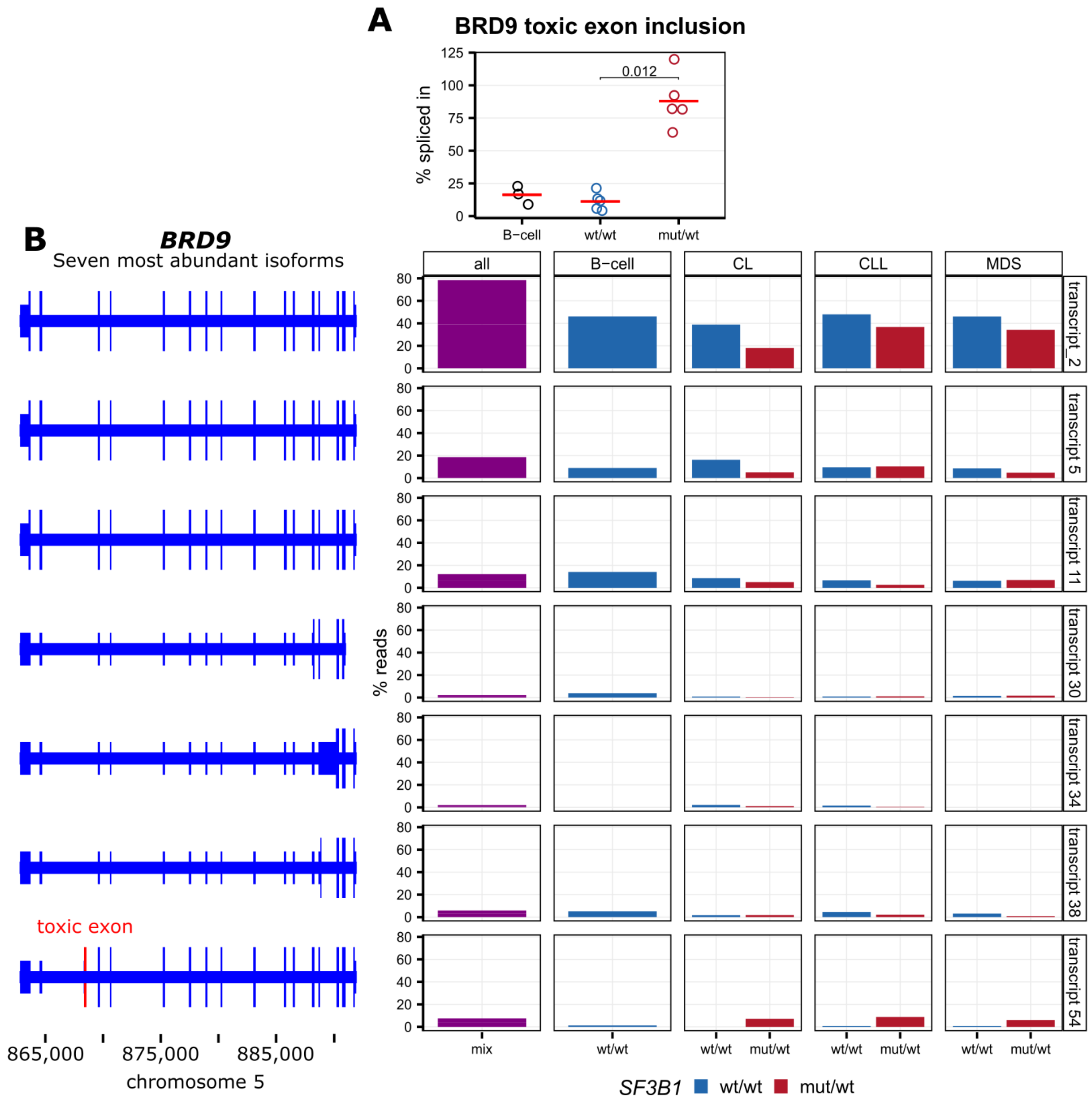


**Supplementary Figure S11.** BRD9 exon inclusion in *SF3B1* mutated samples. **A** – Proportion of the exon inclusion based on qPCR using a subset of CLL and healthy B cell samples used in IsoSeq. **B** – Most abundant isoforms of *BRD9* identified with IsoSeq. **C** - Expression shown as percentage of total reads mapped to *BRD9* gene. The toxic exon is marked in red (B) and was identified in transcript 54 isoform (C), which had a higher proportional expression in samples with an *SF3B1* mutation.

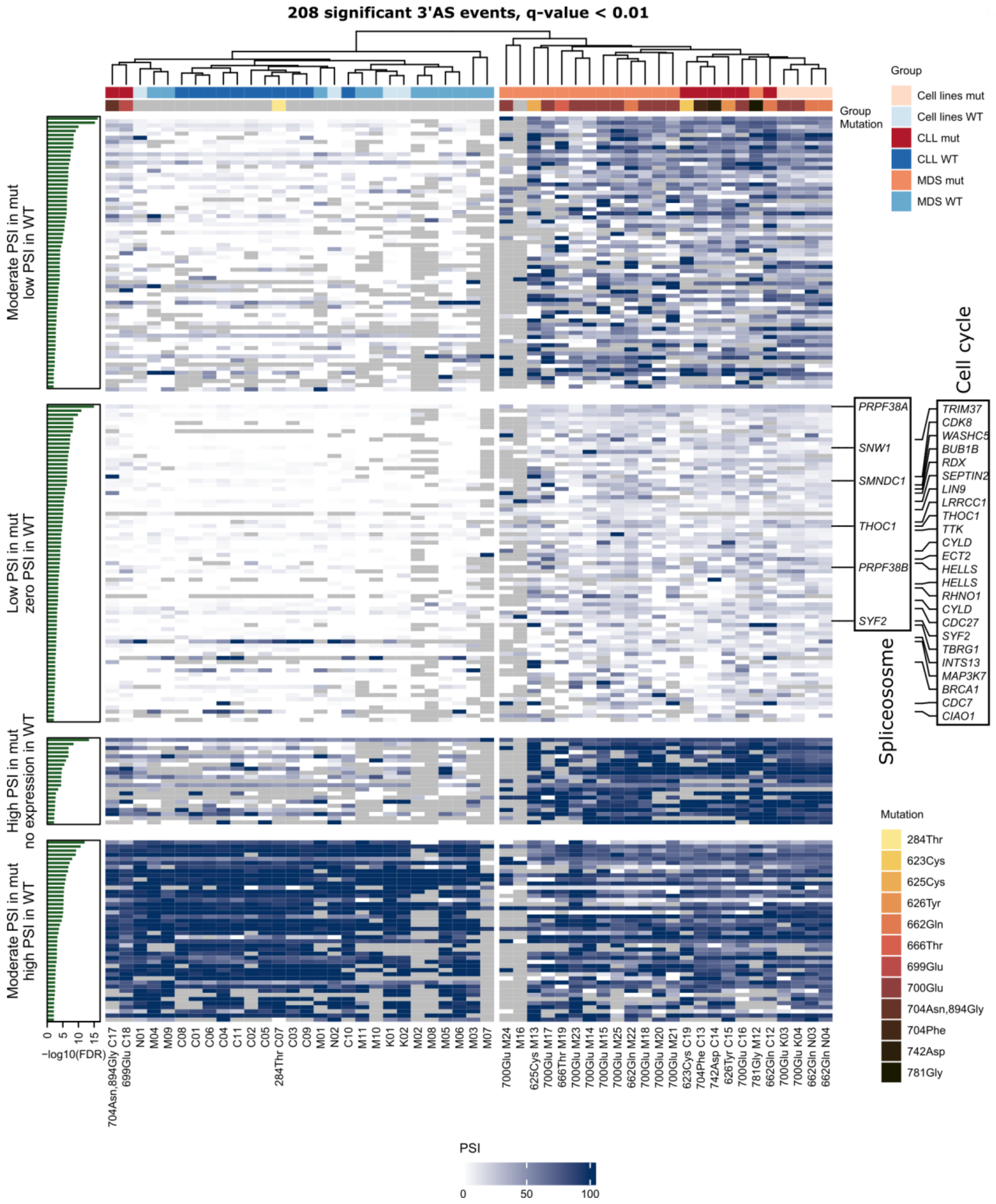


**Supplementary Figure S12.** Clusters of 3' alternative splice sites strongly affected by *SF3B1* mutations (p.adjusted < 0.01) in leukaemia cell lines as well as CLL and MDS patients show enrichment in spliceosome and cell cycle pathways among events with low occurrence in *SF3B1*$^{mut/wt}$ and very low to none in *SF3B1*$^{wt/wt}$.

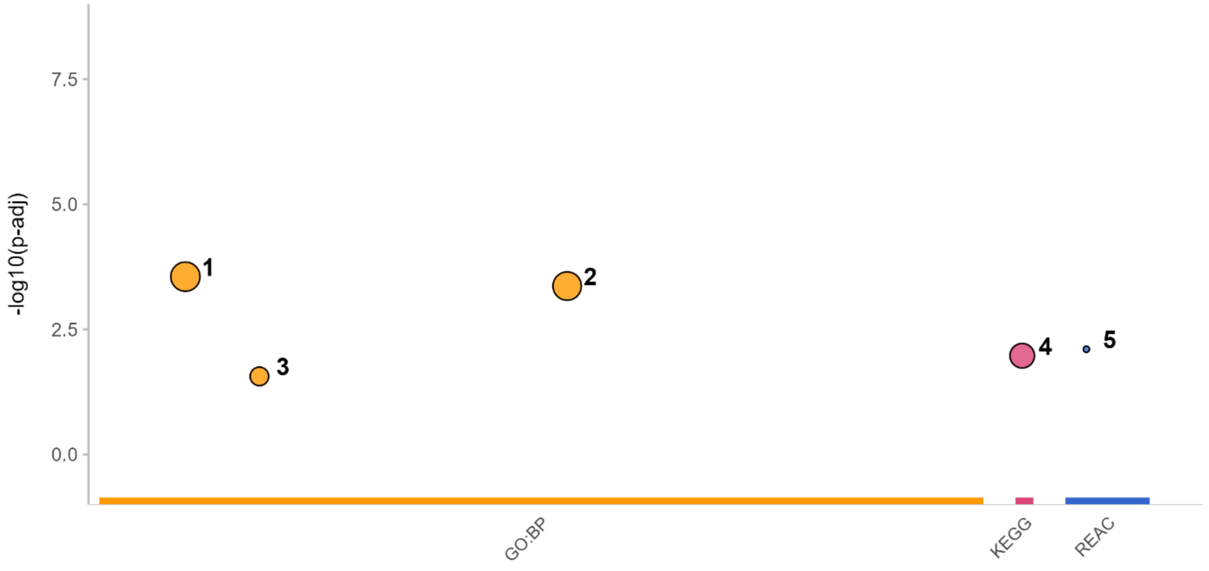


| id | source | term_id | term_name | term_size | | p_value |
|---|---|---|---|---|---|---|
| 1 | GO:BP | GO:0007049 | cell cycle | 1811 | | 2.8e-04 |
| 2 | GO:BP | GO:0051726 | regulation of cell cycle | 1112 | | 4.3e-04 |
| 3 | GO:BP | GO:0015803 | branched-chain amino acid transport | 14 | | 2.7e-02 |
| 4 | KEGG | KEGG:03040 | Spliceosome | 150 | | 1.1e-02 |
| 5 | REAC | REAC:R-HSA-5660862 | Defective SLC7A7 causes lysinuric protein intolerance (LPI) | 2 | | 7.9e-03 |

g:Profiler (biit.cs.ut.ee/gprofiler)

**Supplementary Figure S13.** Overrepresentation analysis of the genes with 3’AS belonging to “Low in mut, zero in WT” cluster from Supplementary Figure 11. “p_value” – adjusted p-value.

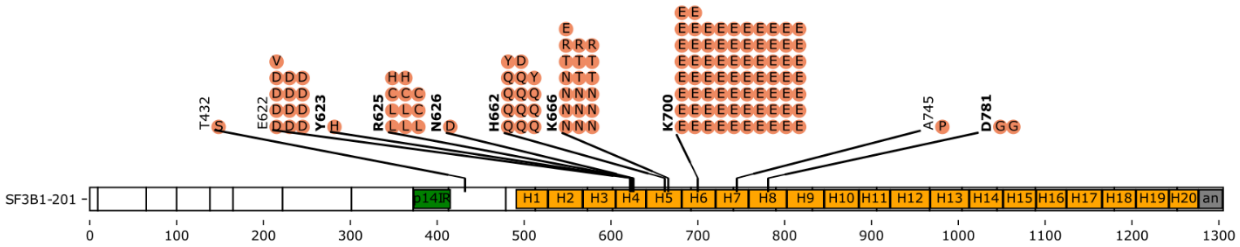


**Supplementary Figure S14.** *SF3B1* mutations detected in the samples used in this and publicly available MDS patients' RNA-seq data (see Supplementary Table S1 for details).

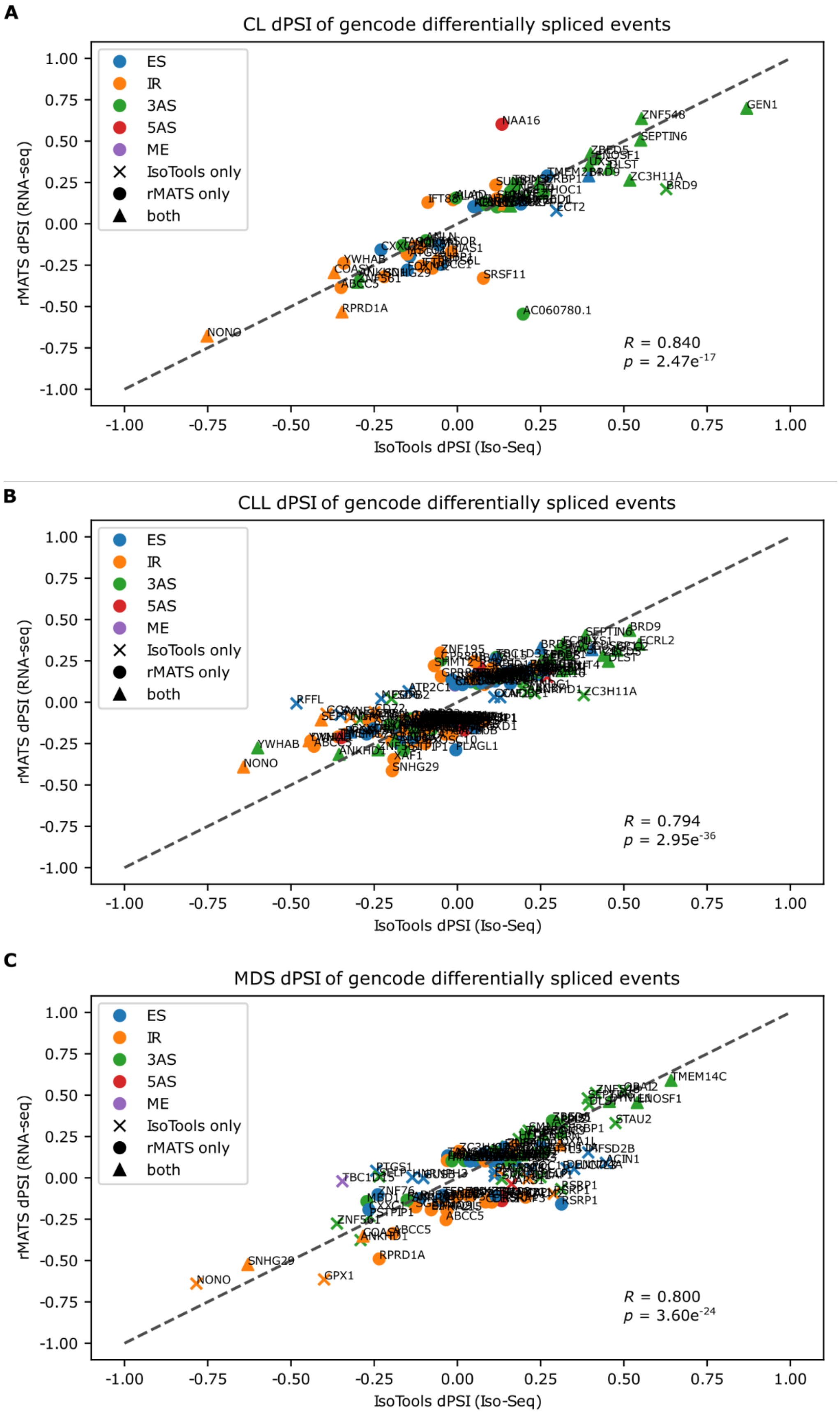
A
CL dPSI of gencode differentially spliced events
ES
IR
3AS
5AS
ME
IsoTools only
rMATS only
both
rMATS dPSI (RNA-seq)
IsoTools dPSI (Iso-Seq)
1.00
0.75
0.50
0.25
0.00
-0.25
-0.50
-0.75
-1.00
GEN1
ZNF548
SEPTIN6
NAA16
ZC3H11A
BRD9
SRSF11
AC060780.1
RPRD1A
NONO
YWHAB
$R$ = 0.840
$p$ = 2.47e$^{-17}$
B
CLL dPSI of gencode differentially spliced events
ES
IR
3AS
5AS
ME
IsoTools only
rMATS only
both
rMATS dPSI (RNA-seq)
IsoTools dPSI (Iso-Seq)
1.00
0.75
0.50
0.25
0.00
-0.25
-0.50
-0.75
-1.00
BRD9
FCRL2
ZNF195
ZC3H11A
RFFL
YWHAB
NONO
SNHG29
PLAGL1
ANKHD1
$R$ = 0.794
$p$ = 2.95e$^{-36}$
C
MDS dPSI of gencode differentially spliced events
ES
IR
3AS
5AS
ME
IsoTools only
rMATS only
both
rMATS dPSI (RNA-seq)
IsoTools dPSI (Iso-Seq)
1.00
0.75
0.50
0.25
0.00
-0.25
-0.50
-0.75
-1.00
TMEM14C
STAU2
MFSD2B
ACIN1
RSRP1
ABCC5
RPRD1A
SNHG29
NONO
GPX1
ANKHD1
$R$ = 0.800
$p$ = 3.60e$^{-24}$

**Supplementary Figure S15.** Comparison of the *SF3B1* mutations effect on splicing using Iso-Seq coupled with IsoTools and RNA-seq coupled with rMATS analyses in A) K562 and Nalm6 cell lines, B) CLL, and C) MDS patients. The colour codes for splicing event type, whereas the point shape indicates significance detected by each of the method used. Only events affecting known isoforms annotated in GENCODE v36 were taken into consideration. B) CLL samples: for the RNA-seq, 7 additional patients' samples were sequenced whereas in C) MDS IsoSeq samples are from this study and RNA-seq samples are derived from publicly available datasets (see Supplementary Table S1 for details).

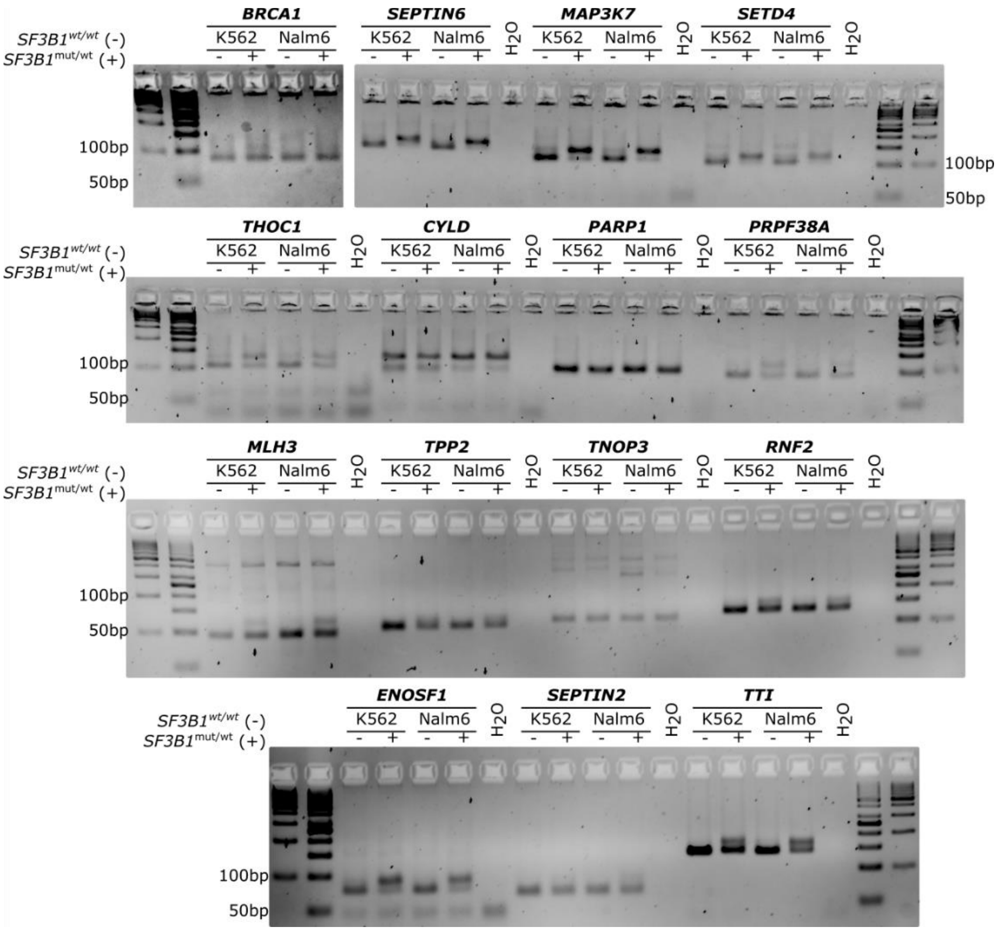


**Supplementary Figure S16.** Validation of alternative splicing detected with Iso-Seq. Agarose gels showing alternative splicing resulting in longer fragments in the *SF3B1*$^{\text{mut/wt}}$ cell lines.

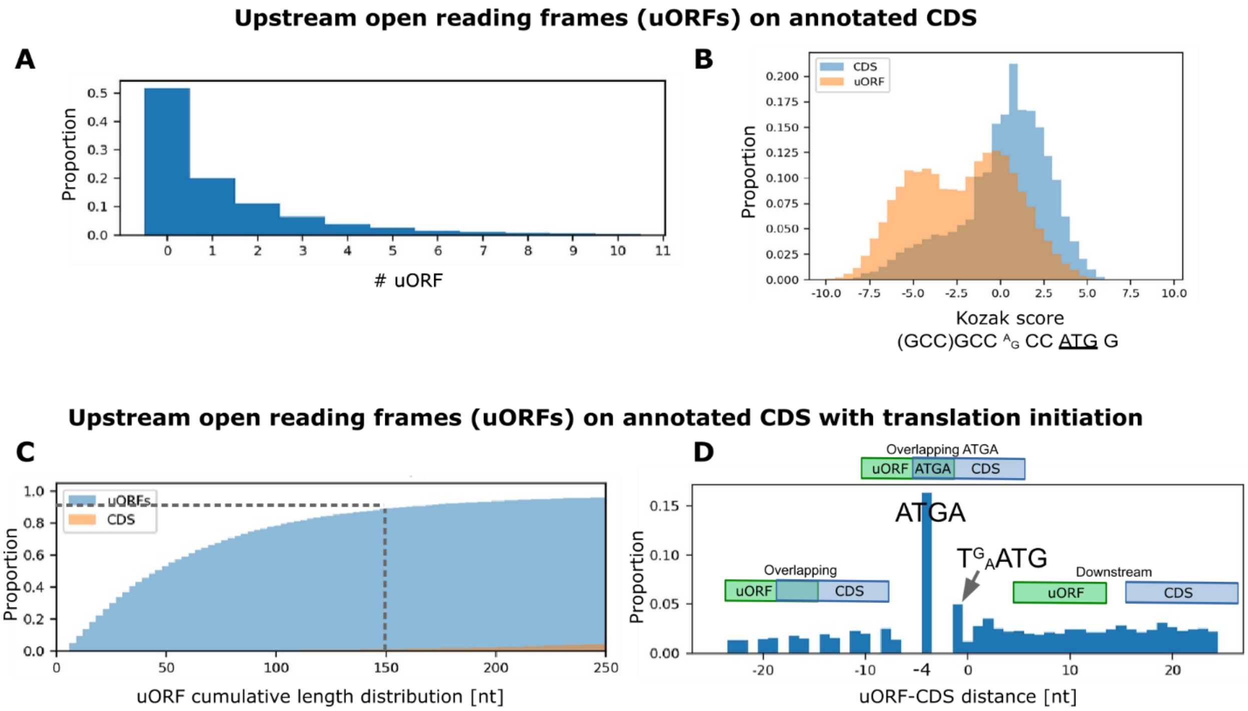


**Supplementary Figure S17.** Investigation of upstream open reading frames (ORFs) on annotated coding DNA sequences (CDS). **A** – The distribution of the number of uORFs per CDS showing > 50% protein coding transcript have upstream start codons. **B** – The distribution of Kozak sequence[1] scores on annotated CDS. **C** – uORF length on annotated CDS showing 99% of CDSs > 150 bases, whereas around 99% of uORFs were < 150 bases [2]. **D** – The distribution of the distance between uORF and CDS. Negative values denote overlapping sequences. The most common distance observed was four bases overlap and these bases were ATGA. The overlapping uORF-CDS were observed up to 50 bases. The downstream instances were most common and dropped down slowly at a distance around 300 bases.

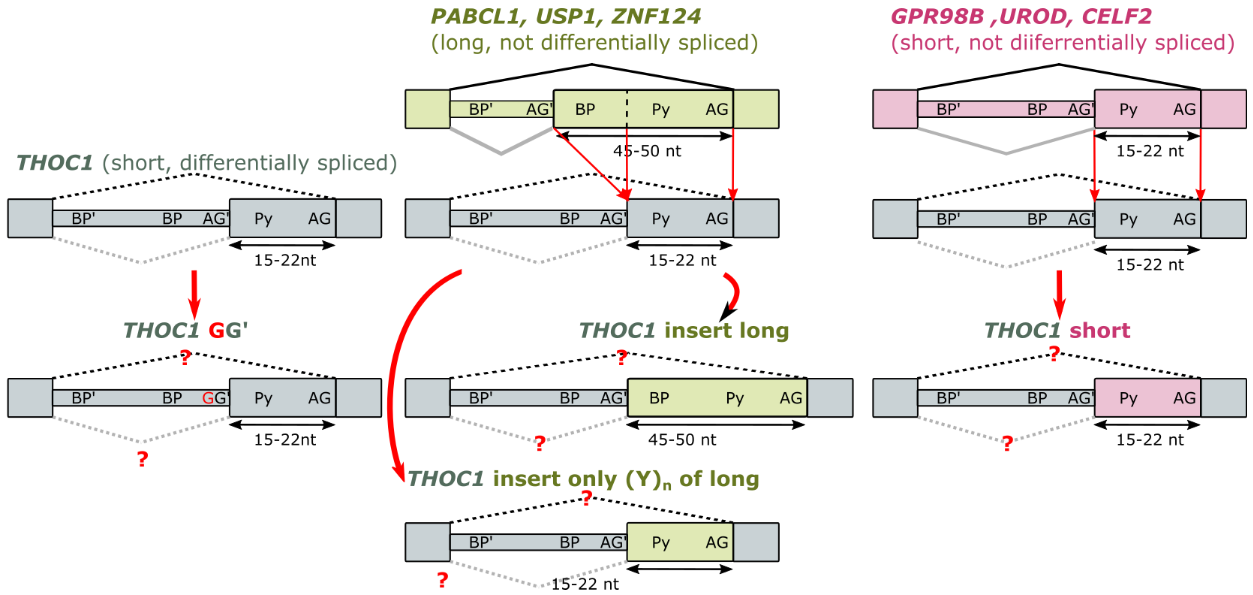


**Supplementary Figure S18.** Design of the minigene assay shown in Figure 3B. Briefly, the intron fragment between AG' and AG from differentially spliced *THOC1* was replaced with a long intron AG'-AG fragment from a not differentially spliced gene (*PABCL1*, *USP1*, or *ZNF124* shown in green), or a short intron AG'-AG fragment from a not differentially spliced gene (*GRP98B*, *UROD*, or *CELF2* shown in pink). Additionally, the long fragments were cut to match shorter fragment length but with polypyrimidine tract (Py) preserved.

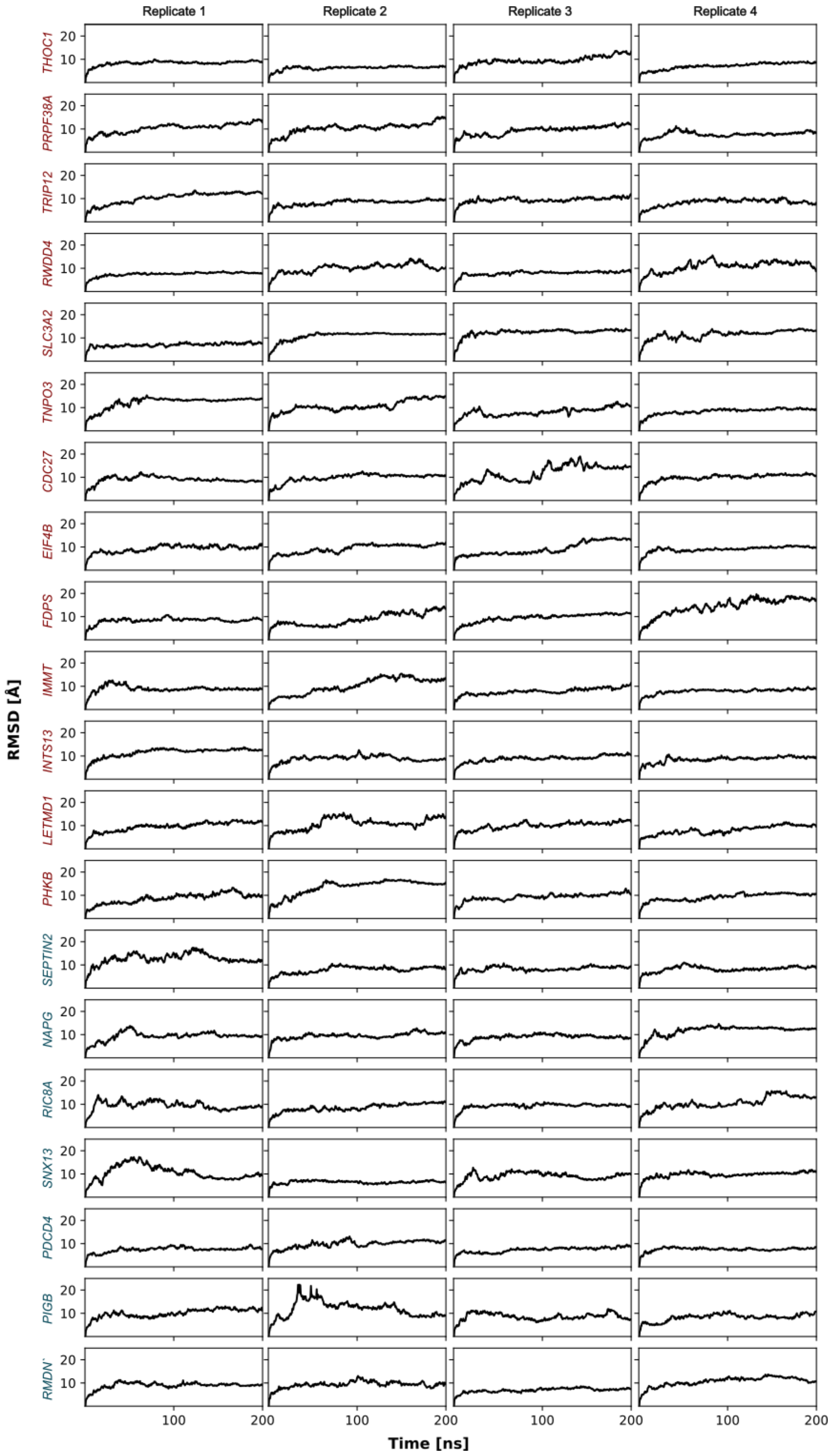


**Supplementary Figure S19.** Root mean square deviation (RMSD) of the SF3B1 backbone (CA, C, N) for molecular dynamics simulations of the downstream BP bound to SF3B1$^{wt}$. mRNAs with 3’ alternative splice site differentially spliced between SF3B1$^{mut/wt}$ and SF3B1$^{wt/wt}$ mRNAs are shown in red and non-differentially spliced are shown in blue.

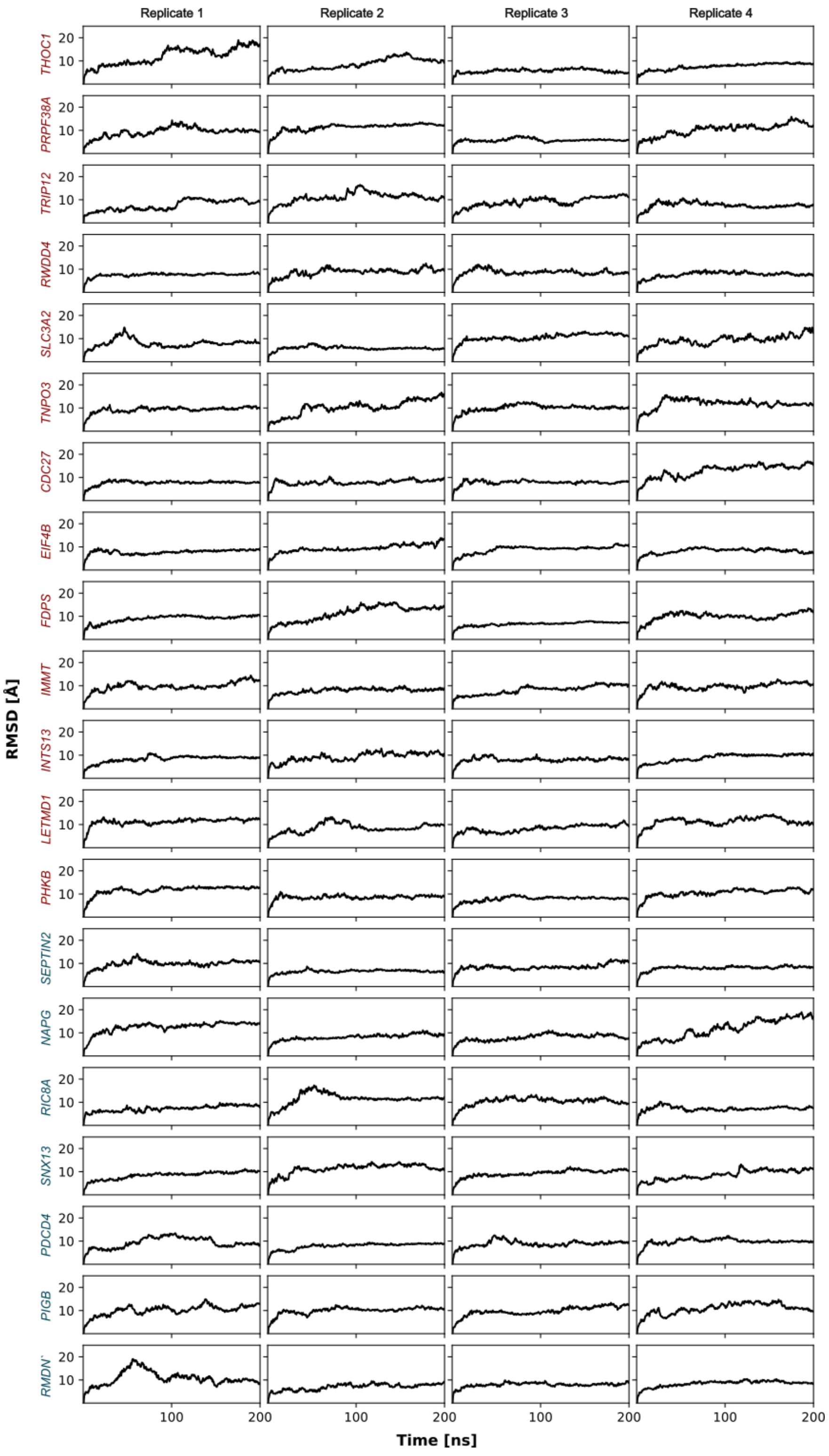


**Supplementary Figure S20.** Root mean square deviation (RMSD) of the SF3B1 backbone (CA, C, N) for molecular dynamics simulations of the downstream BP bound to SF3B1$^{K700E}$. mRNAs with 3' alternative splice site differentially spliced between SF3B1$^{mut/wt}$ and SF3B1$^{wt/wt}$ mRNAs are shown in red and non-differentially spliced are shown in blue.

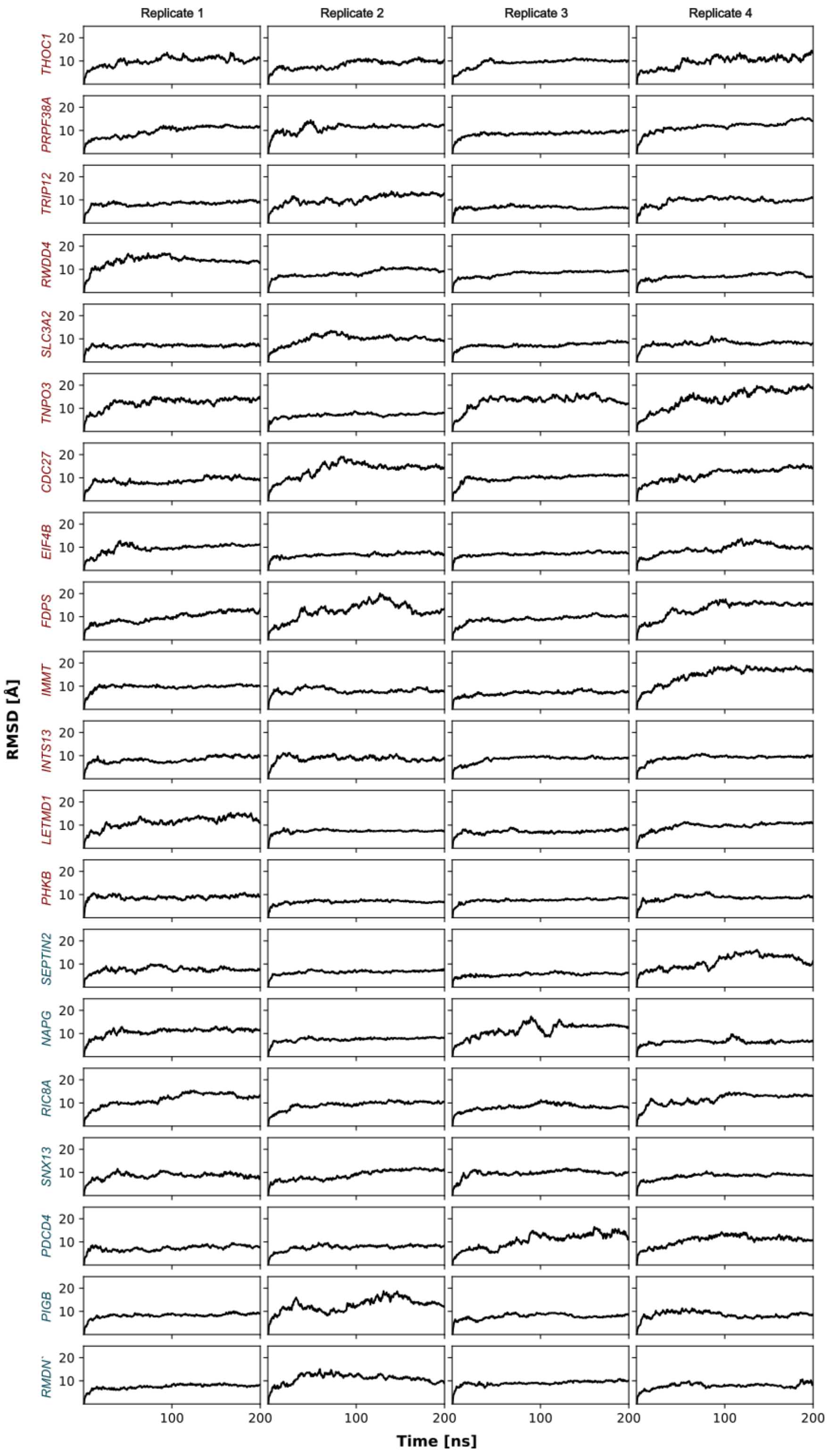


**Supplementary Figure S21.** Root mean square deviation (RMSD) of the SF3B1 backbone (CA, C, N) for molecular dynamics simulations of the alternative upstream BP (BP') bound to SF3B1$^{wt}$. mRNAs with 3' alternative splice site differentially spliced between SF3B1$^{mut/wt}$ and SF3B1$^{wt/wt}$ mRNAs are shown in red and non-differentially spliced are shown in blue.

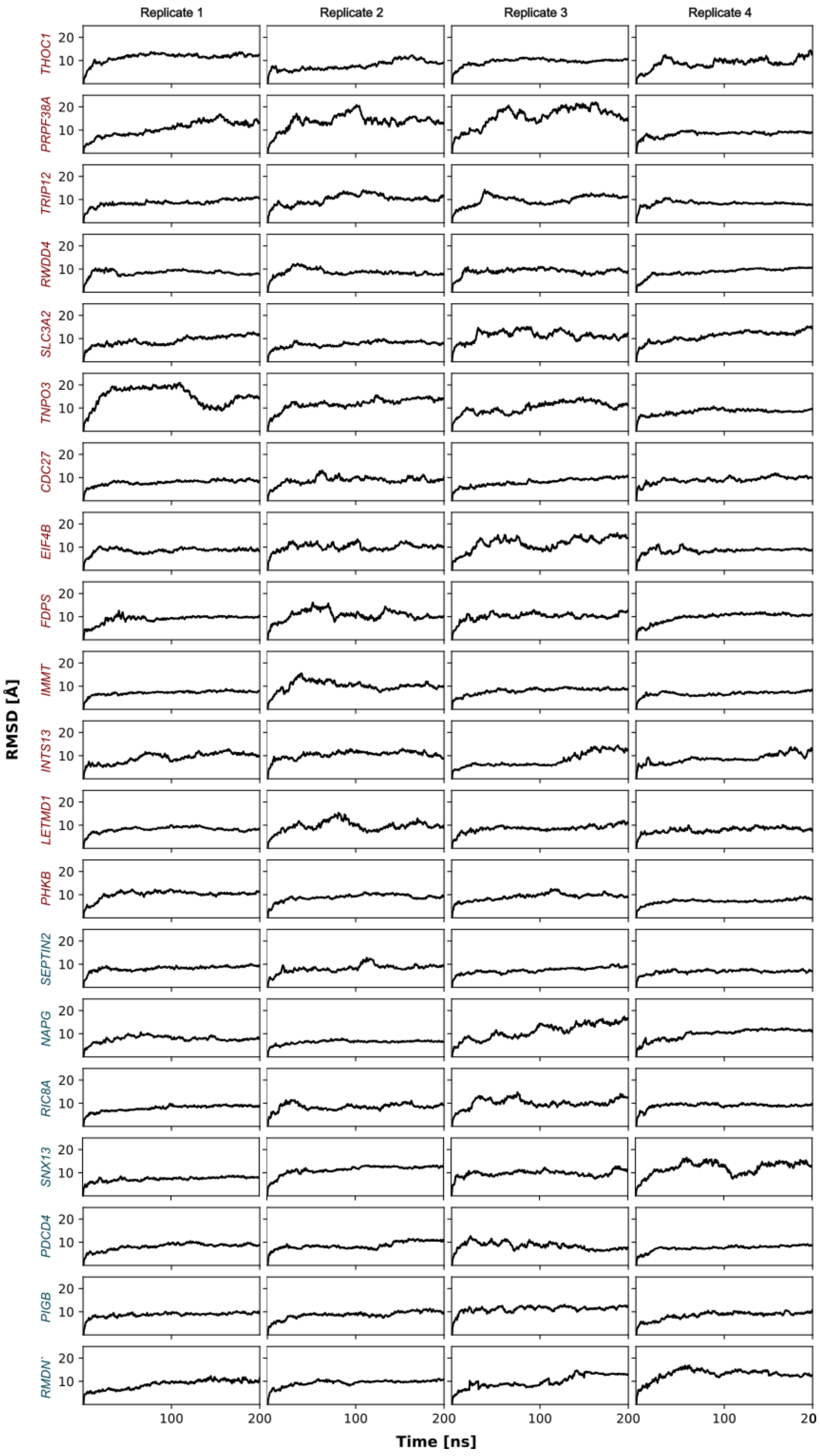


**Supplementary Figure S22.** Root mean square deviation (RMSD) of the SF3B1 backbone (CA, C, N) for molecular dynamics simulations of the alternative upstream BP (BP') bound to SF3B1$^{K700E}$. mRNAs with 3' alternative splice site differentially spliced between SF3B1$^{mut/wt}$ and SF3B1$^{wt/wt}$ mRNAs are shown in red and non-differentially spliced are shown in blue.

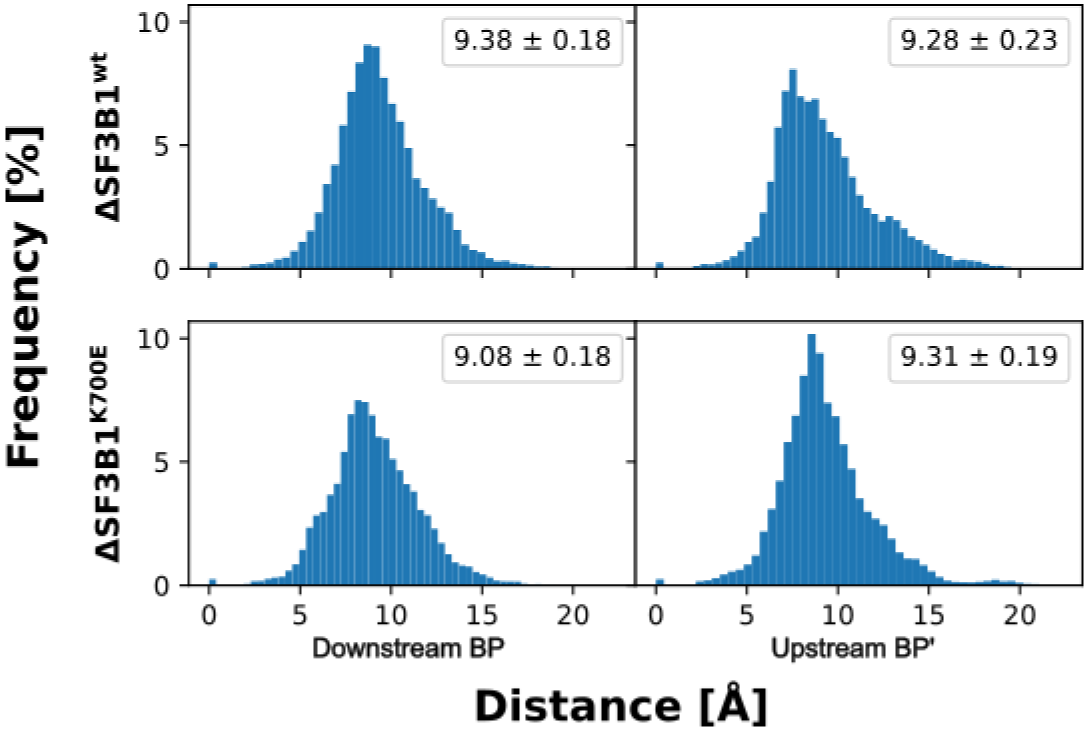


**Supplementary Figure S23.** Distribution of root mean square deviation (RMSD) values of the SF3B1 backbone (CA, C, N) for molecular dynamics simulations of the downstream branch point (BP) bound to SF3B1$^{wt}$ (top left), downstream BP bound to SF3B1$^{K700E}$ (bottom left), alternative upstream BP (BP') bound to SF3B1$^{wt}$ (top right), and alternative upstream BP (BP') to SF3B1$^{K700E}$ (bottom right). Values in the boxes indicate the mean ± SEM.

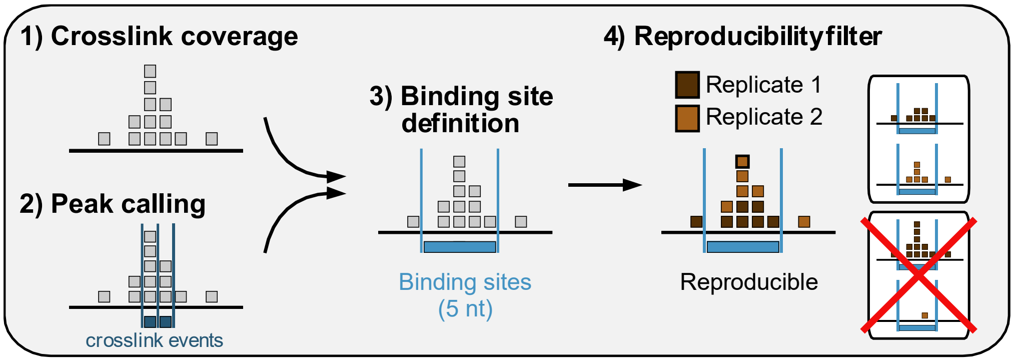


**Supplementary Figure S24.** Schematic workflow of processing iCLIP reads and calling SF3B1 binding sites.

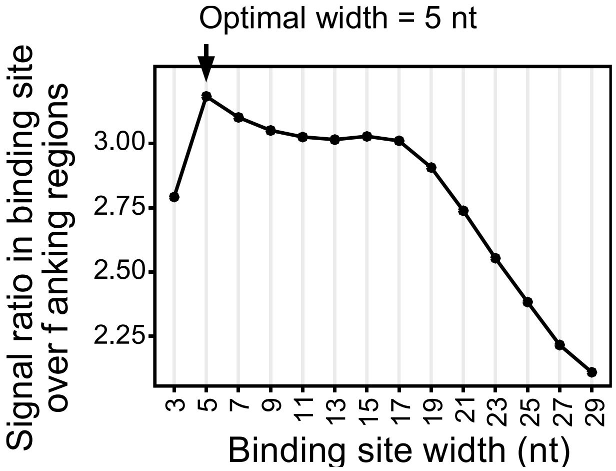


**Supplementary Figure S25.** Defining optimal site binding width. A binding site width of 5 nt optimally captures the SF3B1 crosslink events. Dot plot shows average ratio of crosslink events within binding sites of increasing widths (x-axis) over the mean background signal in flanking windows of the same size, indicating how much more signal occurs within the binding sites compared to their immediate surrounding.

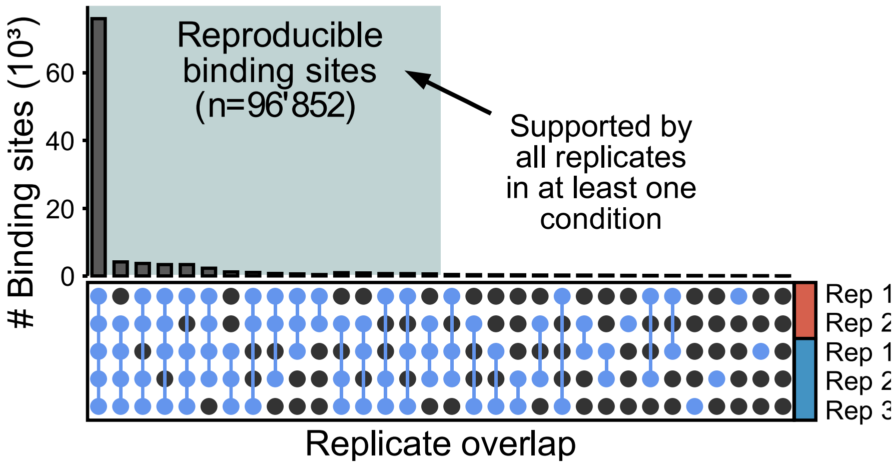


**Supplementary Figure S26.** SF3B1 binding sites reproducibility across replicates. Upper panel shows overlaps of supported binding sites in the replicates, with threshold for sufficient coverage individually adjusted to the signal depth in each replicate[3].

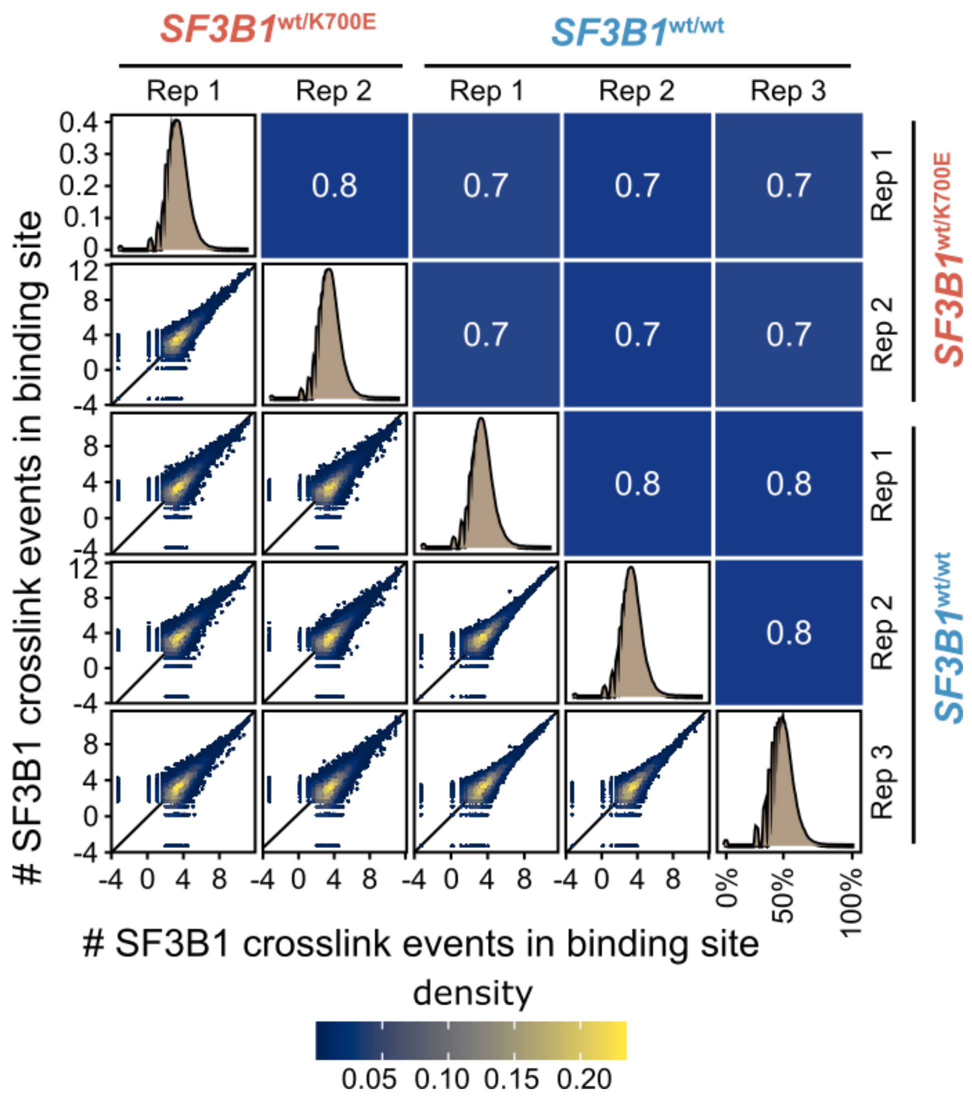


**Supplementary Figure S27.** The SF3B1 iCLIP signal within the binding sites was highly reproducible between replicates. Scatter plot shows the correlation of SF3B1 crosslink events per binding site between biological replicates, with Pearson correlation coefficients given in the opposing quadrant. Distribution of crosslink events per binding sites in each replicate are shown along the diagonal.

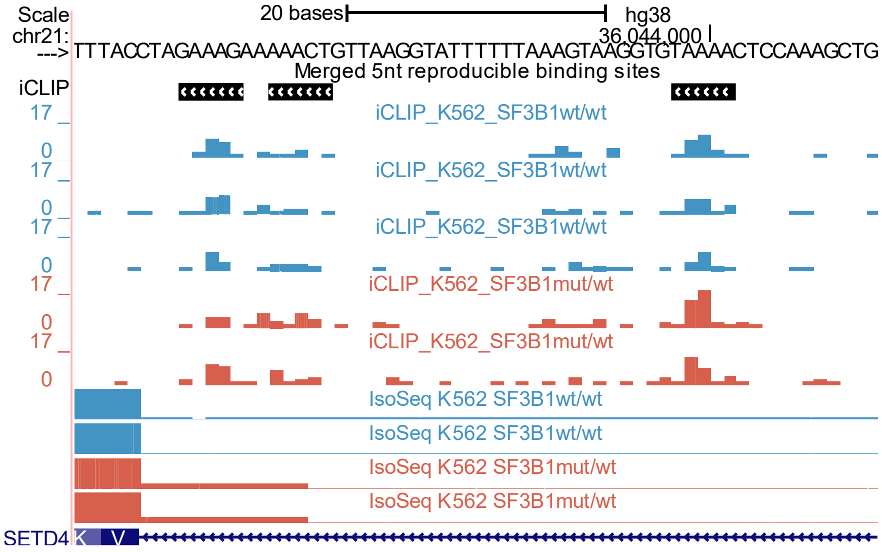


**Supplementary Figure S28.** iCLIP signal within C3 DoubleWide region at *SETD4* exon/intron junction. The UCSC browser tracks show the iCLIP minus strand coverage (top) and IsoSeq read coverage (bottom) in K562-SF3B1$^{K700E/wt}$ (red) and K562-SF3B1$^{K700E/wt}$ cells (blue).

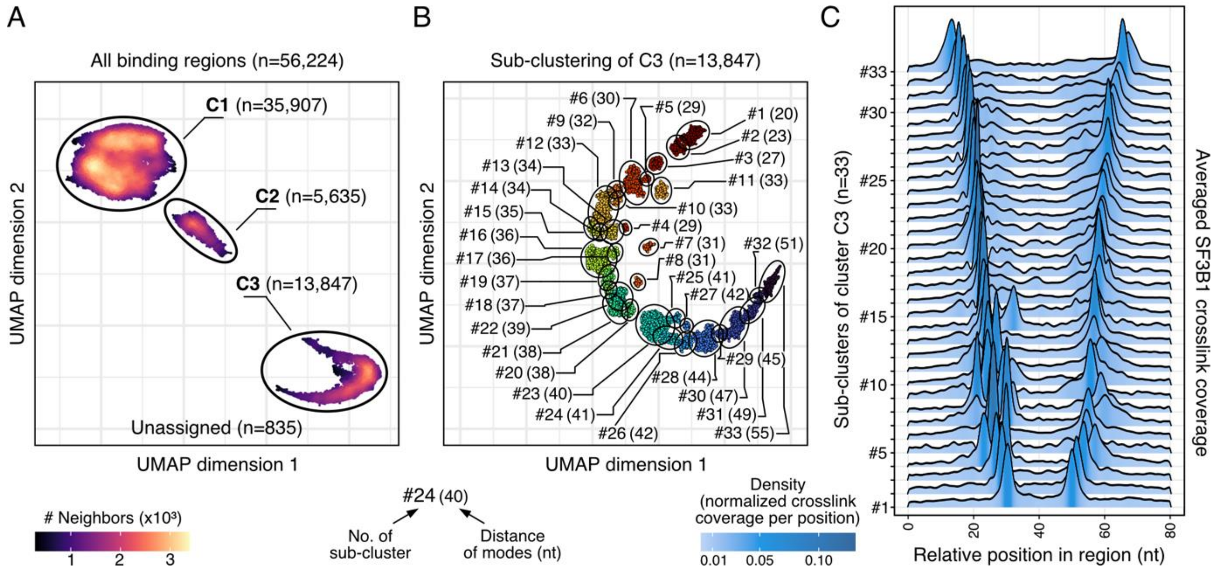


**Supplementary Figure S29.** SF3B1-bound regions show distinct binding patterns. **A** – Unsupervised clustering separates SF3B1-bound regions into three distinct clusters (C1, n = 35,907 regions; C2, n = 5,635; C3, n = 13,847). The SF3B1 crosslink coverage (sum of all replicates) in 81-nt windows around SF3B1 binding sites was subjected to min-max normalization and spline-smoothing, followed by dimension reduction using uniform manifold approximation and projection (UMAP) and density-based clustering of applications with noise (DBSCAN). Regions in cluster C0 (n = 835) could not be assigned to any of the fitted density centres and were excluded from further analysis. **B,C** – Regions of cluster C3 show two broadly spaced modes at variable distance. **B** – UMAP orders regions by increasing distances. Smoothing of the regions in cluster C3 was repeated with higher resolution, followed by a second round of UMAP and DBSCAN, yielding 33 clusters. Clusters were numbered based on the increasing distance between the two modes (indicated in brackets). **C** – Ridge plot displays smoothed SF3B1 crosslink coverages, averaged over the regions within each cluster.

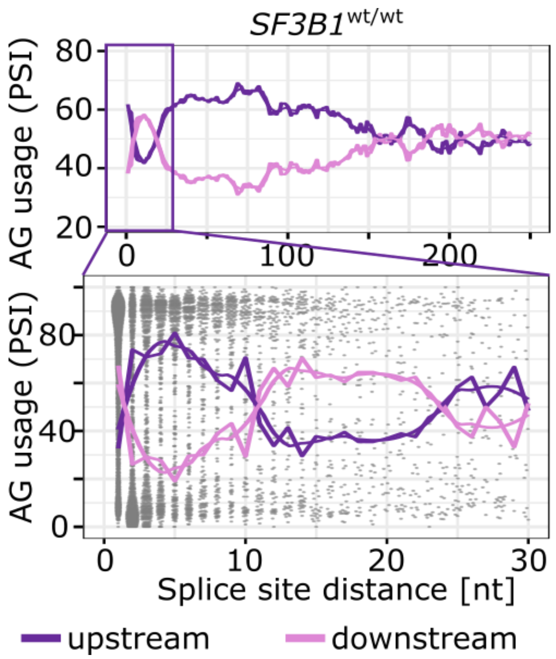


**Supplementary Figure S30.** SF3B1 choice of AG within a narrow window of 12-21 nt is strongly affected by the alternative splice site distance. The AG usage is shown as percent spliced index (PSI) as a function of the splice site distance in SF3B1wt/wt. Rolling mean across 20nt and smoothed (loess method) trend line are shown for upstream (violet) and downstream pink) AG.

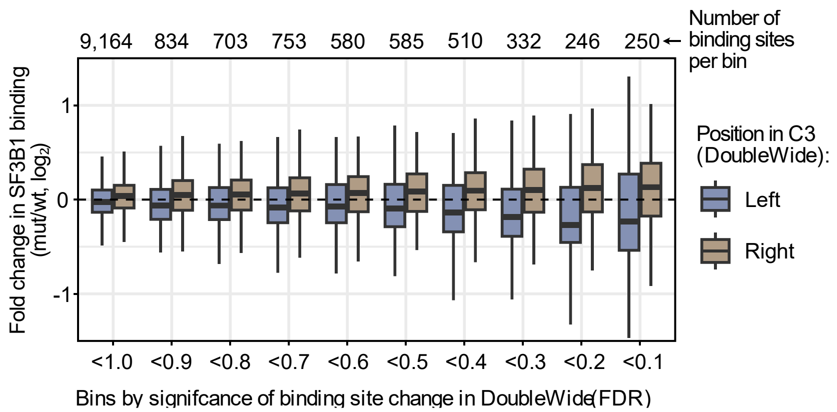


**Supplementary Figure S31.** SF3B1 binding differences between K562-SF3B1$^{K700E/wt}$ and K562-SF3B1$^{wt/wt}$ among cluster C3 DoubleWide regions.

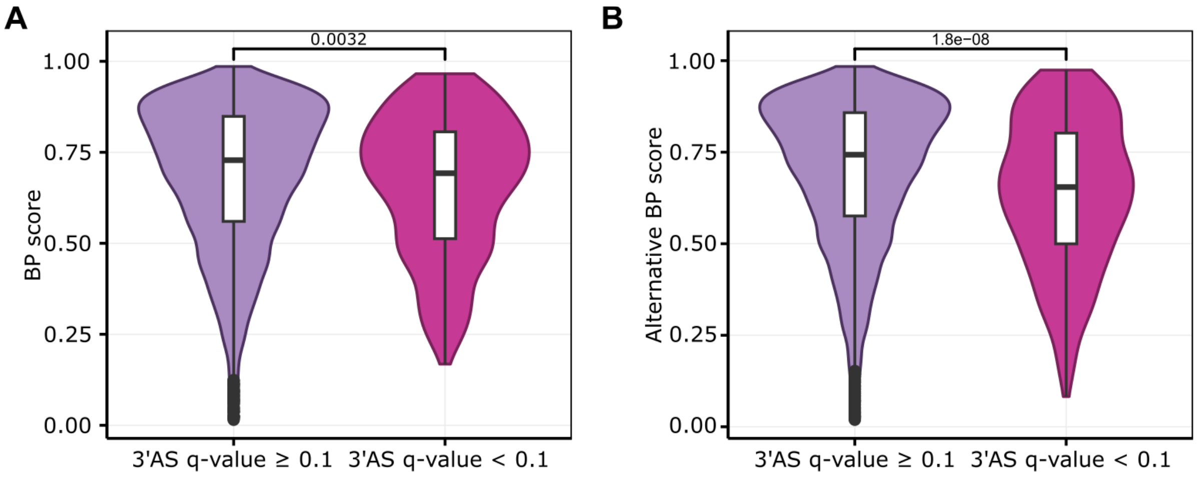


**Supplementary Figure S32.** Branch points (BP) predicted for canonical (panel A) and alternative (panel B) AGs among significant and non-significant 3’ alternative splicing events detected in *SF3B1*$^{mut/wt}$ vs *SF3B1*$^{wt/wt}$ samples based on IsoSeq data.

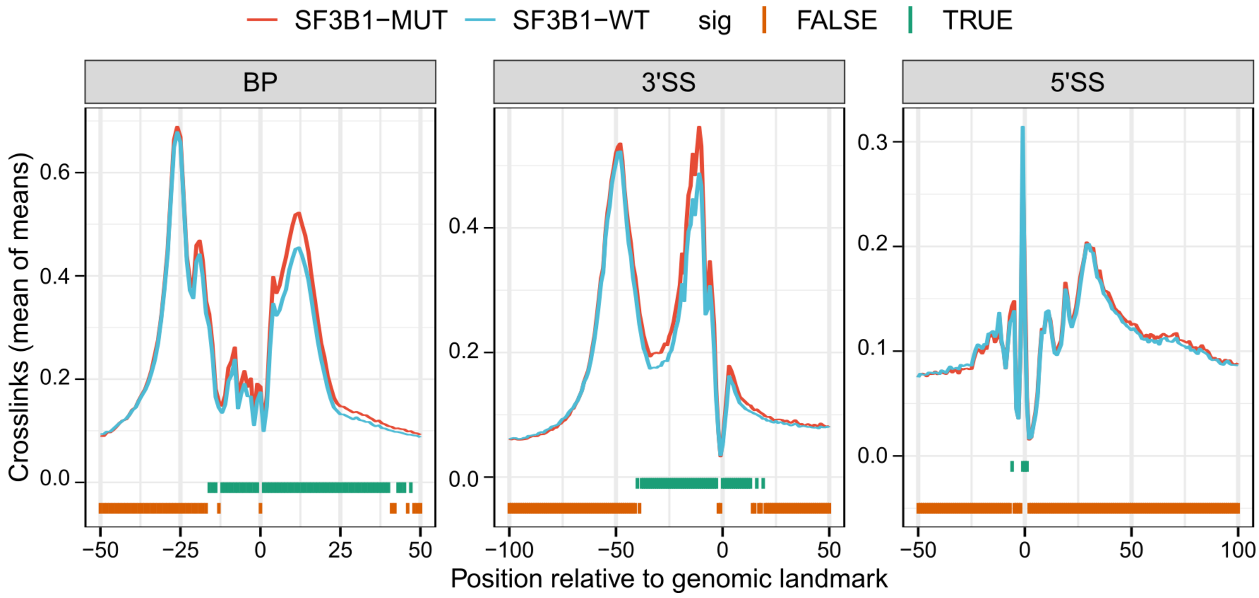


**Supplementary Figure S33.** SF3B1 binding meta-profiles based on the iCLIP experiment with K562-SF3B1$^{K700E/wt}$ and parental K562-SF3B1$^{wt/wt}$ cells. The signal was aligned to branch point (BP), 3' splice site (3'SS) or 5' splice site (5'SS).

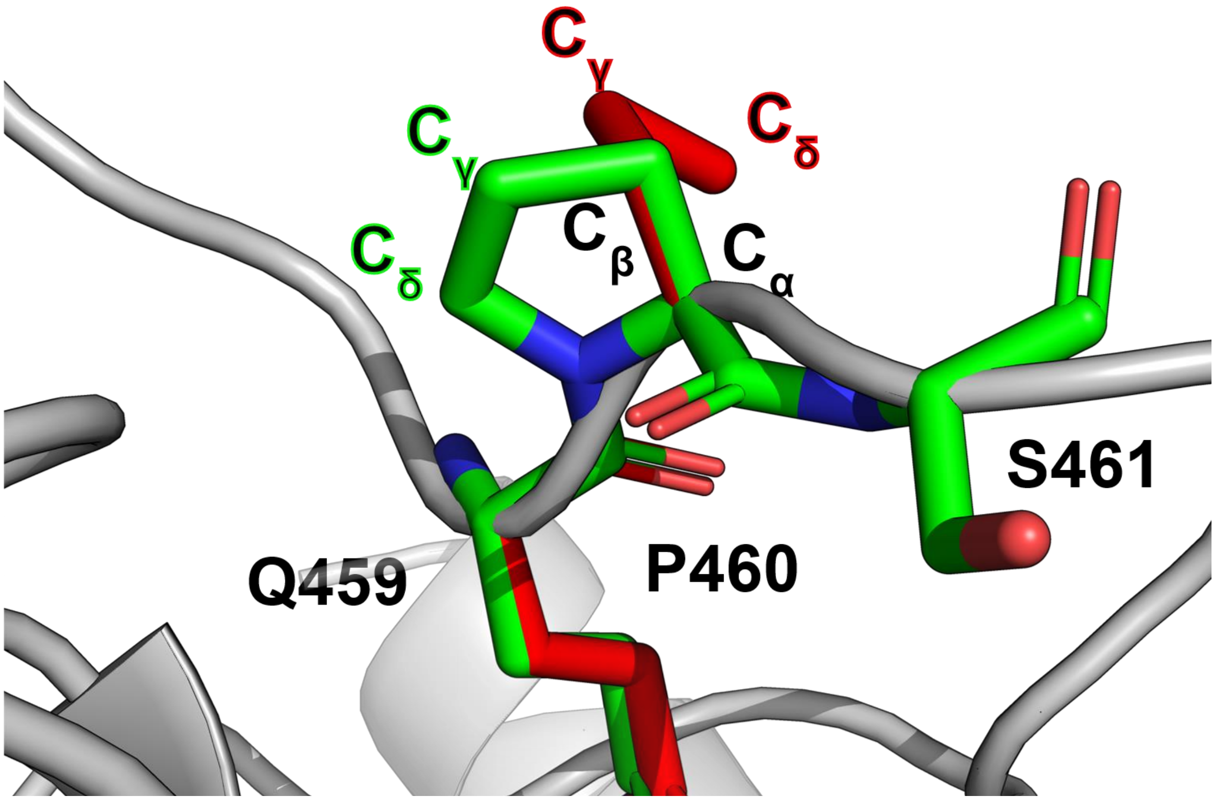


**Supplementary Figure S34.** Incorrect proline conformation after side chain filling using Maestro[4] (red) with a missing bond between $C_\delta$ and N. After further optimization using MOE[5] (green) $C_\gamma$ is correctly bound to N. P460 of SF3B1 is shown exemplarily.

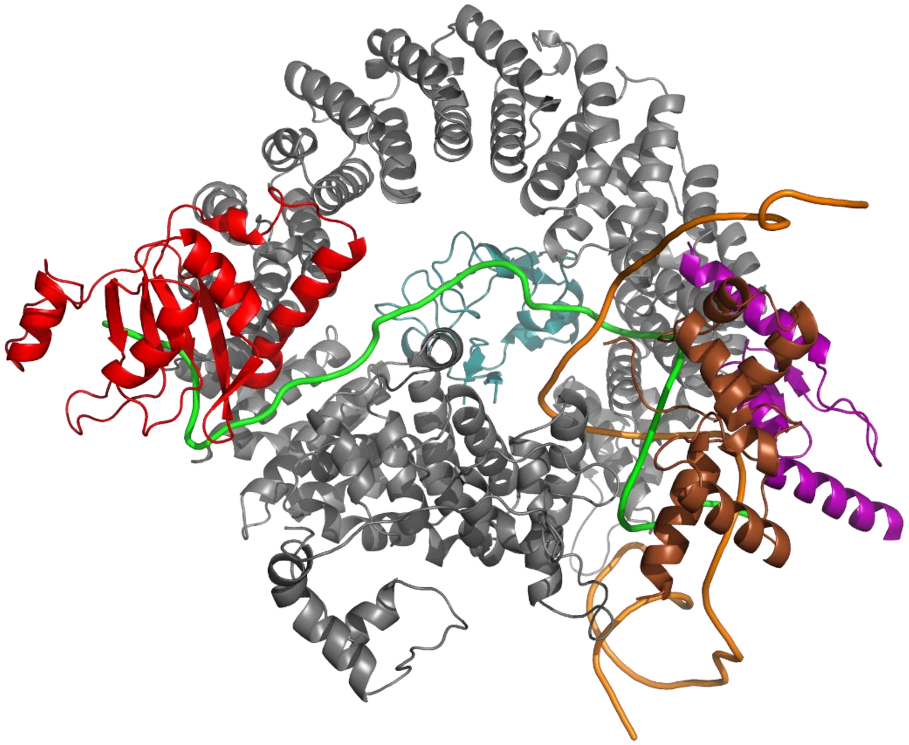

**Supplementary Figure S35.** Final model used for molecular dynamics simulations featuring SF3B1 (grey), the pre-mRNA (green), the PHD finger-like domain containing protein 5A (light blue), the RNA-binding motif protein, x-linked 2 (red), the cell division cycle 5-like protein (brown), the splicing factor 3A subunit 2 (purple), and the u2-snRNA (orange).
